# Spectral envelopes of facial movements predict intention, cortical representations, and neural prosthetic control

**DOI:** 10.1101/2025.09.10.675423

**Authors:** Richard Hakim, Akshay Jaggi, Gyuryang Heo, Hideyuki Matsumoto, Naoshige Uchida, Mitsuko Watabe-Uchida, Sandeep Robert Datta, Simon Musall, Bernardo L. Sabatini

## Abstract

Animals, including humans, use coordinated facial movements to sample the environment, ingest nutrients, and communicate. Rodents, in particular, produce rhythmic facial movements during spontaneous behavior and cognitive tasks. Measuring these movements precisely and linking them to neural activity remains challenging. We introduce face-rhythm, an unsupervised pipeline that combines markerless point-tracking, spectral analysis, and non-negative tensor component analysis to decompose facial video into a small set of interpretable components. Applied to videos of mice during a Pavlovian odor-reward task, a brain-machine interface (BMI) task, and free behavior, face-rhythm recovers human-interpretable behaviors such as whisking, sniffing, licking, and subtler motifs. The resulting components are consistent across animals, are sufficient to decode task variables or internal belief states, and explain cortical activity using a low-rank representation. We also find that the activity of neurons in face-associated primary motor cortex (M1) is predicted well by a phase-invariant spectral transformation of facial movements above ∼0.5 Hz, while slower movements retain a phase-variant representation better predicted by the instantaneous position of the face; individual neurons can show either or both forms of tuning. A systematic comparison against deeplearning point-tracking models, contrastive-learning embeddings, and vision-transformer features places face-rhythm competitively across tasks while also achieving the goal of producing a low-dimensional, interpretable description of rodent facial behavior that is closely linked to cortical activity.

## 2 Introduction

A central question in motor neuroscience is how motor cortical activity is transformed into behavior. The face is a uniquely useful system for investigating this question: sensory organs and dozens of muscles are densely clustered in a compact area, enabling a rich repertoire of coordinated movements with many ethological roles (Galen 1968). Facial behavior also embodies a high-dimensional projection of internal state (Darwin 1872; Dolensek et al. 2020), representing what an animal has experienced, is experiencing, or will do. Yet despite descriptions of several circuits that produce specific facial behaviors, the principles governing how cortical activity is transformed into coordinated facial movement remain incompletely understood.

Early facial-behavior analysis used manual codification of static expressions and actions (Ekman and Friesen 1976; Bartlett et al. 1999; Calder et al. 2001; Cohn and Ekman 2008); machine learning has since produced more automated and objective methods that extract and decompose facial pose and movement. For nonhuman animals, these fall into a few categories. Supervised pose-tracking algorithms such as DeepLabCut (A. Mathis et al. 2018) and SLEAP (Pereira et al. 2022) perform markerless point-tracking of defined body parts and have democratized behavioral quantification, though, despite ongoing efforts (Ye et al. 2024; Syeda et al. 2024), they often require extensive manual curation across video sources and laboratories. Pose-description methods, including most facial action coding and expression recognition (Li and Deng 2022), label static poses but do not analyze time-varying patterns. One example in rodents uses histograms of oriented gradients to describe emotional states (Dolensek et al. 2020). Behavioral-description methods instead chunk continuous behavior into interpretable ‘syllables’ such as rearing or scurrying, including MoSeq (Wiltschko et al. 2015), B-SOiD (Hsu and Yttri 2021), and VAME (Luxem et al. 2022), among others (Batty et al. 2019; Bohnslav et al. 2021; Mackevicius et al. 2019). However, these methods represent movement as discrete states, which does not capture the continuous, overlapping motor patterns that animals express simultaneously.

Whole-face video can also associate facial movement with neural data. “Motion-energy” decomposition (PCA of pixel-intensity changes) of uninstructed facial movements generates a set of behavior-related signals that explain a large fraction of neural variance (Musall et al. 2019; Stringer et al. 2019; Salkoff et al. 2020), and FaceMap decomposition improves neural prediction using markerless point-tracking and neural networks (Syeda et al. 2024). These approaches are insightful but yield high-dimensional representations that are hard to interpret when regressing neural data. There remains a need for unsupervised models with stronger inductive biases that produce low-dimensional, interpretable components matching latent neural activity.

Here we present face-rhythm (FR), a pipeline that performs dense-mesh point-tracking, spectrally transforms the tracked movements, and decomposes them into a sparse set of interpretable components. It is motivated by the function and connectivity of central pattern generators (CPGs) for facial movement (Cramer and Keller 2006; Kaku et al. 2025; Kleinfeld et al. 2023; Moore et al. 2013) and by the success of tensor factorization in finding interpretable components in single-unit data (Williams et al. 2018) and time-frequency EEG representations (Mørup, Hansen, and Arnfred 2007). Spectral representations are especially natural for CPG-driven behaviors, which produce intrinsically oscillatory output whose amplitude envelope, rather than instantaneous phase, may be modulated by descending cortical commands. FR aims to discover and demix facial behaviors that overlap in space, time, and spectral patterning to reveal latent “rhythms”.

FR is designed to discover short temporal patterns (milliseconds to seconds) that map onto known rodent behaviors (whisking, sniffing, licking, and subtler motifs) or onto performance in cognitive tasks. Applied to facial video from classical-conditioning and brain-machine interface (BMI) tasks, a small set of FR components decodes task variables and performance. Motivated by the specialized circuitry for rhythmic facial behavior, we further hypothesized that FR-identified patterns map onto the natural modes of facial motor cortex activity; indeed, FR temporal factors align closely with the dominant modes of M1 population activity. Building a regression model with an explicit spectral transformation, we find that higher-frequency movements (above ∼0.5 Hz) tend to be represented in cortex as a phase-invariant spectral envelope. We also compare FR against deep-learning point-tracking and behavior-embedding methods. FR thus decomposes spontaneous and operant facial behavior into interpretable movements that are consistent across animals and represented in facial motor cortex.

The face-rhythm pipeline code was refined for efficiency and general use and is publicly available here: https://github.com/RichieHakim/face-rhythm

## 3 Results

### 3.1 Face-rhythm methodology

The face-rhythm (FR) pipeline has three stages: unsupervised markerless tracking of a mesh of keypoints across the face; transformation of the point-tracking signals into spectrograms; and decomposition of the resulting spatial-by-frequency-by-time tensor into components that describe discrete parts of behavior (**Figure 1A**; see **Methods 5.1**). Each output ‘component’ describes a behavior and is defined by three ‘factors’: a spatial factor (which points move), a spectral factor (their dominant frequencies), and a temporal factor (the component’s movement magnitude over time) (**Figure 1B-C**).

**Figure 1.**
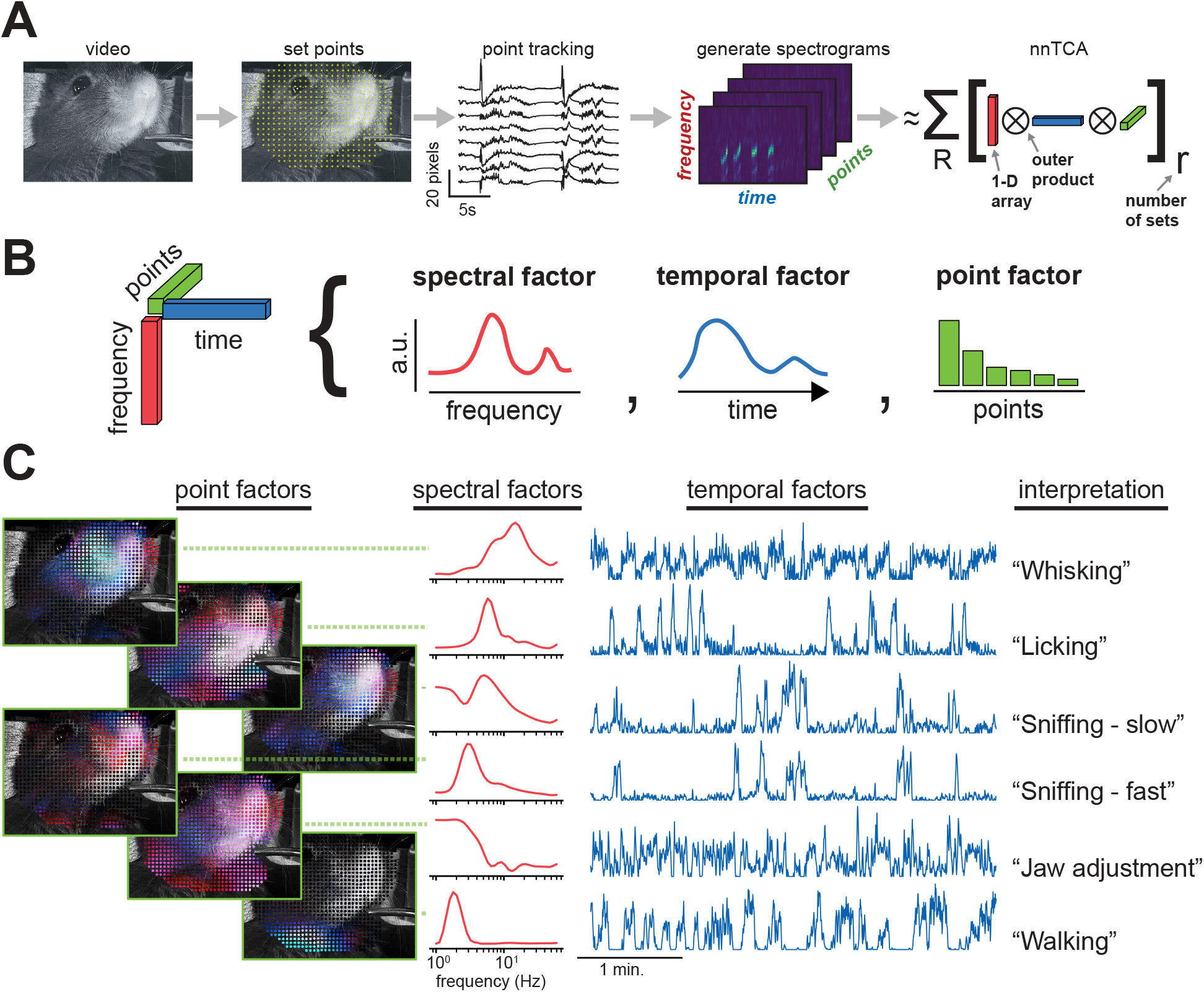
Face-rhythm pipeline. Overview of the face-rhythm pipeline and its output components. A) Pipeline schematic. Face video is the input; point-tracking uses a modified Lucas-Kanade algorithm; per-point spectrograms (‘x’ and ‘y’) are generated with a variable-Q transform; and non-negative PARAFAC tensor decomposition yields the output components. B) Each component comprises three factors: a spectral factor (frequency content of the movement), a temporal factor (when the component is active), and a spatial factor (which points move). The number of components defines the ‘rank’ of the reconstruction. C) Representative face-rhythm components; colors match (A–B). Left: spatial factors (saturation, magnitude; hue, angle, with horizontal ‘x’ blue and vertical ‘y’ red). Middle-left: spectral factors. Middle-right: temporal factors. Right: interpretation of each component.

**Figure 1 supplement 1.**
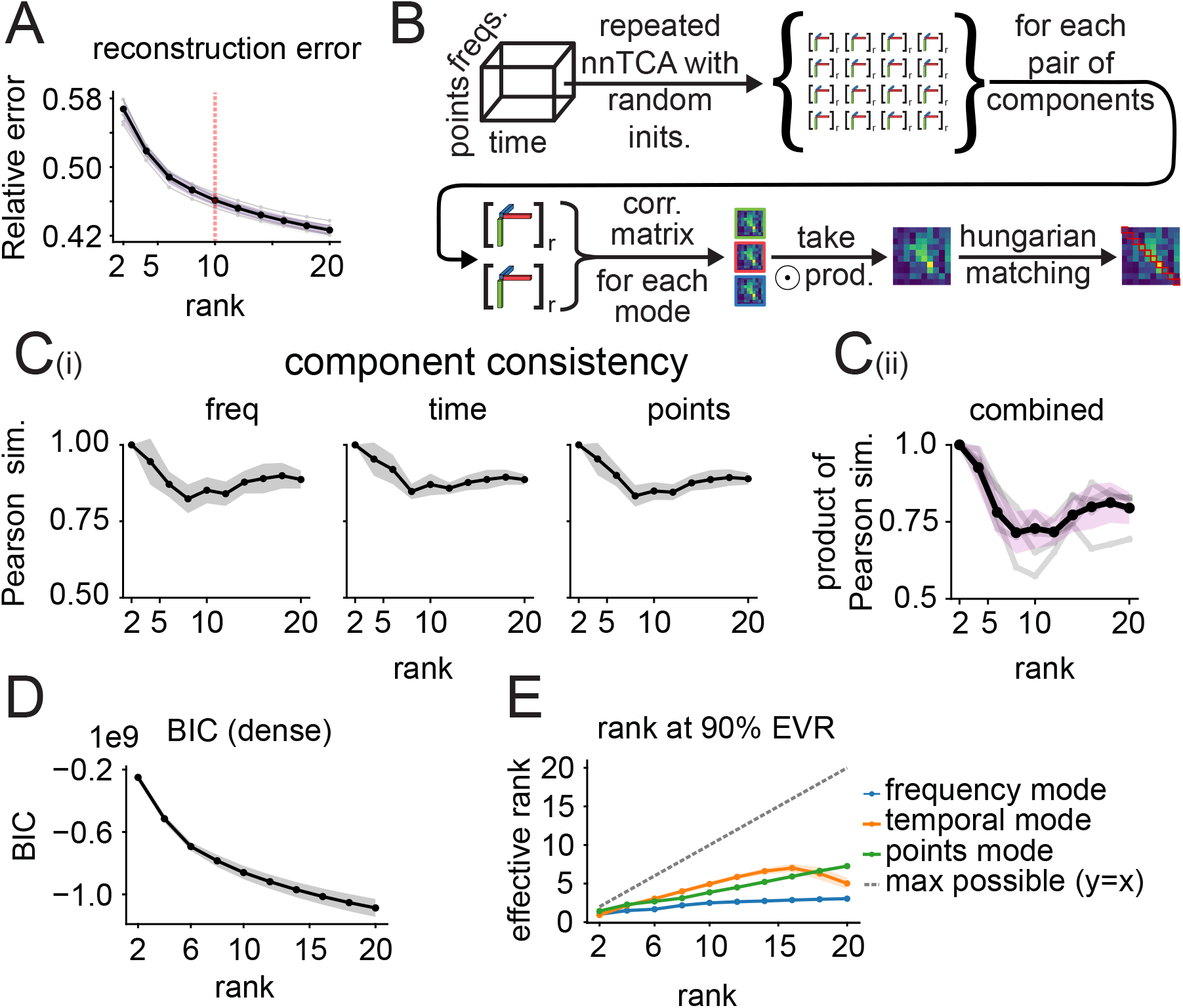
Tensor decomposition rank selection and component consistency. Rank-selection and component-consistency diagnostics for non-negative TCA. A) Relative reconstruction error versus rank for nnTCA of the FR spectrogram tensor; dashed red line, rank used downstream. B) Schematic of the consistency analysis: for each rank, nnTCA is repeated with random starts, mode-wise similarity matrices (frequency, temporal, points) are multiplied, and components are aligned by Hungarian matching. C) Component consistency across rank. Ci) Per-mode Pearson similarity between matched components. Cii) Combined consistency score (product of mode-wise similarities). In (A) and (C), black lines are the session mean and bands ±1 s.d. (*N* = 5 mice, 13 sessions); light traces, sessions. D) Bayesian information criterion versus rank under a dense Gaussian residual approximation. E) Effective dimensionality of each factor matrix (principal components for 90% variance); dashed gray, maximum (*y* = *x*).

**Figure 1 supplement 2.**
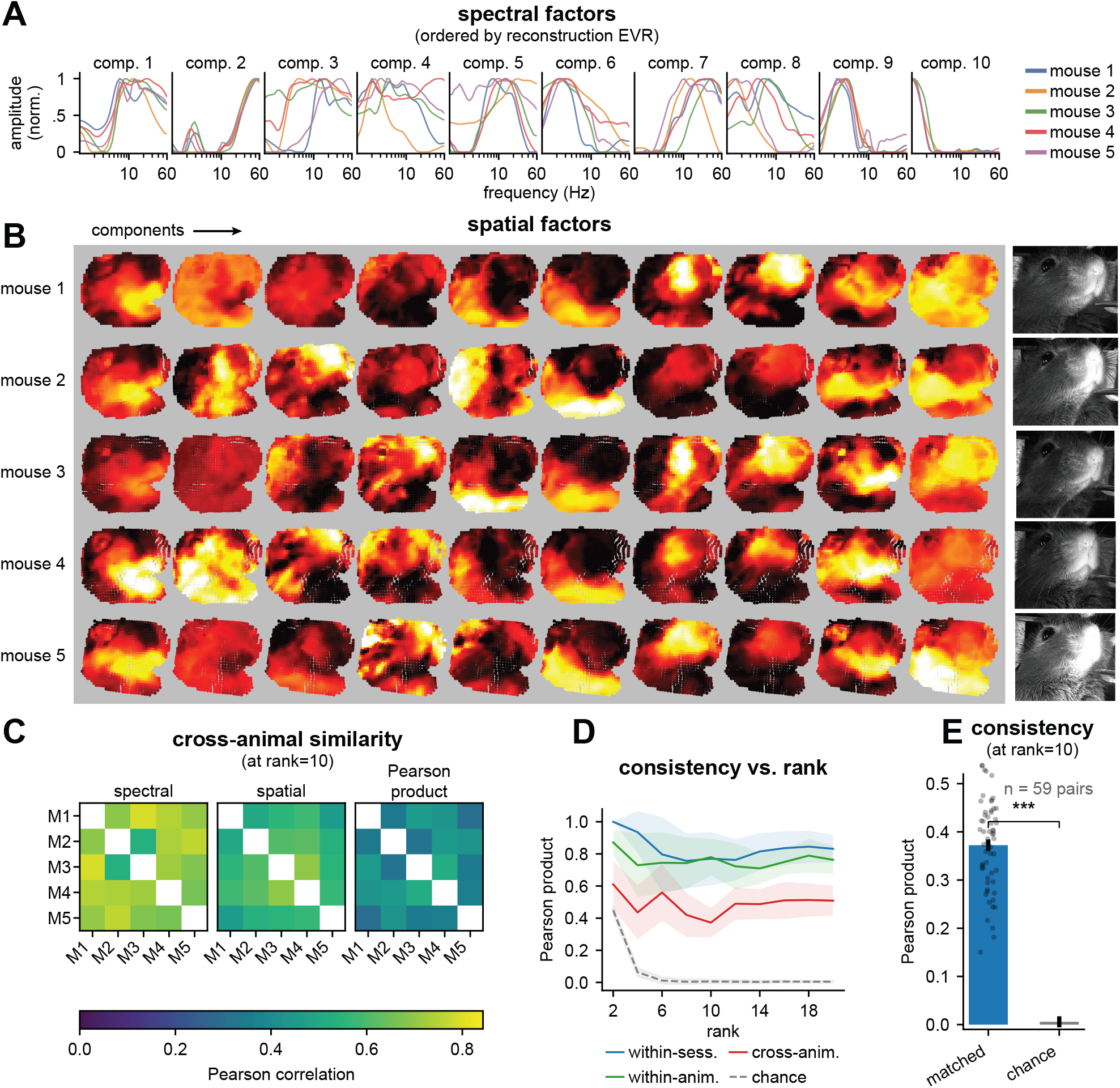
Cross-animal consistency of face-rhythm components. Cross-animal component matching analysis. The pipeline is run on each mouse independently and components are matched across animals with the Hungarian algorithm on the product of spectral- and spatial-factor similarities (see Methods 5.2). A) Matched spectral factors for all 10 components at rank 10 across mice, ordered by average marginal EVR. B) Matched spatial factors (magnitude of *x* and *y*) for the same components (columns matched to A) across mice (rows); reference camera images at right. C) Cross-animal pairwise similarity heatmaps at rank 10: mean spectral correlation (left), spatial correlation (center), and their product (right), averaged over session pairs. D) Pearson product score (spectral × spatial correlation) versus nnTCA rank (*R* ∈ {2, 4, …, 20} ) at three levels: within-session across initializations (blue), within-animal cross-session (green), and cross-animal (red); dashed gray, permutation-null chance; bands, s.d. (13 sessions, 5 mice). E) Consistency statistics at rank 10: mean Pearson product score for matched versus randomly assigned components (error bars, s.e.m. matched, s.d. chance; dots, *n* = 59 cross-animal session pairs). ^∗∗∗^ *p* < 0.001, permutation test.

**Figure 1 supplement 3.**
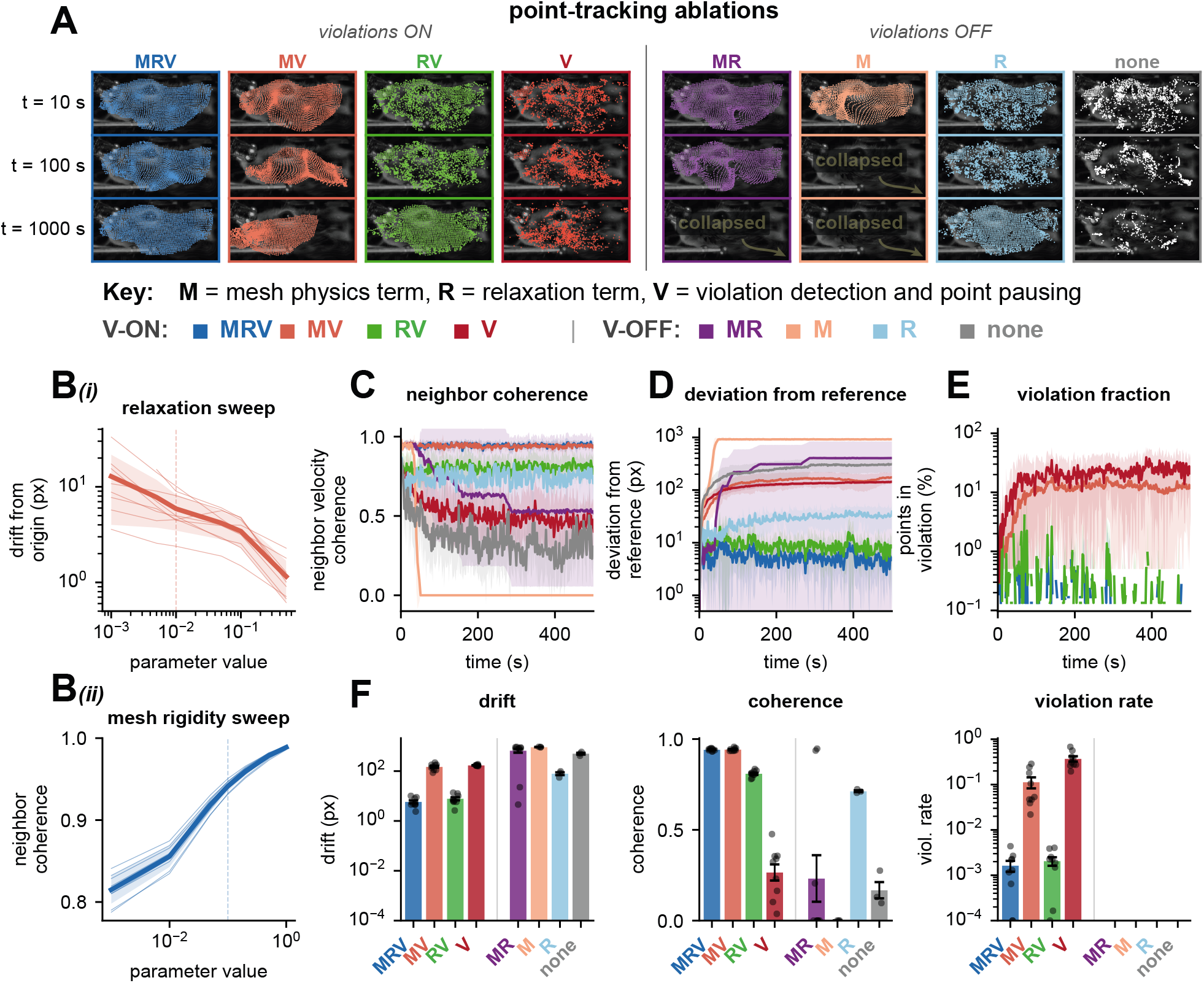
Ablation analysis of Lucas-Kanade point-tracking corrections. Ablation of Lucas-Kanade point-tracking corrections. A) Tracked-point positions at three timepoints (*t* = 10, 100, 1000 s) for eight ablation conditions (columns) of the Lucas-Kanade tracker, where M = mesh physics, *R* = relaxation, V = violation detection with point pausing. Conditions are grouped by violations on (MRV, MV, RV, V) and off (MR, M, R, none), overlaid on a reference frame from a representative widefield session. B) Parameter sensitivity. Bi) Cumulative drift versus relaxation strength (λ_relax_; dashed, final value 0.01). Bii) Neighbor velocity coherence versus mesh rigidity (λ_mesh_; dashed, final value 0.1). Thin lines, sessions; bold, across-session mean; band, ±1 s.d. C) Local neighborhood coherence (mean cosine similarity of frame-to-frame displacement between each point and its 4 nearest neighbors) over the first 500 s. D) Deviation from original point positions over time (log y). E) Fraction of points in violation over time (see Methods 5.3). F) Summary of cumulative drift (left), final neighbor coherence (middle), and violation rate (right). Bars, across-session means; dots, sessions; error bars, s.e.m. *n* = 9 sessions from 7 mice.

The pipeline uses high-performing unsupervised methods that require minimal parameter tuning: a modified Lucas-Kanade algorithm enhanced with mesh physics for tracking (Lucas and Kanade 1981; Baker and Matthews 2004), a custom variable-Q transform (VQT) to compute spectrograms (Brown 1991; Chatterji et al. 2004), and non-negative tensor component analysis (nnTCA, also known as non-negative PARAFAC) for the decomposition (Carroll and Chang 1970; Paatero and Tapper 1994; Harshman 1970; Williams et al. 2018) (see **Methods 5.1**). The number of components *R* is user-defined; for rodent facial behavior we find 4–12 discriminable, human-interpretable components, with rank selected from reconstruction error, component consistency across random initializations, Bayesian information criterion, and interpretability (see **Methods 5.1.5**; diagnostics in **Figure 1 supplement 1**).

Spectral analysis yields non-negative data, which naturally admits a non-negative decomposition constraint; such factorizations tend to extract a ‘parts-based’, interpretable, demixed representation (D. D. Lee and Seung 1999). Applied to mouse faces, FR components correspond to human-recognizable behaviors such as whisking, licking, and multiple forms of sniffing and twitching (**Figure 1C**; **Supplementary Movie 1– 2**), without training neural networks and even for individual mice recorded in single sessions. Unlike PCA, nnTCA imposes no orthogonality constraint, so multiple components can be coactive and contribute to compound behaviors. For example, whisking and sniffing are often expressed together but decompose into separate, coactive components. The full set of components sums to approximate the magnitude spectrograms of the point-tracking signals, with each component tending to align with a statistical mode that is typically interpretable as a discrete behavior.

FR components can also be matched across animals in similar conditions. We matched nnTCA components across 5 mice using Hungarian assignment (Williams et al. 2018) of FR spectral and spatial factors and found cross-animal similarity significantly exceeding chance at all tested ranks (**Figure 1 supplement 2**; *p* < 0.001, permutation test; 13 sessions; see **Methods 5.2, 5.11.2**).

An ablation of the three Lucas-Kanade corrections (mesh physics, relaxation, and displacement-violation detection with point pausing) shows that each targets a distinct error mode (**Figure 1 supplement 3**; see **Methods 5.3**; all *p* = 0.002, one-sided Wilcoxon signed-rank, *n* = 9 sessions from 7 mice). Removing relaxation produces monotonic drift from the initial positions (**Figure 1 supplement 3A, D**), removing mesh physics collapses local neighbor velocity coherence (**Figure 1 supplement 3C, F**), and removing violation detection allows catastrophic deformations during occlusive events such as grooming and inflates the violation rate (**Figure 1 supplement 3E, F**). Only the complete model maintains stable tracking throughout ∼60 min recordings.

### 3.2 Comparison with motion-energy principal component analysis

Given that FR components align with natural facial behaviors, we examined how well FR temporal factors explain neural activity (**Figure 2**). We compared FR against an established approach for relating facial behavior to neural data (Musall et al. 2019) based on motion-energy PCA (ME PCA). The standard FR pipeline tracked and decomposed facial movement (**Figure 2A**) while widefield GCaMP6s imaging recorded cortexwide activity in mice performing a visual discrimination task with stimulus, delay, and response epochs (**Figure 2B, Supplementary Movie 3**). Posterior visual regions respond most to the stimulus, medial regions during the delay, and frontal regions during the response (**Figure 2B**).

**Figure 2.**
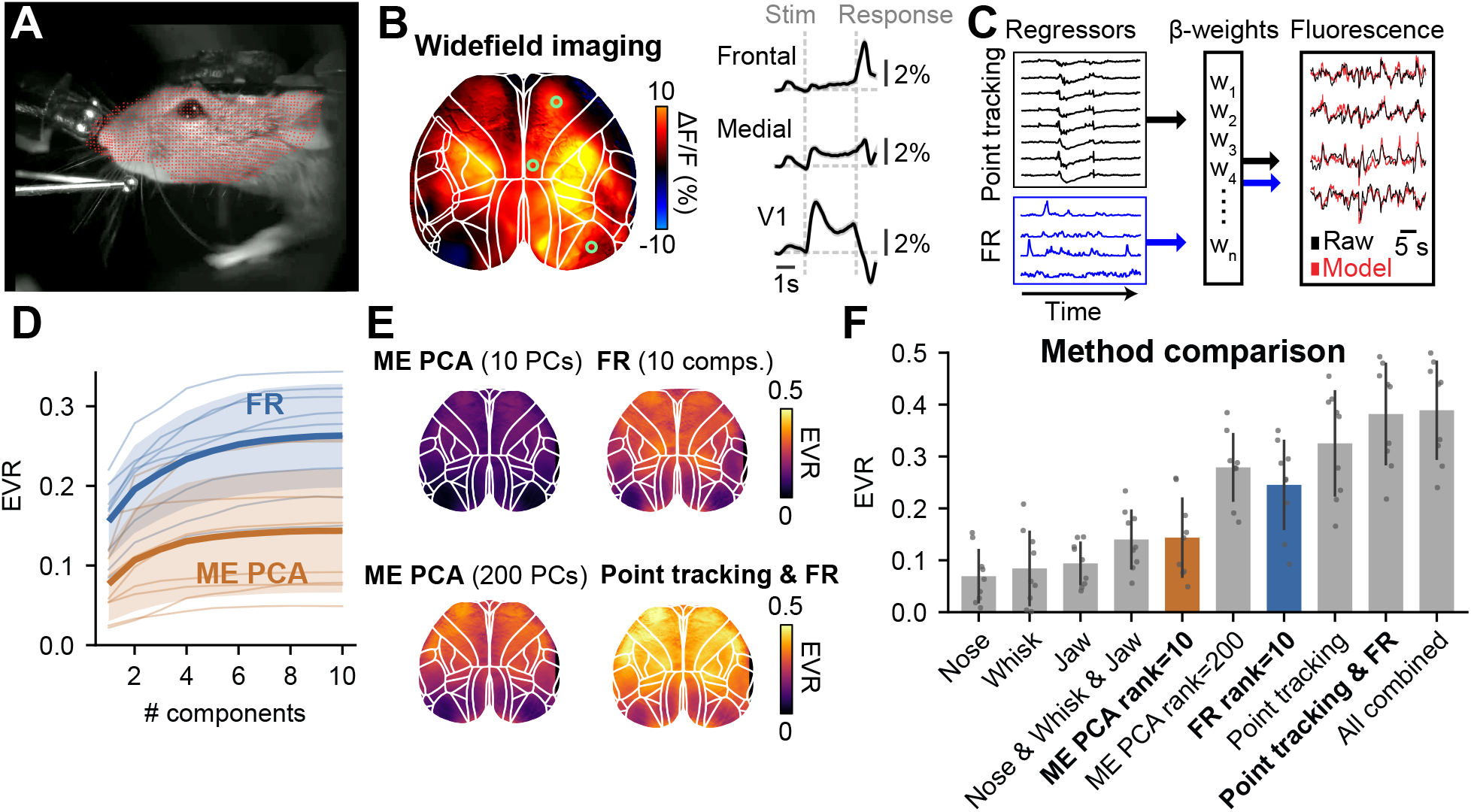
Face-rhythm factors predict cortex-wide widefield neural activity. Prediction of cortex-wide widefield activity from face-rhythm factors. A) Example task-performing mouse on a running wheel with tracked points in red. B) Simultaneous widefield imaging of cortex-wide activity. Left: example frame, with primary visual (V1), medial, and frontal regions marked (light green circles). Right: trial-averaged activity from these regions. C) Schematic of the ridge regression model predicting cortical activity from point-tracking and FR factors. D) Hierarchical regression with increasing numbers of factors, either video motion-energy PCs (ME PCA) or FR temporal factors (FR). Error bars are 95% confidence intervals (*N* = 7 mice, 9 sessions). E) Per-pixel explained variance ratio (mean across mice) for ME PCA, FR, or FR combined with point-tracking. F) Mean EVR across cortex for different regressor combinations; error bars are s.e.m. across mice. *N* = 7 mice throughout.

**Figure 2 supplement 1.**
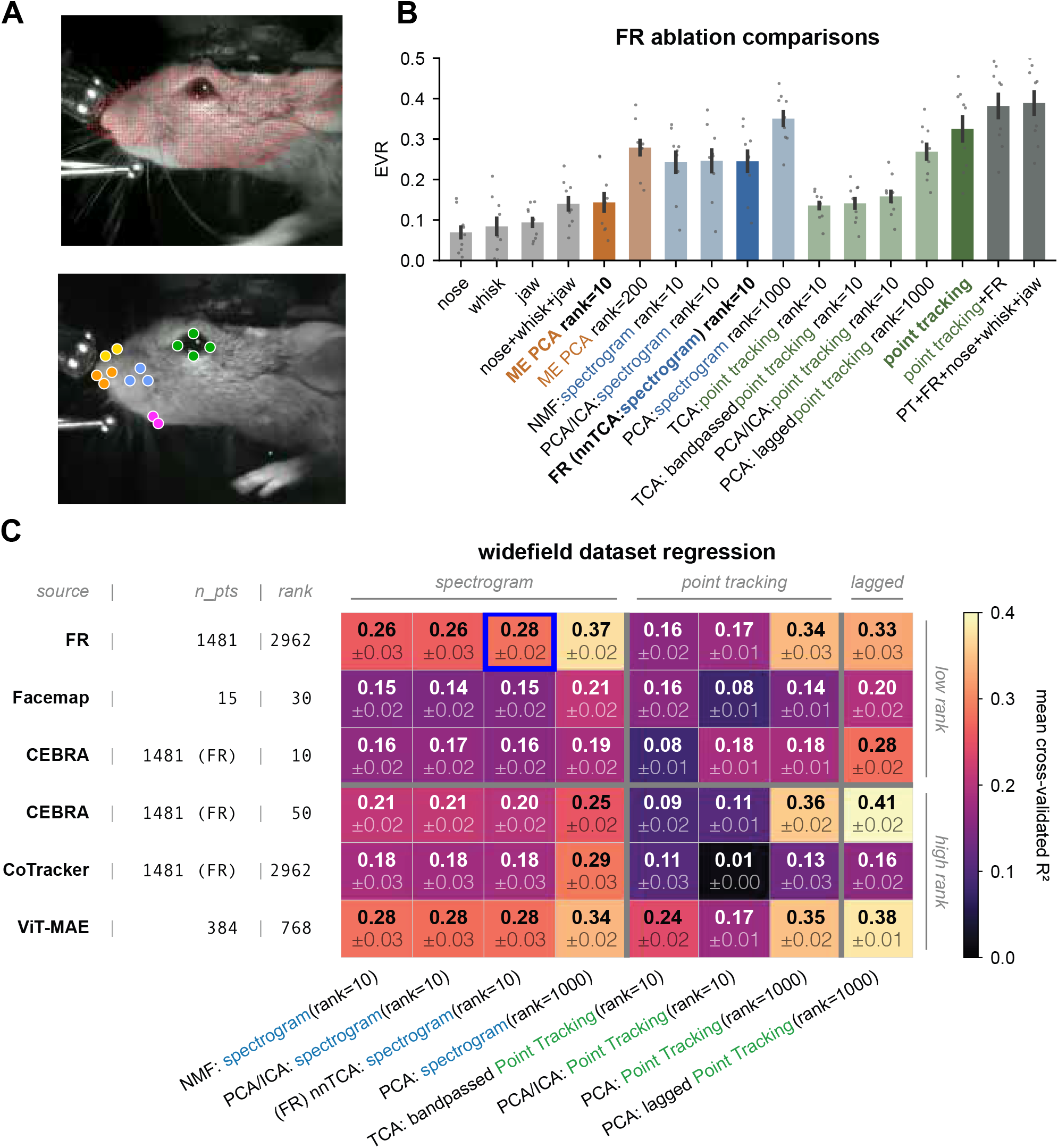
Comparison of keypoint-tracking and decomposition methods for predicting widefield cortical activity. Comparison of trackers and decomposition methods for predicting widefield cortical activity. A) Example frames showing FR dense-mesh point-tracking (top; ∼1,481 points, red) and FaceMap keypoint tracking (bottom; 15 points). B) FR-pipeline comparison of decomposition methods against motion-energy PCA (ME PCA). Bar height is cross-validated EVR of widefield GCaMP6s activity from each regressor. Leftmost bars (nose, whisk, jaw): mean ME within each area; blue bars: decompositions of FR point-tracking + spectrograms; green bars: decompositions of raw FR point-tracking (matrix methods on the flattened 2*P* × *T* displacement matrix; “TCA: point-tracking” on the *P* × 2 × *T* tensor; see Methods 5.4.3); rightmost bars: regressor combinations. Dots, sessions; error bars, s.d. (N = 7 mice, 9 sessions). C) Peer-method comparison of behavioral input representations. Rows: input source (n_pts, number of keypoints if relevant; rank, input feature dimensionality). Columns: decomposition methods grouped by domain (spectrogram, point-tracking, lagged); the displayed rank is the requested upper bound, with achieved rank min(requested, input dimensionality), so low-dimensional inputs are capped at their feature count. Cells show mean cross-validated *R*^2^ ± s.e.m. (*N* = 7 mice, 9 sessions). The top row is the standard FR pipeline; the blue-bordered cell is the standard FR output. Input sources: FR (Lucas-Kanade tracker); FaceMap (Syeda et al. 2024); CEBRA (Schneider, J. H. Lee, and M. W. Mathis 2023) and CoTracker (Karaev et al. 2024) on FR point-tracking; frozen ViT-MAE CLS-token embeddings (768-d) (He et al. 2021; Y. Wang et al. 2025).

**Figure 2 supplement 2.**
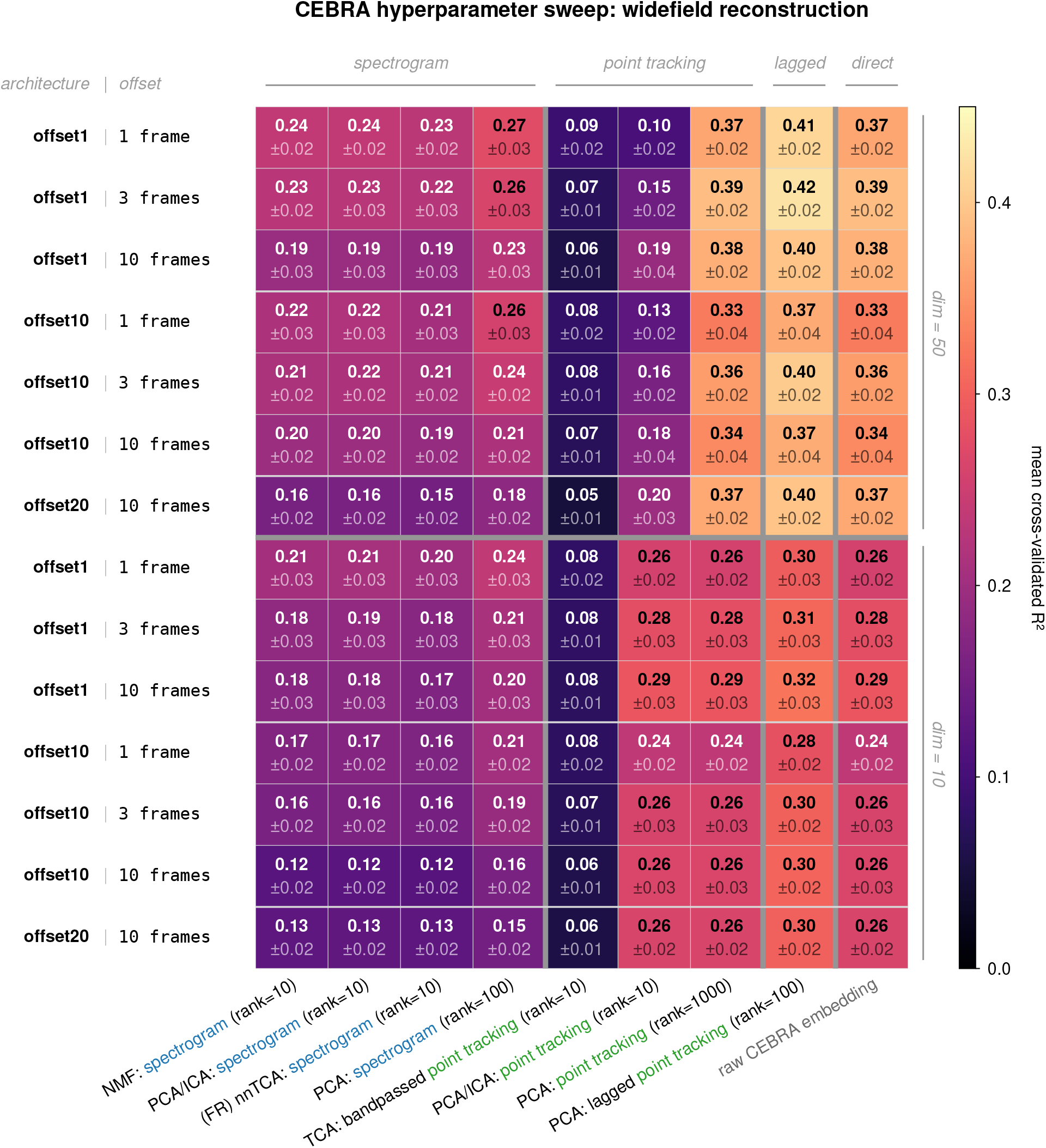
CEBRA architecture and hyperparameter sweep used to select the embedding configuration for the main comparison. Mean cross-validated EVR (equivalently R^2^) for predicting widefield activity from CEBRA embeddings of FR point-tracking, across a sweep of configurations (n = 9 sessions from 7 mice; see Methods 5.4.6). Rows: encoder architecture (<monospace>`offset1`/`offset10`/`offset20`</monospace>) crossed with contrastive time-offset (1, 3, 10 frames), blocked by embedding dimension (50 above, 10 below; `offset20` only at time-offset 10). Columns: the decompositions applied to each embedding plus the raw embedding. Hyperparameters were tuned per cell on held-out data using Optuna (Akiba et al. 2019).

**Figure 2 supplement 3.**
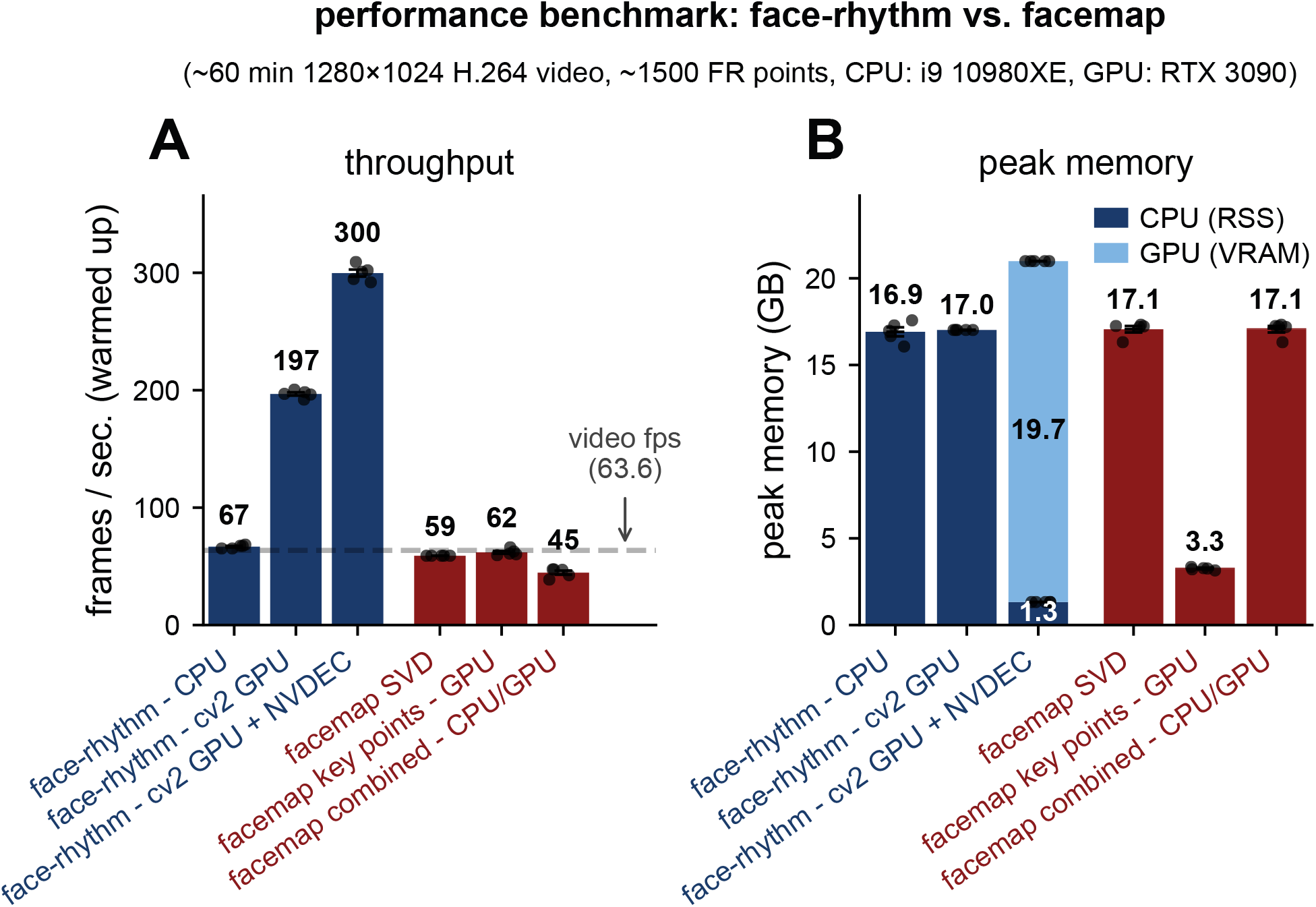
Speed and memory benchmarks. Throughput and memory benchmarks comparing face-rhythm and FaceMap. A) Throughput comparison between face-rhythm (FR; blue) and FaceMap (red) on the same ∼ 60 min, 1280 × 1024 H.264 video and hardware (Intel i9-10980XE CPU, NVIDIA RTX 3090 GPU). Conditions: FR-CPU; FR-cv2 GPU (OpenCV CUDA backend); FR-cv2 GPU+NVDEC (adds NVDEC video decoding); FaceMap-SVD (CPU); FaceMap keypoints-GPU; and FaceMap combined-CPU/GPU. Values annotated above bars; dots, sessions (*n* = 4); error bars, s.e.m. across sessions; dashed line, source video rate (63.56 Hz). Same recording as **Figure 2**. B) Peak memory for the same modes: CPU resident-set size (dark) and GPU VRAM where applicable (light).

**Figure 2 supplement 4.**
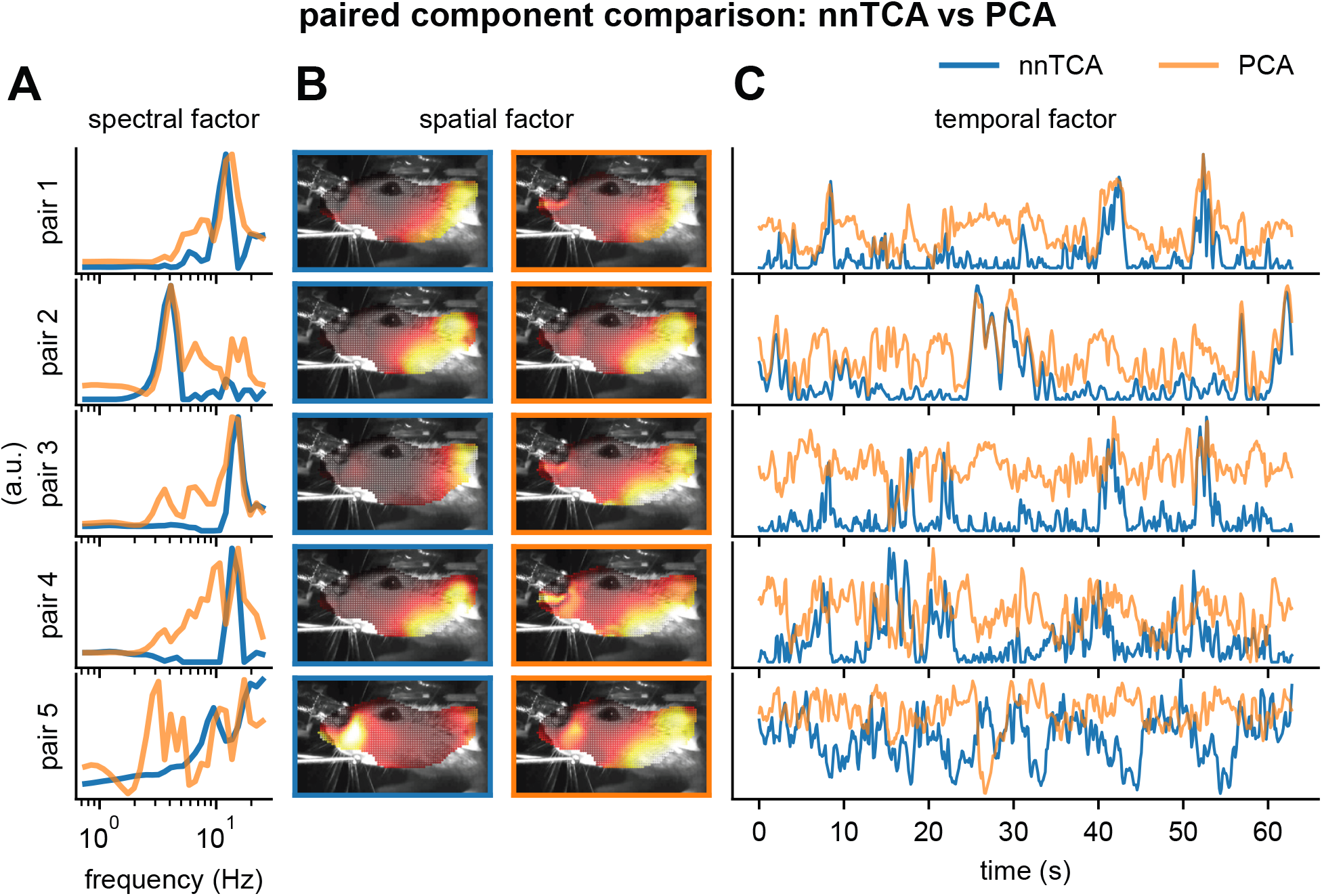
Comparison of non-negative TCA and PCA decompositions of the spectrogram tensor. Paired comparison of non-negative TCA and PCA factors on the spectrogram tensor. A) Spectral factors for five matched component pairs comparing FR nnTCA (blue) and PCA on the flattened spectrogram tensor (orange); rows are pairs matched on temporal factors by the Hungarian algorithm. B) Spatial factors for the same pairs over an example face frame (left, nnTCA; right, PCA; borders colored by method). C) Temporal factors for the same five pairs over time, from one representative session.

We fit a ridge regression model predicting each widefield pixel’s activity (fluorescence transient time series) using behavioral inputs such as FR components or point-tracking (**Figure 2C**). Hierarchical regression shows that FR temporal factors form a low-rank representation significantly more aligned with cortical activity than ME PCA factors (**Figure 2D**). In line with previous findings (Musall et al. 2019), the cross-validated explained variance ratio (EVR) is high across much of cortex, especially anterior regions (**Figure 2E**). In a direct comparison, 10 FR components (0.245 ± 0.087 s.d.) outperform 10 ME PCA components (0.144 ± 0.078; *p* = 0.0039, Wilcoxon signed-rank, *n* = 9 sessions from 7 mice; linear mixed-effects model for repeated sessions, *p* = 4.27 × 10^−8^) and approach the performance of 200 ME PCA components (0.279 ± 0.066), indicating that FR components are more predictive at matched rank.

A more detailed comparison supports FR’s low-rank explanatory power (**Figure 2F**). Ten FR components (0.245 ± 0.087 s.d.) are more predictive than a combination of whisker, nose, and jaw movements (0.143 ± 0.062; *p* = 0.008, Wilcoxon signed-rank, *n* = 9 sessions from 7 mice). Appending the rank-10 FR components to the high-dimensional point-tracking data improves EVR from 0.33 ± 0.10 to 0.38 ± 0.10, suggesting that the cortex maintains a separable representation of the spectral envelope of movement. Appending all signals together changes it only modestly (0.38 to 0.39 ± 0.10), showing that point-tracking and FR signals largely subsume ME PCA and individual body-part signals. Together, these results demonstrate that spectral transformation and tensor factorization yield low-dimensional components with high explanatory power that align with the dominant modes of cortical activity.

We reanalyzed the widefield data after ablating individual pipeline steps, and found that spectrogram generation is a critical transformation (**Figure 2 supplement 1B**; see **Methods 5.4.3**). Among low-rank decompositions (∼10 components), the standard FR pipeline (nnTCA on VQT spectrograms) achieves the highest cross-validated EVR, and spectrogram-domain decompositions consistently outperform point-trackingdomain methods at matched rank.

To test whether FR’s advantage depends on the tracker or on the spectral decomposition, we compared five input sources through the same decompositions (**Figure 2 supplement 1C**): FR, FaceMap (Syeda et al. 2024), CoTracker (Karaev et al. 2024) (a transformer-based dense tracker at FR mesh positions), CEBRA (Schneider, J. H. Lee, and M. W. Mathis 2023) (contrastive embeddings of FR tracking at dimensions 10 and 50; architecture selected by a sweep, **Figure 2 supplement 2**, see **Methods 5.4.6**), and frozen ViT-MAE embeddings (He et al. 2021) (768-d CLS-token features, motivated by BEAST (Y. Wang et al. 2025)).

FR-derived decompositions perform favorably against other keypoint-tracking approaches across nearly all low-rank methods. Frozen ViT-MAE also performs well, but it can draw on extra frame information such as hand and body posture, can require extensive fine-tuning, and yields embeddings that are less directly relatable to fiducial points or muscle dynamics. We conclude that FR point-tracking yields a low-rank representation predictive of widefield activity, and that the spectral transformation and nnTCA steps are critical at low rank.

We benchmarked the computational performance of FR against FaceMap (**Figure 2 supplement 3**; see **Methods 5.15**). FR processes a ∼60 min, 1280 × 1024 video at ∼67 frames/s on a CPU and ∼197–300 frames/s on a GPU, with peak memory ∼17 GB. Runtime scales linearly with video duration and is dominated by point-tracking rather than VQT or nnTCA steps.

Although the nnTCA step’s value is not immediately evident from the regression analysis, a qualitative comparison shows it yields more interpretable components. By matching nnTCA and standard PCA component pairs (**Figure 2 supplement 4**), nnTCA factors appear more population-sparse, decorrelated, demixed, and aligned to behavioral modes (**Figure 2 supplement 4A-C**). For example, a high-frequency whisking factor is localized specifically to the whisker pad.

### 3.3 Conditioning of facial behaviors during reward-guided tasks

We then asked whether FR captures uninstructed movements predictive of conditioned-stimulus identity during Pavlovian/classical conditioning. We analyzed facial video from a previously described reward-probability task (Matsumoto et al. 2016) (**Figure 3**; see **Methods 5.5**). Head-restrained thirsty mice receive one of three conditioned stimuli (CS+, CS0, CS-); after a 1 s odor and 1 s delay they receive water, nothing, or air puff, respectively (**Figure 3A**). FR components show CS-dependent modulation during the cue and delay, including factors consistent with jaw/licking for CS+, respiration for CS0, and ear twitching for CS-(**Figure 3B, Figure 3D-E**). Elastic-net multinomial logistic regression on FR temporal factors (20 sessions, 5 mice) decodes CS identity with high accuracy: 0.92 ± 0.05, 0.87 ± 0.03, and 0.83 ± 0.01 (mean ± s.e.m.) for CS+, CS0, and CS-(all *p* < 0.003 vs. chance of 1/3, one-sample *t-*test, Bonferroni-corrected, *N* = 5 mice; **Figure 3F**).

**Figure 3.**
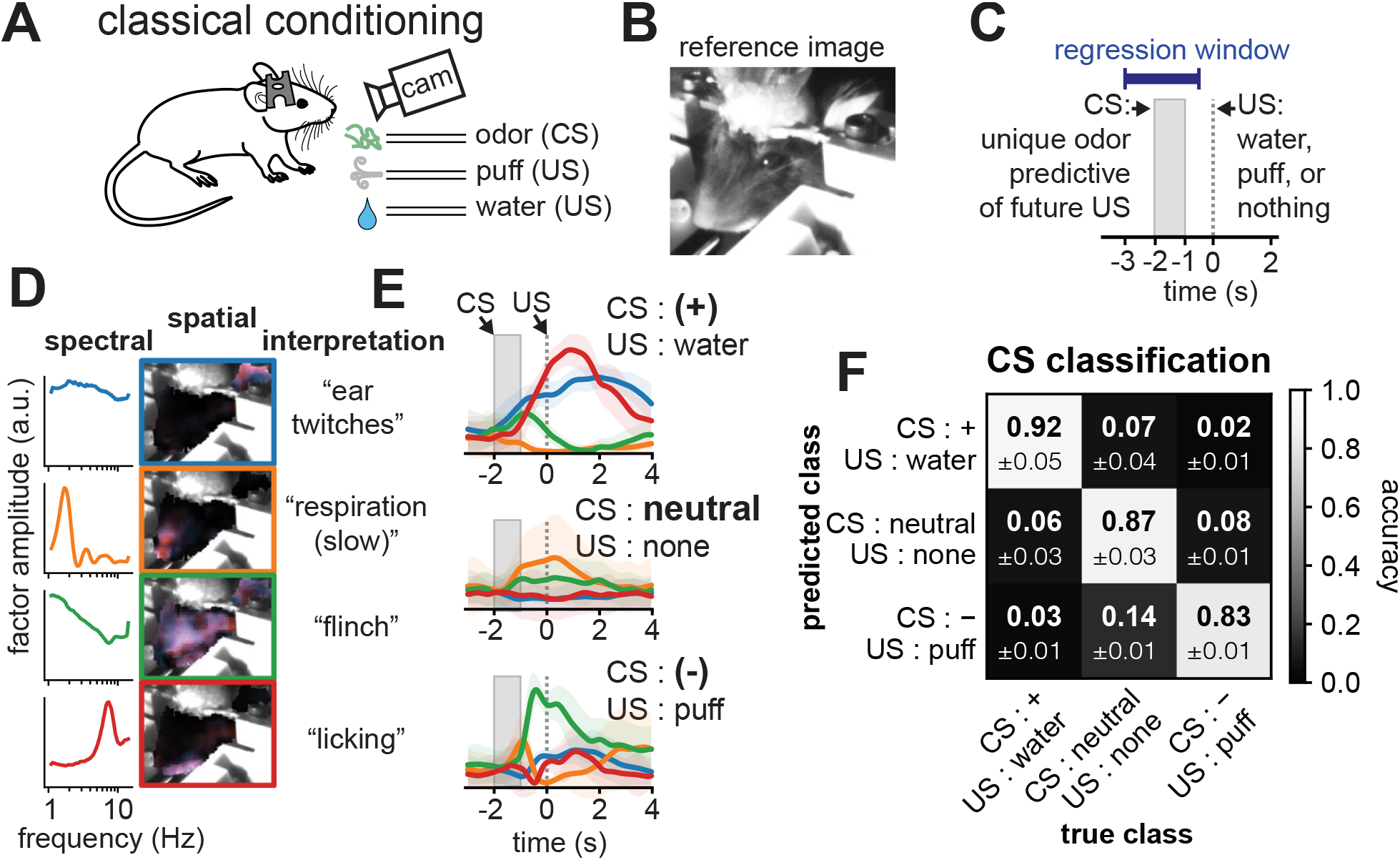
Facial dynamics classify conditioned stimulus identity during classical conditioning. Classification of conditioned stimulus identity from face-rhythm factors during classical conditioning. A) Experiment diagram. Face video was recorded during conditioning with three odors (conditioned stimulus; CS) followed by appetitive (water), neutral (nothing), or aversive (air puff) outcomes. B) Camera field-of-view. C) Classification analysis: FR temporal factors during the CS and delay window predict CS identity via elastic-net multinomial logistic regression (see Methods 5.5.2). D) Representative FR components. Left: spectral factors. Middle: spatial factors (horizontal ‘x’ blue, vertical ‘y’ red). Right: interpretation. E) Representative temporal-factor dynamics around CS presentation; colors match (D), error bars are s.d. across trials. F) Confusion matrix for CS classification (20 sessions, 5 mice); values are mean *P*(predicted | true) ± s.e.m. across animals. Diagonal accuracies exceed the chance level of 1/3 (*p* < 0.003; see Methods 5.5.2).

We also examined uninstructed movements shaped by reward in an operant brain-machine interface (BMI) experiment (**Figure 4**; see **Methods 5.11**), in which an animal was rewarded for increasing neural population activity along a predefined decoder axis (**Figure 4A,C**). Two-photon imaging at 30 Hz was used to record ∼1,000 layer 2/3 jGCaMP8m neurons in facial motor cortex (Mayrhofer et al. 2019; Tamura et al. 2025) while continuous face video is recorded for FR analysis (**Figure 4B**).

**Figure 4.**
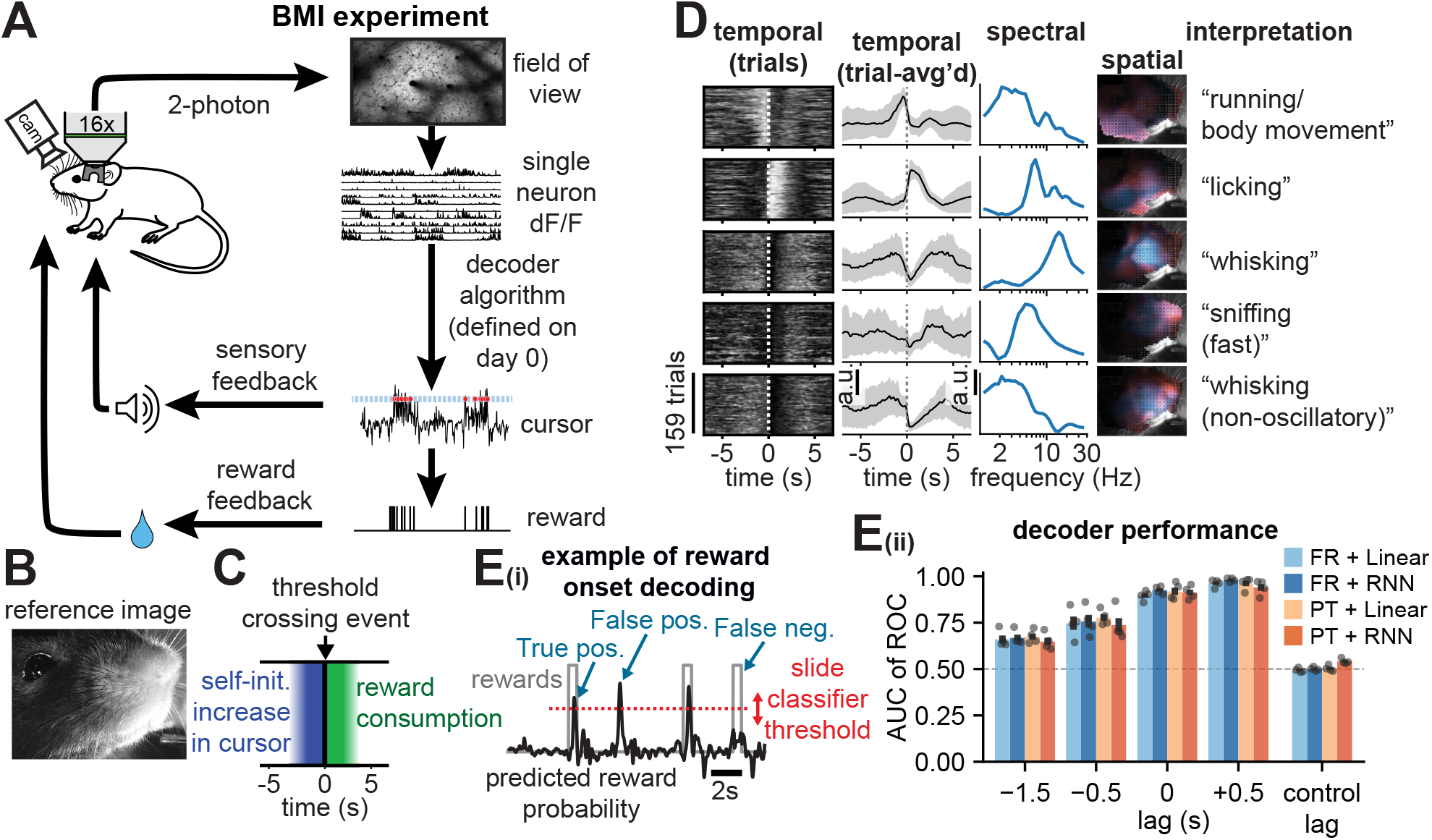
Uninstructed facial behavior predicts brain-machine interface reward events. Uninstructed facial behavior around BMI reward events and decoding analysis. A) Experiment diagram. The mouse performs a brain-machine interface (BMI) task, increasing the activity of a set of neurons selected on a baseline day (‘day 0’) to earn rewards at a threshold (see Methods 5.11). B) Face-camera field-of-view. C) Diagram of a single threshold-crossing reward event. D) Representative example illustrating consistent uninstructed behavioral sequencing around threshold crossings. Rows are FR components from an asymmetric (right-side-only) sliding window (see Methods 5.14.2). Left: temporal-factor rasters aligned to threshold crossings. Left-middle: averaged temporal factors (error bars, s.d. across events). Middle: spectral factors. Right-middle: spatial factors (‘x’ displacement blue, ‘y’ red). Right: interpretation. E) Decoding analysis. Ei) Decoders predict the instantaneous probability of currently achieving reward from either raw tracked points (Points-linear) or FR temporal factors (FR-linear, FR-RNN), assessed by receiver-operating-characteristic (ROC) analysis (see Methods 5.6). Eii) Area under the ROC curve (AUC) for Points-linear, FR-linear, FR-RNN, and Points-RNN versus temporal lag of inputs relative to outputs (negative lags predict future events; controls randomly shifted; Points-RNN uses PCA-reduced points). Shaded regions are s.e.m. across mice (*N* = 4).

**Figure 4 supplement 1.**
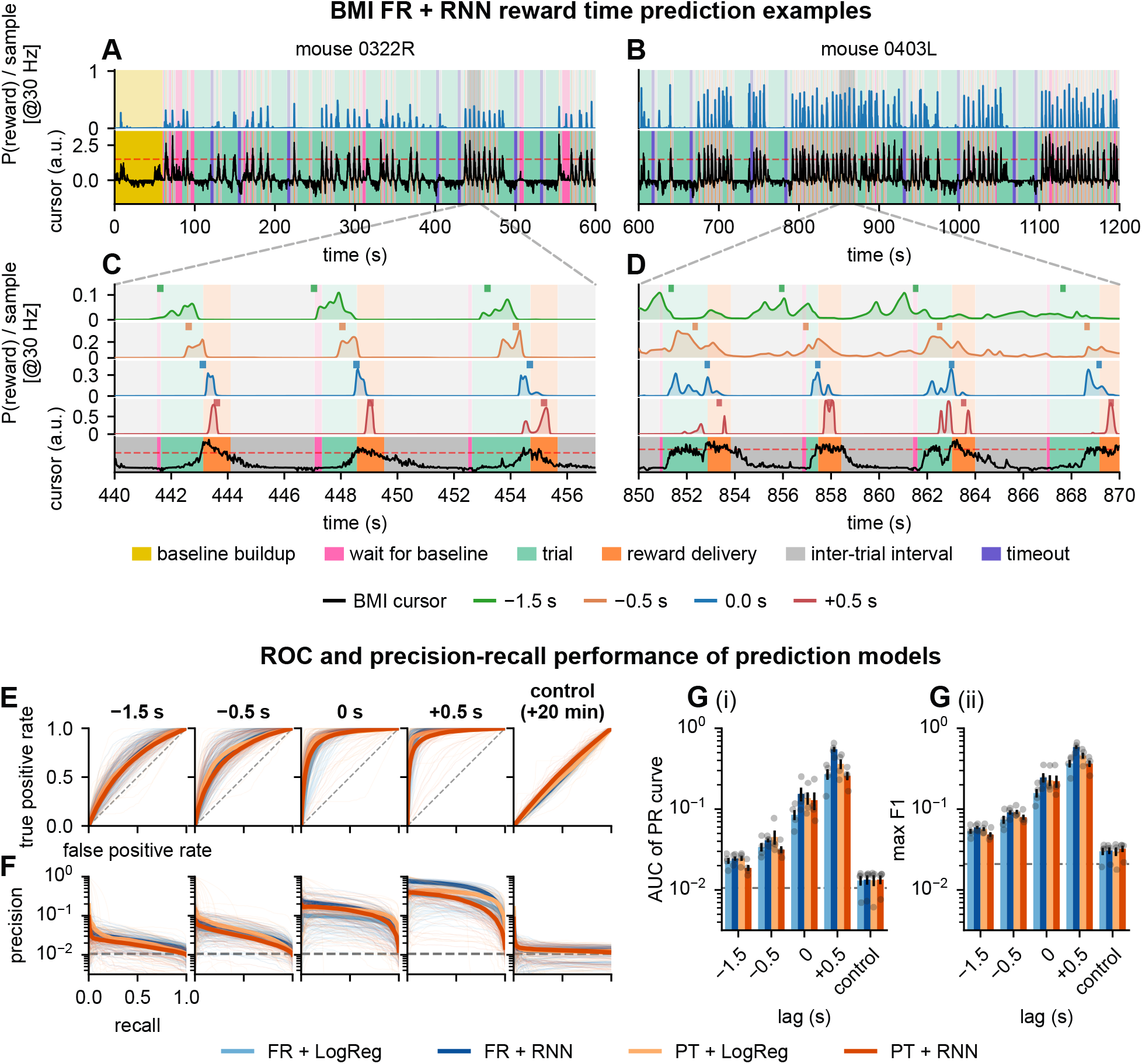
Reward-time prediction from facial behavior during the BMI task. BMI reward prediction model example traces and alternative accuracy metrics. A-B) Representative 600 s segments from two mice showing predicted reward probability *P*(reward) from the FR-RNN model (blue, top) and the online BMI cursor (black, bottom). Background shading marks BMI epochs: baseline buildup (yellow), wait (pink), trial (mint), reward (orange), inter-trial interval (silver), and timeout (purple) (see Methods 5.6). C-D) Zoomed views (∼20 s) of the same sessions, showing FR-RNN *P*(reward) at four target lags (−1.5, −0.5, 0.0, +0.5 s; positive lags predict the future) above the cursor trace. E) ROC curves per lag (columns; including a +20 min shifted control) for the four models (FR-linear, FR-RNN, Points-linear, Points-RNN). Light traces, per-session; bold, across-session mean ± 1 s.e.m.; dashed diagonal, chance. F) Precision-recall curves for the same layout; dashed line, positive-frame prevalence. G) Precision-recall summaries across lags and models. Gi) Area under the precision-recall curve (log scale). Gii) Maximum *F*_1_ (log scale). Bars, across-mice mean; error bars, s.e.m. across mice (*N* = 4); dots, mice. All comparisons against the shifted control are significant (Benjamini-Hochberg FDR; see Methods 5.6.5).

**Figure 4 supplement 2.**
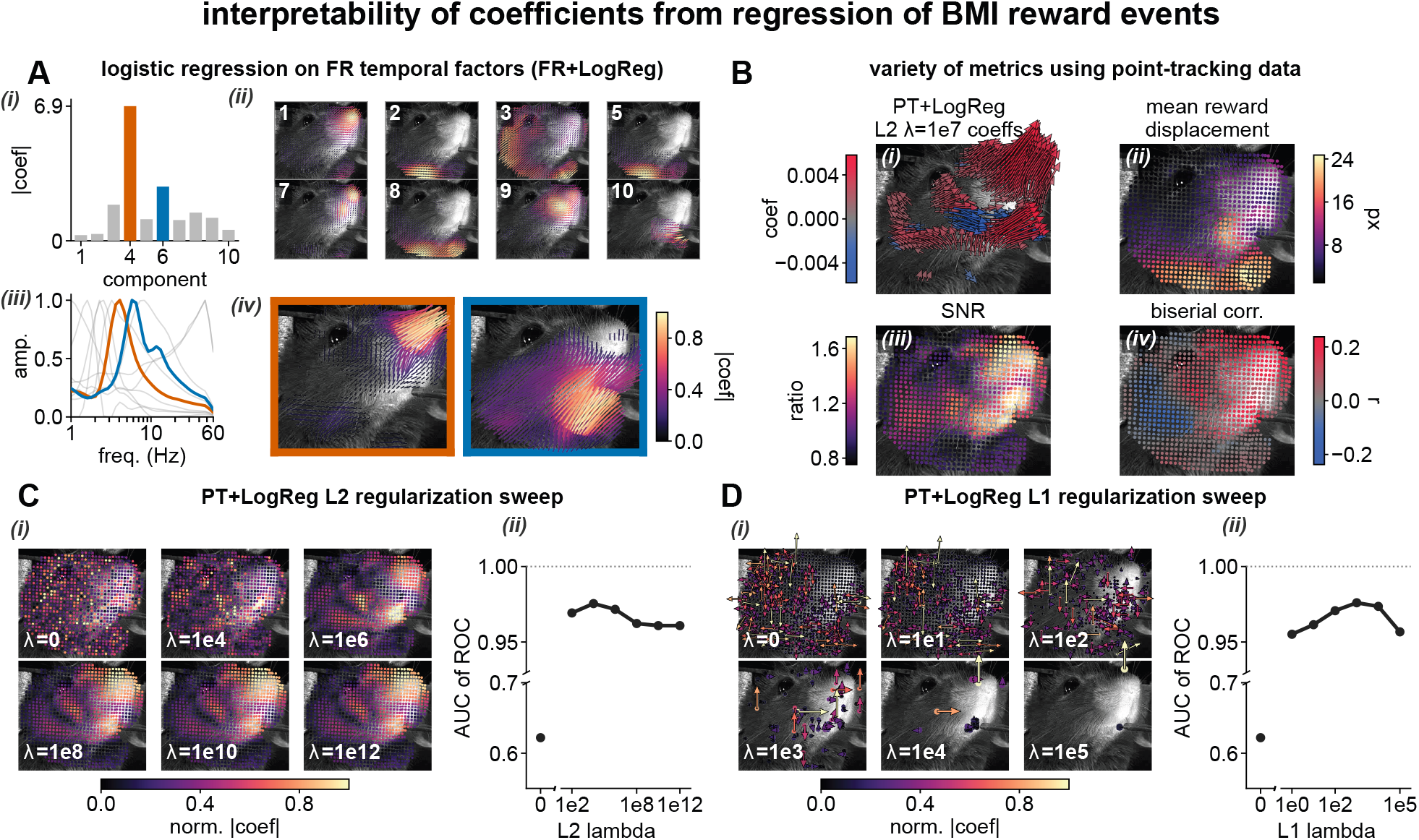
Interpretability of COEFFICIENTS from regression of BMI reward events. Interpretability example for one representative session (see Methods 5.6.6); FR-linear is labeled ‘FR + LogReg’ and Points-linear ‘PT + LogReg’. A) FR-linear decoder coefficients. Ai) Per-component |coefficient| for the 10 FR factors; the two highest-weight components are highlighted (4, orange; 6, blue). Aii) Spatial loadings (‘biquivers’) for the 8 non-highlighted components (angle and length give the relative *x/y* amplitude). Aiii) Spectral factors for all components. Aiv) Enlarged biquivers for components 4 and 6. B) Per-point views of the raw point-tracking data (∼936 points). Bi) Points-linear coefficients as arrows (L2 penalty λ = 10^7^). Bii) Mean reward-triggered displacement. Biii) Signal-to-noise ratio (reward-window / baseline-window mean displacement magnitude). Biv) Signed point-biserial correlation between reward label and signed displacement. C) Points-linear L2 regularization sweep. Ci) |coefficient| maps at six λ values (each face independently max-normalized). Cii) Test AUC versus λ. D) Points-linear L1 sweep. Di) Signed coefficient maps with quivers at L1-selected points. Dii) Test AUC versus λ.

We aligned the FR temporal factors to reward-threshold events, revealing a reproducible sequence of component expression spanning approach, threshold crossing, reward consumption, and the inter-trial interval (**Figure 4D**). We compared four decoders of reward-threshold events (**Figure 4E**; see **Methods 5.6.3, 5.6.2, 5.6.4**): linear and RNN models on FR temporal factors (FR-linear, FR-RNN), a linear model on raw points (Points-linear), and an RNN on PCA-reduced points (Points-RNN). Intermediate spectrograms use causal asymmetric kernels with no future information (see **Methods 5.14.2**). Predicted reward-probability traces align with reward events across sessions (**Figure 4 supplement 1A-D**), and all four models perform similarly across lags (**Figure 4Eii**). At lag 0, AUC is 0.90 ± 0.01 (FR-linear), 0.93 ± 0.01 (FR-RNN), 0.92 ± 0.02 (Points-linear), and 0.91 ± 0.02 (Points-RNN; mean ± s.e.m., *N* = 4 mice), well above shifted controls (0.49–0.54). Precision-recall corroborates these results: area under the precision-recall curve and maximum *F*_1_ exceed an out-of-context shifted control (lag +20 min) at every lag from −1.5 to +0.5 s (**Figure 4 supplement 1E-G**; AUCPR *p*_adj_ ≤ 0.044, max-*F*_1_ *p*_adj_ ≤ 0.019; paired *t*-test, BH-FDR, *N* = 4 mice; see **Methods 5.6.5**). The close agreement across decoders suggests that the decodable reward signal is broadly accessible in both high-dimensional and low-rank FR representations, and that FR yields a more compact, interpretable decoder. Thus, despite the task being conditioned on neural activity alone, mice show stereo-typed uninstructed movements that track task dynamics and reward expectation.

To interpret these decoders, we visualize the regression coefficients for an example session (**Figure 4 supplement 2**). Under an FR-linear model at lag 0, coefficient mass concentrates on a few components, with lateral nose movement at ∼3 Hz as the top predictor of reward onset (**Figure 4 supplement 2A**). The Points-linear decoder and model-free summaries give a complementary view: similar regions around the nose and jaw carry the largest weights, but these loadings are mixed over points and do not expose spectral patterning (**Figure 4 supplement 2B-D**; see **Methods 5.6.6**).

### 3.4 Spectral transformation between facial behaviors and neural activity

The decoding results above show that a spectral transformation of facial movements relates to cognitive and task variables and cortical activity. Together with the existence of central pattern generators for rhythmic facial behaviors, this motivates the hypothesis that a spectral transformation lies between M1 activity and facial movement. We therefore asked whether M1 neural activity is better explained by a direct transformation of facial position or by a spectral transformation of movement (**Figure 5**).

**Figure 5.**
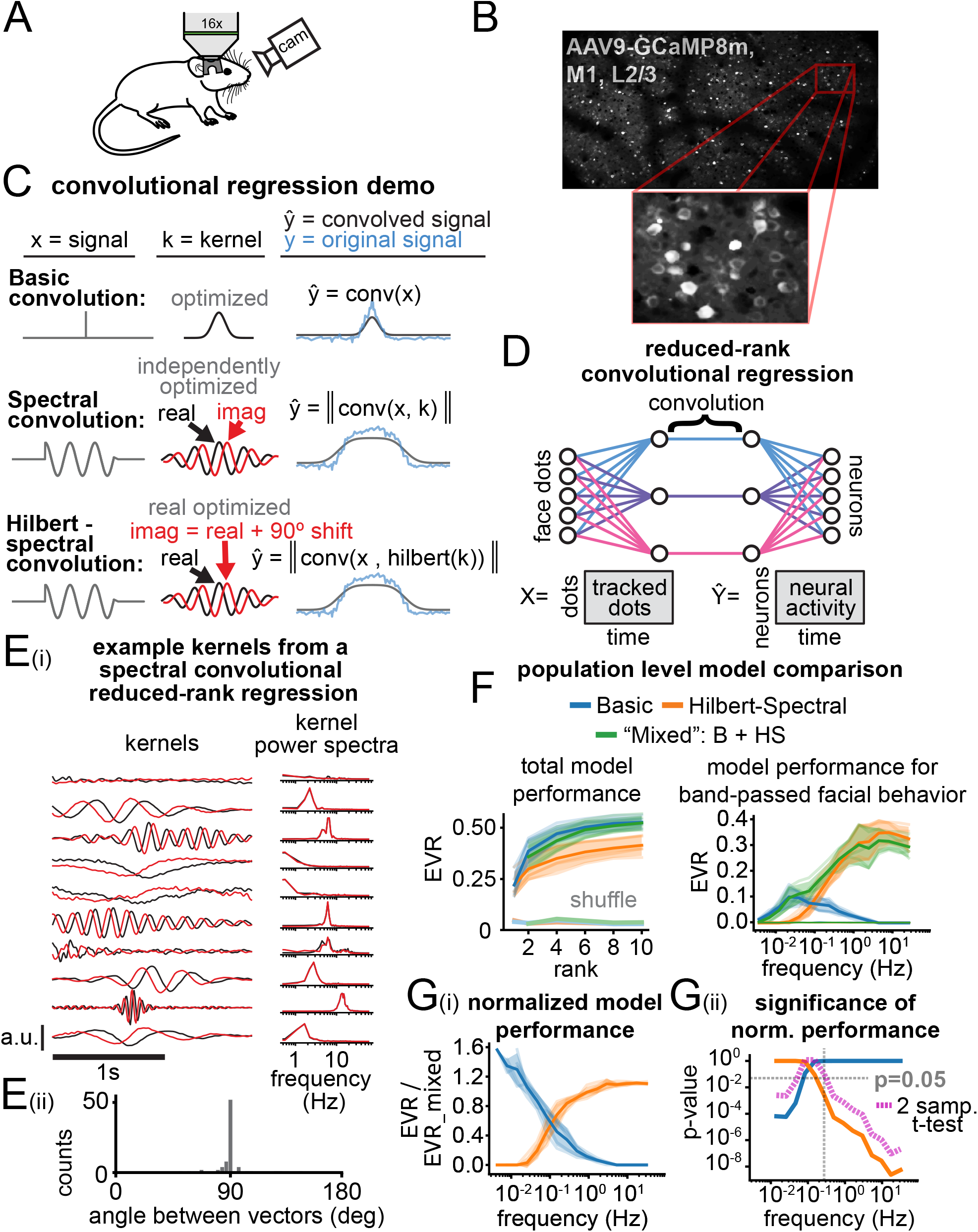
Convolutional reduced-rank regression reveals frequency-dependent phase-variant and phase-invariant coding of facial movement in M1. Convolutional reduced-rank regression of M1 activity from facial point-tracking. A) Experimental setup. The mouse is not performing a task but is on a linear treadmill with open-loop visual/auditory stimulation and uncued infrequent soy-milk droplets (see Methods 5.11.5). B) Example two-photon field-of-view. C) The three convolution models. Basic: a signal is convolved with a real kernel whose shape is optimized to minimize output error. Spectral: a complex kernel with independently optimized real and imaginary parts, taking the output magnitude. Hilbert-spectral: as spectral, but the imaginary part is constrained to the 90° (Hilbert-transformed) version of the real part. D) Reduced-rank convolutional regression. Tracked-point inputs (after only the first pipeline step) are encoded into a reduced space, convolved, then decoded to predict neural activity; optimization is over the encoding matrix, kernels, and decoding matrix (see Methods 5.7). E) Representative spectral-convolution kernels, which naturally form 90°-offset sinusoids after optimization. Ei) Raw kernels (real, black; imaginary, red) and their Fourier transforms. Eii) Angle between real and imaginary components. F) Performance of basic, Hilbert-spectral, and a hybrid (half-each) model; shuffles randomly shifted, error bars s.e.m. across mice. G) Performance by frequency band of the output neural data, with models optimized per band; error bars, s.e.m. across mice. Gi) Raw EVR per model. Gii) EVR ratios of the pure models to the hybrid; significance by paired *t-*test per band with Bonferroni correction across bands. *N* = 5 mice throughout (F–G).

**Figure 5 supplement 1.**
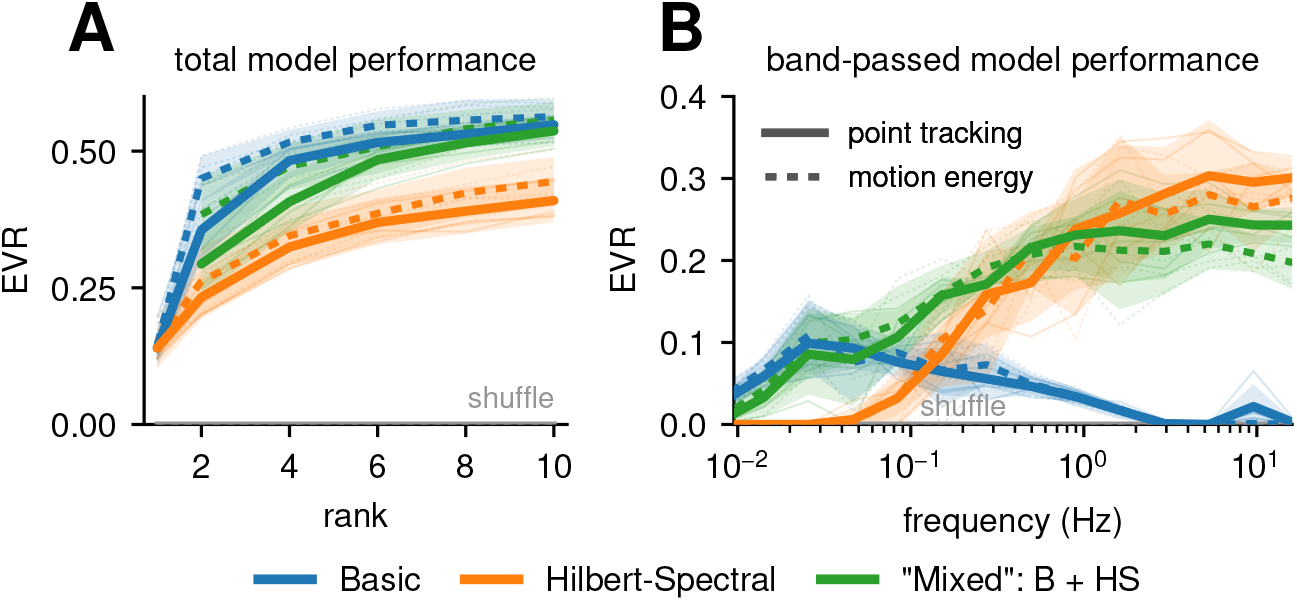
Motion-energy PCA performs similarly to point-tracking in the convolutional reduced-rank regression of M1 activity. Comparison of motion-energy PCA and point-tracking predictors (see **Methods 5.7.5**). FR point-tracking is raw and uncompressed. In both panels, color encodes the model arm (basic, Hilbert-spectral, hybrid) and line style the predictor (point-tracking, solid; motion energy, dashed); thin lines, individual mice; bands, 95% confidence intervals across mice; thick lines, crossanimal mean; grey traces, temporally shifted null (*N* = 5 mice). A) Total held-out EVR versus model rank; each predictor’s ridge penalty tuned independently. B) Frequency-resolved EVR versus the center frequency of the behavioral signals’ filtered band (rank fixed at 6).

**Figure 5 supplement 2.**
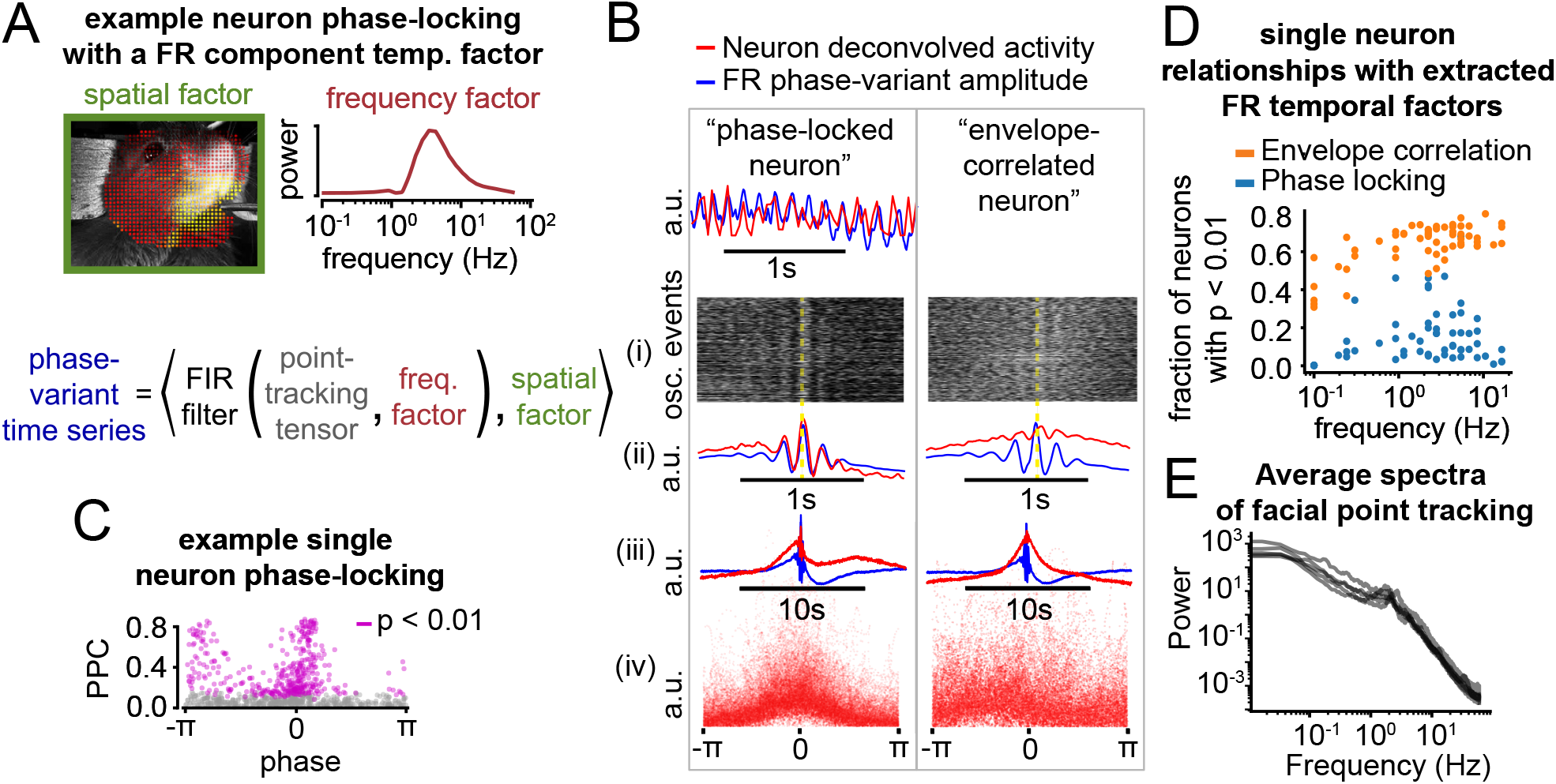
Phase-locking and envelope correlations between facial movements and single neurons. Single-neuron phase-locking and envelope-correlation analysis of facial components. A) Upper: example spatial and spectral factor from one FR component. Lower: phase-variant behavioral signals are generated by FIR-filtering the raw point-tracking signals with the component’s spectral factor and projecting onto its spatial factor (see Methods 5.8). B) Two neurons related to the temporal factor of the component in (A): one phase-locked to the instantaneous movement (left), one correlated with the spectral envelope (right). Bi) Neural-activity raster triggered on oscillatory events (brightness, amplitude; yellow line, behavioral phase zero). Bii) Average over events; overlap of red and blue traces shows phase-locking. Biii) As Bii, zoomed out to show envelope correlation. Biv) Neural amplitude versus instantaneous behavioral phase. C) Phase-locking of all neurons to the component in (A–B). Pairwise phase consistency (PPC) gives the phase-locking strength, and phase the mean phase difference between behavior and neural activity; purple dots are significant against a shift null. D) Fraction of neurons with significant PPC or envelope correlation. Each dot is a behavioral component (rank-12 decompositions; 5 mice, 60 components); *x-*axis is the component’s peak frequency. Significance at *p* < 0.01, Bonferroni-corrected across neurons (see Methods 5.8). E) Example raw power spectra of the behavioral time series for each of the 5 mice.

To assess how much phase information is conserved between behavior and M1 activity, we developed three regression models that handle behavioral phase differently (**Figure 5C**). In each model, the input is convolved with a learned kernel that is optimized along with input and output linear projection weights to minimize output prediction error of neural time series. The models differ in whether this convolution performs a spectral transformation (i.e., taking the magnitude of a complex convolution) and thus whether behavioral phase is retained. The “basic convolution” model uses a real kernel and is phase-variant (it cannot extract an oscillatory envelope). The “spectral convolution” uses a complex kernel with independently optimized real and imaginary parts and takes the output magnitude, which makes it phase-invariant. The “Hilbert-spectral convolution” constrains the imaginary part to the 90° (Hilbert-transformed) real part, giving a phase-invariant model matched in parameter count to the basic model (see **Methods 5.7**). Comparing the three models isolates the relative contributions of phase-variant and phase-invariant representations, and bounds how much M1 encodes about absolute facial position versus the spectral content of movement.

Each convolution model is wrapped in a reduced-rank regression architecture (**Figure 5D**) that constrains the number of kernels to a user-defined “rank”, which regularizes the model and gives each kernel an input-output mapping. The inputs are the raw point-displacement time series (before any spectral processing), and the outputs are the inferred deconvolved event rates of the layer 2/3 M1 neurons (see **Methods 5.7**). The architecture resembles a convolutional neural network, except that the spectral convolutions are defined explicitly in a single layer without nonlinear activations.

The fitted spectral-convolution kernels are approximately 90°-offset amplitude-modulated sinusoids tiling the frequency axis (**Figure 5E**). This illustrates the model’s natural internal representation and motivates a direct comparison of the parameter-matched basic (phase-variant) and Hilbert-spectral (phase-invariant) models. Fitting the first 10 principal components of neural activity, we find that the basic model explains more variance than the spectral model, and that a hybrid model (where half the ranks are basic, and half are Hilbert-spectral; same total parameters as basic) performs similarly to the Hilbert-spectral (**Figure 5F**). We conclude that the major dimensions of neural variance are generally better fit by a phase-variant transformation, and that there is substantial overlap between the dimensions explained by the basic and spectral models (basic vs. Hilbert-spectral EVR at rank 10, *p* = 0.0015, two-sided paired *t-*test, *N* = 5 mice).

Motivated by sensory systems (Bendor and X. Wang 2007; Ewert, Vahle-Hinz, and Engel 2008), we asked whether a movement’s frequency predicts whether M1 represents it via a phase-variant or phase-invariant transformation. Natural behavioral spectra tend to have higher power at lower frequencies (Bassingth-waighte, Liebovitch, and West 1994; Kafetzopoulos, Gouskos, and Evangelou 1997). Consistent with this, the raw point-tracking spectra show high low-frequency power (**Figure 5 supplement 2E**) and power-law decay (*β* = −1.7; *p* < 10^−4^, two-sided Wald test, *N* = 5 mice). We refit the models on movements filtered into narrow frequency bands, and the Hilbert-spectral model outperforms the basic model above 0.5 Hz (**Figure 5G**; paired *t-*test, Bonferroni-corrected *p* < 0.05 per band above ∼0.5 Hz, *N* = 5 mice). This held for both FR point-tracking and pixelwise motion-energy PCA (**Figure 5 supplement 1**; see **Methods 5.7.5**). Higher-frequency facial behavior is therefore better represented in neural activity via a spectral transformation, up to the 30 Hz imaging limit.

We also examined individual neurons’ tuning to phase-variant and phase-invariant movements. Many neurons are visibly phase-locked to facial movements, and others correlate with the oscillatory envelope (**Figure 5 supplement 2A-B**). For each FR component, we built a phase-variant signal and its corollary phase-invariant envelope (see **Methods 5.8**), and for each neuron we computed the pairwise phase consistency with the former and the correlation with the latter (**Figure 5 supplement 2C-D**). Across mice, 93.2% of neurons (4,207/4,513; 93.4 ± 0.8% s.e.m.) show significant envelope correlation to at least one phase-invariant component and 65.4% (2,951/4,513; 66.2 ± 5.9% s.e.m.) are significantly phase-locked to at least one phase-variant component (Bonferroni-corrected, *p* < 0.01; *R* = 12; *N* = 5 mice), with some phase-locked to signals near the ∼15 Hz neural Nyquist limit. More neurons show phase-invariant tuning at higher movement frequencies (permutation test, *p* < 0.01).

### 3.5 Similarity between face-rhythm components and neural activity components

Given the anatomy of the facial motor system and the utility of spectral transformations in explaining neural variance, we hypothesized that FR transformations approximate the inverse of those in the facial motor pathway. This predicts that applying a similar tensor decomposition to neural data would yield temporal factors closely related to FR temporal factors.

We examined whether a one-to-one mapping exists between FR temporal factors and the temporal factors from non-negative tensor decomposition of simultaneously recorded neural data (**Figure 6A**). Mice ran freely on a linear treadmill with infrequent soy-milk rewards while we recorded M1 activity by two-photon imaging and facial behavior by camera. Despite the two analyses being fully independent, the temporal factors are pairwise similar (**Figure 6B**), with greater correspondence for FR components of higher peak frequency (**Figure 6C**). Overall, 41 of 50 matched pairs (82%; 70–90% per mouse, *N* = 5 mice) are significantly correlated (Bonferroni-corrected *p* < 0.001 against 25,000 circular shuffles; median *r* = 0.35, range 0.03–0.69; **Figure 6D**). Thus, the dominant modes of neural activity resemble those of spectrally transformed facial movement.

**Figure 6.**
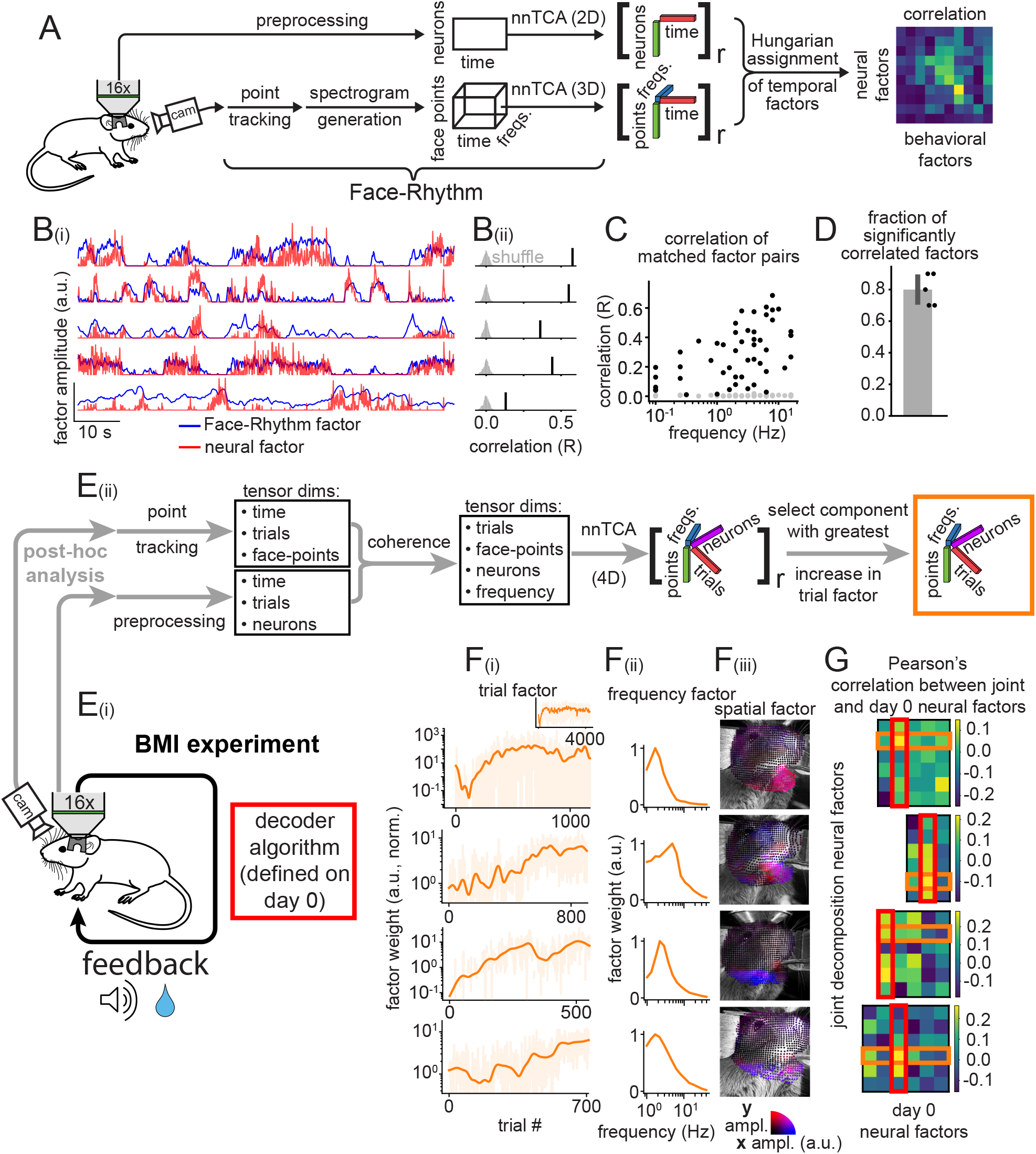
Joint neural-behavioral decomposition reveals learning-associated face-brain coupling. Joint decomposition of simultaneous neural and facial recordings during spontaneous behavior and multi-day BMI learning. A) Analysis pipeline comparing independently extracted tensor components from neural data (top) and face data (bottom) during spontaneous behavior (see Methods 5.11.5). M1 neural data undergo 2D non-negative TCA (equivalent to NMF) and face data the standard FR pipeline; the two sets of temporal factors are 1-to-1 matched by linear-sum (Hungarian) assignment. B) Representative matched temporal factors from one mouse. Bi) Time series of matched factor pairs. Bii) Pearson correlation per pair; shuffles randomly time-shifted. C) Correlation of matched temporal factors versus the peak frequency of the corresponding facial component (black, real; gray, shuffled). D) Fraction of significantly correlated factor pairs (N = 5 mice; Bonferroni-corrected *p* < 0.001 against 25,000 circular shuffles); error bars s.e.m. across animals. E) Joint decomposition during a multi-day BMI task. Ei) Setup: on day 0 (open-loop) one M1 neural factor is selected as the decoder/cursor dimension (red box; see Methods 5.11.6); during BMI sessions, feedback and reward follow activity along this axis. Eii) Post-hoc joint factorization (see Methods 5.9): coherence between all neuron:point pairs forms a 4D tensor (trials × frequencies × points × neurons) decomposed by non-negative TCA into components with trial, spectral, spatial, and neuron factors. The component whose trial factor increases most over sessions is highlighted (orange box). F) Factors of the learning-associated joint component for each of 4 mice (one row per mouse). Fi) Trial factor (selected component, orange; others, light). Fii) Spectral factor. Fiii) Spatial factor (‘x’ blue, ‘y’ red/magenta). G) Pearson correlation between all joint-component neural factors (rows) and all day-0 neural factors (columns) per mouse. The red box marks the decoder axis; the orange box marks the learning-associated component (same as F).

**Figure 6 supplement 1.**
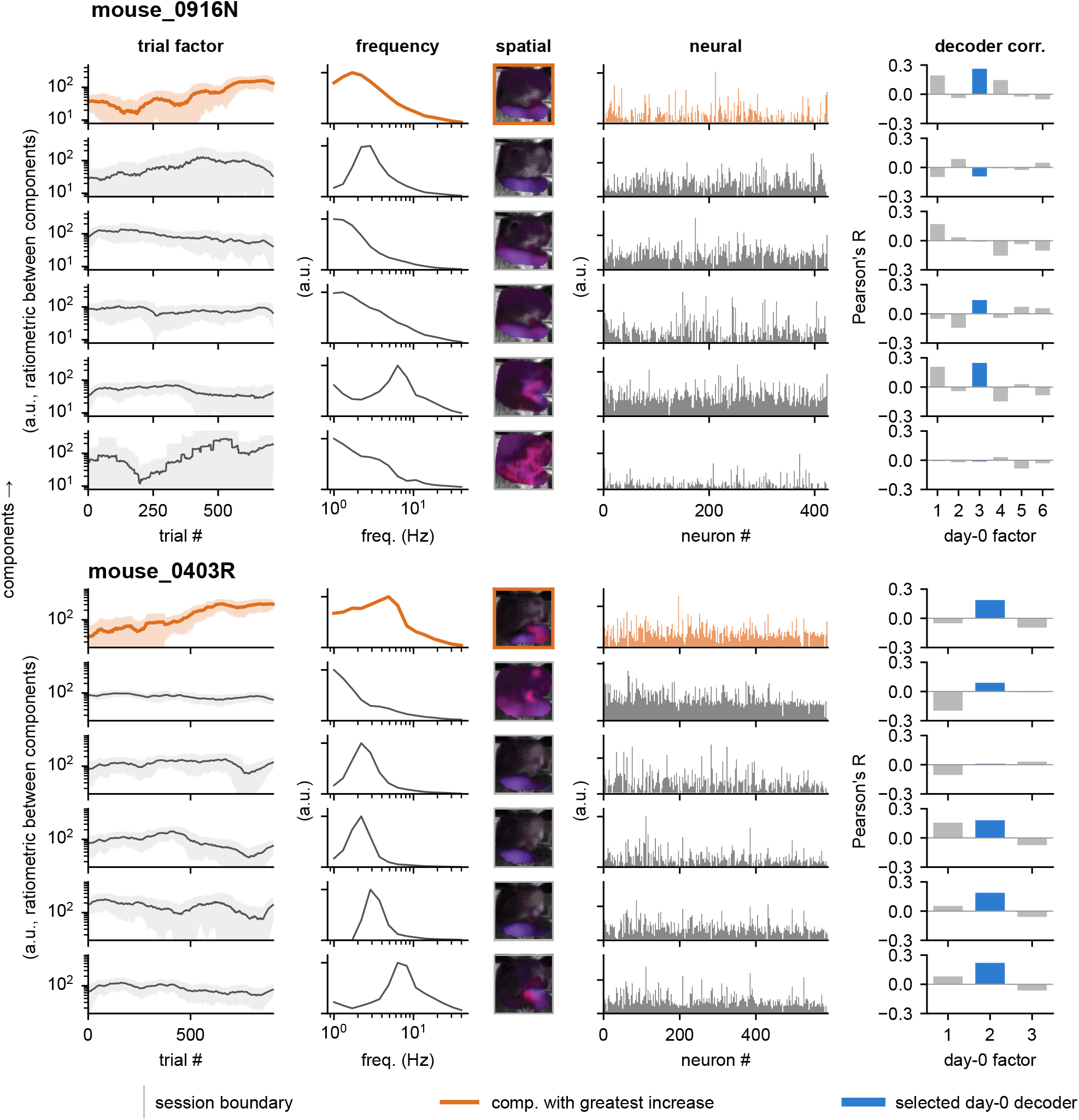
All joint neural-behavioral components for representative mice. Each row is one component from the joint coherence-tensor decomposition (**Figure 6Eii**) for two representative mice. Columns, left to right: trial factor (magnitude over trials), spectral factor, spatial factor (‘x’ blue, ‘y’ red), neural factor (per-neuron loading), and the decoder correlation (Pearson *R* between the neural factor and each day-0 factor; blue bar marks the day-0 decoder factor). The component whose trial factor increases most across sessions is orange (same as **Figure 6F**); others are gray. Shaded regions in the trial column are the interquartile range of a sliding-window smooth.

We therefore developed a joint decomposition of the shared space between FR-transformed behavior and neural activity to identify correlated neural-behavioral motifs. Inspired by canonical correlation analysis, we computed the coherence between all neurons and all face points to form a 4-dimensional tensor (trials × frequencies × points × neurons) and decomposed it by non-negative tensor factorization (**Figure 6Eii**). Each component has four factors: a trial factor (expression across sessions), a spectral factor, a spatial factor (points), and a neuron factor.

We applied this to 4 mice performing a multi-day BMI task (**Figure 6Ei**). The decoder axis is defined on a baseline day (“day 0”) by decomposing neural activity with sparse PCA and selecting one neural factor as the cursor dimension (red boxes, **Figure 6G**; see **Methods 5.11.6**) before BMI training begins. The joint decomposition is then applied post-hoc to subsequent sessions with no knowledge of the chosen decoder, and all components are extracted in an unsupervised manner.

To find the neural-behavioral motif most shaped by learning, we selected the joint component whose trial factor increased most over training (orange, **Figure 6Fi**; all components in **Figure 6 supplement 1**). We then correlated each joint component’s neural factor with each day-0 neural factor (**Figure 6G**) to test whether the learning-associated component (orange box) aligns with the decoder axis (red box). In 3 of 4 mice it was the top match (highest correlation) among the 6 joint components, and the second-highest in the fourth (*p* ≈ 0.004 under a uniform null). Its behavioral factors indicate movements consistent with preparatory licking and running (**Figure 6Fii-Fiii**). Thus, the joint decomposition independently discovers that the neural-behavioral motif most shaped by learning aligns with the task-relevant (decoder) neural dimension, revealing behavioral strategies that become coupled to task-relevant activity during learning.

## 4 Discussion

Face-rhythm extracts interpretable facial behaviors that predict task and internal variables and align closely with demixed neural activity components. Together, our results support spectral analysis as a natural featurization for relating ongoing facial behavior to motor cortical activity.

### 4.1 Comparison to other methods

Automated behavior analysis has advanced rapidly, and many recent tools exploit deep neural networks and semi-supervised frameworks (e.g., DeepLabCut-Live! (Kane et al. 2020), SLEAP (Pereira et al. 2022), VAME (Luxem et al. 2022), CEBRA (Schneider, J. H. Lee, and M. W. Mathis 2023)). Although powerful, large networks are challenging for data-constrained experiments: supervised approaches require manual labeling, and trained networks generalize poorly beyond data resembling their training set. In contrast, FR is end-to-end unsupervised and runs on small or even ‘*N* = 1’ datasets. Its tracking, transformation, and decomposition steps are not trained models and carry no dataset-specific learned biases. FR works best when the behavior moves like a deformable mesh without occlusions, a spectral transformation is a reasonable representation of underlying movement, and a sparse, parts-based, continuous-time description is desirable.

### 4.2 Software and extensibility

FR’s three modular steps (point-tracking, spectral transformation, and tensor decomposition) make it an extensible framework. For instance, FR’s Lucas-Kanade tracking could be replaced with keypoints from DeepLabCut, SLEAP, CoTracker, or FaceMap. Its point-tracking output could feed other decompositions, as shown here, and its nnTCA temporal factors could serve as low-dimensional inputs to segmentation methods such as MoSeq or B-SOiD. We also developed several supporting methods: a mesh-physics-constrained Lucas-Kanade modification that reduces noise and drift, an exact variable-Q transform with flexible filter design, convolutional reduced-rank regression models that build a spectral transformation into the inductive bias, and a coherence-based joint tensor decomposition for discovering shared neural-behavioral components. All code is available (https://github.com/RichieHakim/face-rhythm; see **Methods 5.16**).

### 4.3 Relationship of FR factors to neural activity

Our findings echo the concept of motor primitives (Shadmehr and Mussa-Ivaldi 1994; Hogan 1984; Botvinick, Niv, and Barto 2009) and recent evidence on CPGs in rhythmic behavior (Dempsey et al. 2021; Moore et al. 2013; Kaku et al. 2025), and suggest that M1 multiplexes phase-variant and phase-invariant representations, reminiscent of auditory and somatosensory cortex (Peterson and Heil 2019; Bendor and X. Wang 2007; Ewert, Vahle-Hinz, and Engel 2008). Practically, closed-loop BMIs and neural decoders may benefit from incorporating spectral transformations.

Two of our observations fit existing theories of how rhythmic motor behavior is produced: higher-frequency behaviors are more likely represented in M1 via a spectral transformation, and the major axes of M1 activity align with spectrally transformed behavior. Both are consistent with motor cortex propagating phase-invariant movement signals to lower CPGs, where absolute amplitude is converted into the amplitude of oscillatory components (Cramer and Keller 2006; Hill et al. 2011; Friedman, Zeigler, and Keller 2012).

A spectral-envelope code for higher-frequency movement has implications for coding efficiency and motor architecture: envelopes carry lower frequency content and require less bandwidth and are easier to propagate through the noisy low-rank dynamics of motor cortex (Mastrogiuseppe and Ostojic 2018; Omrani et al. 2017; Laughlin, De Ruyter Van Steveninck, and Anderson 1998), and the rhythmic patterning itself may be compartmentalized into discrete CPGs acting as low-plasticity reservoir networks (Kaku et al. 2025). If cortex sends mainly a phase-invariant envelope to CPGs, fine cortical control of instantaneous phase may be limited, or a secondary descending signal may carry explicit phase information.

### 4.4 Analysis of conditioned uninstructed movements

Subtle uninstructed movements are robustly predictive of BMI performance, highlighting the need for careful behavioral tracking. Such micromovements, though not required to perform the task, may be present in many BMI paradigms, and question the common experimental inclusion criterion of requiring the absence of gross observed movement. More thoroughly identifying uninstructed task-associated behaviors is warranted to interpret how animals achieve control over cursor dynamics. Similar concerns apply to tasks with waiting periods, such as the Pavlovian task here (**Figure 3**), where uninstructed movements and their associated neural activity can produce spurious correlations between movement, cognitive signals, and task variables (Mohebi et al. 2024).

### 4.5 Conclusion

Face-rhythm is an unsupervised, data-efficient method for decomposing facial movement into interpretable components. Applied to rodent facial video, it recovers common behaviors such as whisking, sniffing, and licking, along with subtler movements that track cognitive states, task variables, and the dominant modes of motor cortical activity. These components are consistent across mice, indicating stereotyped motor patterns rather than animal-specific artifacts, and underscore the value of measuring subtle uninstructed movements. Under the conditions used here, most M1 neurons were significantly related to the spectral envelope of higher-frequency movements, and a smaller fraction were phase-locked to instantaneous movement, with individual neurons showing both. Together, these results support phase-invariant spectral representations of movement in rodent facial motor cortex and motivate explicit spectral transformations in models relating cortical activity to ongoing behavior.

## 5 Methods

### 5.1 Face-rhythm pipeline

The standard face-rhythm pipeline consists of three major steps: (1) point-tracking, (2) spectrogram generation, (3) dimensionality reduction via tensor factorization. These steps are separable and composable.

As the data pass through the pipeline, their shape transforms accordingly: the input video has shape *(height, width, time)*; point-tracking produces a *(points, 2, time)* array, where the ‘2’ indexes the *x-* and *y-*displacement of each point; spectrogram generation produces a *(points, 2, frequencies, time)* array (reshaped to a (2 × *points, frequencies, time)* tensor for the factorization; Section 5.1.4); and the PARAFAC decomposition returns, for each of the *R* components, three factor vectors of lengths (2 × *points), (frequencies)*, and *(time)*, corresponding to the spatial, spectral, and temporal factors, respectively.

A user-oriented library has been developed and is available here: https://github.com/RichieHakim/face-rhythm

#### 5.1.1 Defining the region of interest / point positions

A region of the face is circumscribed, and equidistantly spaced points are placed to fill the region. The interpoint distance is set such that there are around 500–3000 points.

For analyses in which multiple sessions or multiple animals are used, a set of points is defined on one mouse face, rigid image registration is performed to align the fields of view, and then the points are warped to matched positions across all recording sessions. Rigid registration is performed using the `roicat.tracking.alignment` module from the `ROICaT` library: https://github.com/RichieHakim/ROICaT

#### 5.1.2 Point-tracking

Point-tracking is performed using the Lucas-Kanade sparse optical-flow algorithm (Lucas and Kanade 1981; Baker and Matthews 2004) and recursively integrating velocities calculated for each point after each frame. Two additional forces were applied to each point after every frame. First, a ‘mesh physics’ force similar to Hooke’s law was used to shape the way individual points could move. Each point was ‘pushed’ and ‘pulled’ by its neighbors to maintain the local inter-point distances. Second, a gentle ‘relaxation’ force was applied to each point toward its initial position to counteract drift.

Optical-flow was chosen for its efficiency and lack of domain priors (i.e., it is not trained on a specific dataset). We use OpenCV’s (Bradski 2000) implementation which exposes a variety of parameters to the user. The optical-flow local integration window size (parameter name: ‘winSize’) is set to be equivalent to roughly 1–3 mm in the field of view, which is typically around 20–60 pixels in our recordings. The optimization parameters were chosen to minimize drift and jump errors (‘maxLevel’ ≈ 3, ‘maxCount’ ≈ 3).

Contrast-limited adaptive histogram equalization (CLAHE) is applied to each frame of the input video prior to point-tracking using OpenCV’s `createCLAHE` class. This normalization step improves robustness to occlusions (such as whisker and tongue movements), changes in lighting, and drift. The face-rhythm pipeline exposes arguments for CLAHE and other parameters.

In each frame *t*, given the prior positions 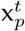 and the original positions 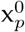, each point *p* is updated in four steps:

##### Step 1. Lucas-Kanade prediction

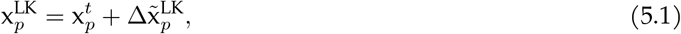

where 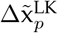 is the displacement vector (i.e., ‘velocity’) obtained from the Lucas-Kanade optical-flow algorithm using OpenCV’s `calcOpticalFlowPyrLK` function.

##### Step 2. Violation freezing

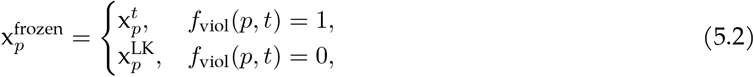

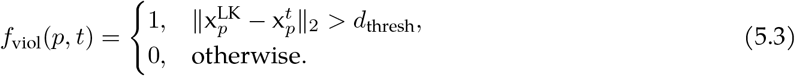

##### Step 3. Mesh correction

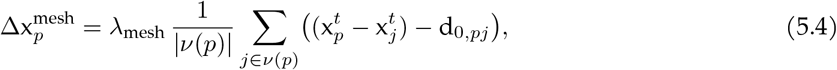

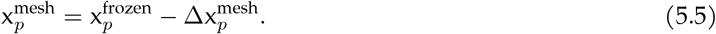

##### Step 4. Relaxation

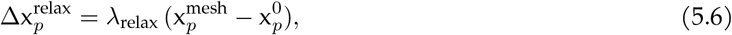

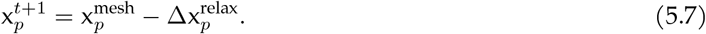

##### Variable definitions

- 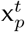: position of point *p* on frame *t*.
- 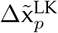 : displacement vector (i.e., ‘velocity’) from the Lucas-Kanade algorithm.
- *ν*(*p*): neighbor indices of point *p* in the mesh.
- *d*_0,*pj*_: initial / ‘resting’ displacement vector (i.e., ‘relative position’) between point *p* and neighbor *j*.
- λ_mesh_, λ_relax_: mesh rigidity and relaxation scaling factors.
- 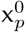 : initial position of point *p*.

##### Output

The resulting point-tracking tensor is denoted as *X* ∈ ℝ^*T* ×*P* ×2^, where *T* is the number of frames, *P* is the number of points, and the last dimension contains the coordinates (*x* and *y*) of each point. The elements of the output tensor contain the tracked position of each point for each time frame.

#### 5.1.3 Spectrogram generation

Spectrograms are computed using a custom implementation of the variable-Q transform (VQT) (Brown 1991; Chatterji et al. 2004) (see Section 5.14 for details), followed by normalization. A spectrogram is computed for each point from the point-tracking output separately for the *x* and *y* dimensions and is denoted as ***S*** ∈ ℝ^*T* ×*P* ×*F* ×2^, where *F* is the number of VQT filters/frequencies. For consistency in future steps, the spectrogram is reshaped to ***S*** ∈ ℝ^2*P* ×*F* ×*T*^, by concatenating the *x-* and *y-*displacements along the points dimension. The time dimension is typically downsampled (∼15 Hz) relative to the original point-tracking tensor (∼250 Hz).

##### Normalization

The spectrogram is normalized in several ways. For subsequent decomposition steps, it is useful to partially equalize along the frequency, temporal, and point dimensions. This normalization boosts the variance allotted to higher frequencies, time periods with less movement, and points with more subtle movements.

The normalization step includes three sequential operations controlled by user-set hyperparameters `spectrogram_exponent` *α*, `normalization_factor` *γ*, and `one_over_f_exponent` *β*:

1. **Exponentiation**

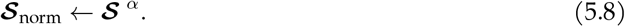
2. **Frame-wise power normalization** The total power at each time point *t* is

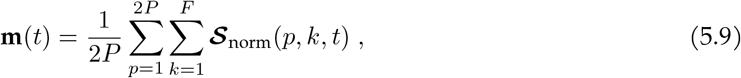

where **m** ∈ ℝ^*T*^, and each time point is rescaled via

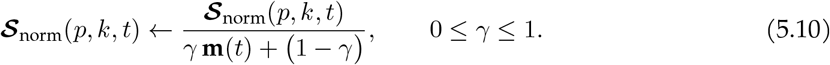

Thus, *γ* = 0 leaves the spectrogram unchanged, while *γ* = 1 enforces identical total power at each time bin.
3. 1/*f* **correction (optional)** To attenuate the natural 1/*f* decay in movement spectra, every frequency slice is multiplied by a power of its center frequency:

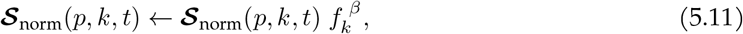

where *β* =`one_over_f_exponent`. *β* = 0 disables the correction; positive *β* flattens the spectrum.

#### 5.1.4 Dimensionality reduction via tensor factorization

Non-negative tensor component analysis (nnTCA) is applied to the spectrogram tensor (which is naturally non-negative) to obtain a set of components. Each output component describes a unique ‘part’ or ‘kind’ of behavior, as defined by each component’s unique combination of spatial, spectral, and temporal factors.

Tensor factorization is performed using CANDECOMP/PARAFAC (CP) decomposition (Harshman 1970; Paatero and Tapper 1994; Carroll and Chang 1970; Mørup, Hansen, and Arnfred 2007; Williams et al. 2018). The primary optimization algorithm used is non-negative hierarchical alternating least-squares (nn-CP-HALS); though for very large tensors, alternative algorithms are used, such as a custom mini-batch implementation of nn-CP-HALS or tensorly’s Alternating Optimization Alternating Direction Method of Multipliers (AO-ADMM) algorithm. The tensorly library was used to perform all tensor factorization operations (Kossaifi et al. 2018).

The input spectrogram tensor is denoted (same as above) as ***S*** ∈ ℝ^2*P* ×*F* ×*T*^ . The output of the tensor factorization is a set of components, each with a spatial, spectral, and temporal factor. The number of components in the decomposition is defined by the user-defined ‘rank’ *R* parameter. Each component can be denoted as a set of vectors (sometimes called a Kruskal-tensor), one for each factor: 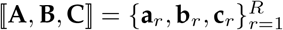, with the following loadings:

- **a**_*r*_ ∈ ℝ^2*P*^ or **A** ∈ ℝ^2*P* ×*R*^: ‘Spatial factor’ loads onto face points (both *x* and y).
- **b**_*r*_ ∈ ℝ^*F*^ or **B** ∈ ℝ^*F* ×*R*^: ‘Spectral factor’ loads onto VQT filters.
- **c**_*r*_ ∈ ℝ^*T*^ or **C** ∈ ℝ^*T* ×*R*^: ‘Temporal factor’ loads onto time bins.

The components can be combined to reconstruct a rank-*R* approximation of the original spectrogram tensor. The reconstruction is given by the following equation using the outer product representation:

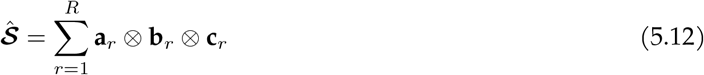

where ⊗ denotes the outer product operation. Alternatively, it can be expressed using Einstein summation notation (see Table 6):

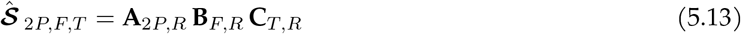

##### Non-negative CP decomposition

Optimization with non-negative constraints is performed over the component vectors to minimize the reconstruction error (mean squared error, MSE) between the original spectrogram tensor and the reconstructed tensor:

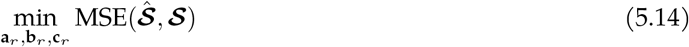

In most cases, the initialization of the component vectors is done using truncated singular value decomposition (SVD) on the reshaped spectrogram tensor, which provides deterministic output components.

#### 5.1.5 Rank selection

Rank is a user-selected hyperparameter that we chose by balancing reconstruction quality, decomposition stability, downstream utility, and interpretability. To evaluate rank systematically, we computed a set of complementary diagnostics across different ranks *R* ∈ {2, 4, 6, …, 20} . Unless otherwise noted, high-order SVD (HOSVD) initialization was used. These diagnostics are summarized in **Figure 1 supplement 1** (panels A-E): panel A shows reconstruction error versus rank, panel B outlines the cross-initialization matching workflow, panel C reports per-mode and combined component consistency, panel D reports BIC, and panel E reports effective dimensionality of each factor matrix.

##### Reconstruction error and explained variance (panel A)

For each rank, we computed the relative reconstruction error

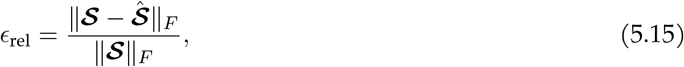

and the explained variance ratio

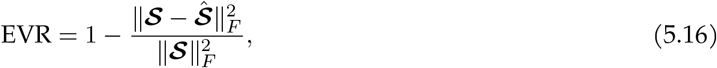

where ***S*** is the input tensor and 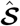 is the CP reconstruction (Eq. 5.12).

##### Component consistency metric

To quantify cross-run reproducibility, we used the consistency framework described previously for tensor decomposition (Williams et al. 2018). For this analysis, we used the BMI dataset (see Section 5.6) and ran 30 random initializations per rank. For a pair of runs *u* and *v*, with factor matrices **A**^(*u*)^, **B**^(*u*)^, **C**^(*u*)^ and **A**^(*v*)^, **B**^(*v*)^, **C**^(*v*)^, we first computed per-mode cosine similarities between component *r* in run *u* and component *s* in run *v*:

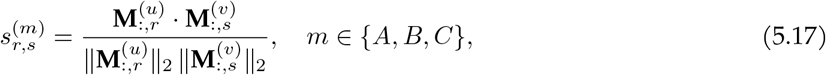

where 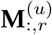 denotes a single factor vector from mode *m* and initialization *u*. These per-mode similarities were combined into a joint cross-similarity matrix by taking the product across modes,

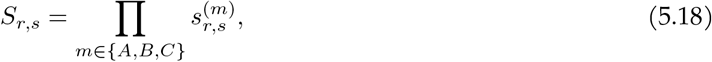

and the Hungarian algorithm was used to find the assignment *π* that maximized 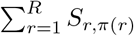.After matching, we evaluated consistency using Pearson correlation *ρ* between matched factor vectors. We report two summary statistics as a function of rank: (1) the per-mode consistency, defined as the Pearson correlation between matched components averaged over components and run pairs, reported separately for each mode; and (2) a combined product consistency score,

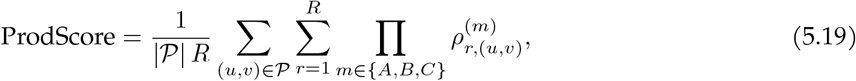

where 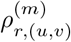 is the Pearson correlation for matched component *r* in mode *m* for run pair (*u, v*) and *P* is the set of all 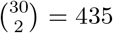 run pairs. The product score provides a stricter measure of agreement that is sensitive to poor consistency in any individual mode.

##### BIC (panel D)

Under a Gaussian residual model, we computed

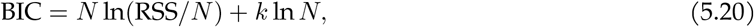

with

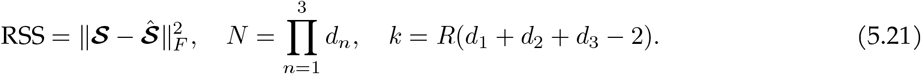

Here, RSS is the residual sum of squares, *N* is the total number of observed tensor entries, and *k* is the effective number of free parameters in the rank*-R* CP model. In this setting, *d*_1_, *d*_2_, *d*_3_ are the mode dimensions of ***S***(spatial, spectral, temporal), so *N* = *d*_1_*d*_2_*d*_3_. The expression for *k* removes two redundant scaling degrees of freedom per component. Lower BIC indicates a better fit-complexity tradeoff. Because non-negative spectrogram residuals violate IID Gaussian assumptions, and because temporal/spatial correlations reduce effective sample size, we treated BIC as a supplementary heuristic rather than a formal model-selection test. We report BIC due to its stronger complexity penalty, but we note that AIC (Akaike information criterion) gives nearly identical scores.

##### Effective dimensionality of factor matrices (panel E)

To quantify redundancy across extracted components, we performed PCA on each factor matrix 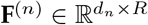 (spatial/points, spectral/frequencies, temporal) and computed the effective dimensionality/rank required to explain 90% variance:

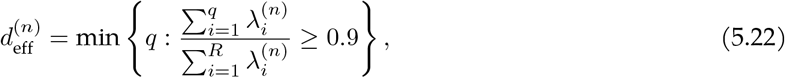

where 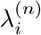 are PCA eigenvalues for mode *n*. Values near *R* indicate decorrelated components in that mode, and values below *R* indicate increasing collinearity/redundancy.

In practice, final rank selection reflects these diagnostics, but is balanced against human interpretability of component factors and downstream utilities like regression.

### 5.2 Cross-animal component consistency

To assess whether FR components generalize across individuals, we extended the within-session consistency framework (Section 5.1.5) to the cross-animal setting. Spectrogram tensors were computed for 13 baseline recording sessions across 5 mice using the standard FR pipeline. For each session, 30 non-negative CP decompositions were run with random initializations at each rank *R* ∈ {2, 4, 6, …, 20}, and the run with the lowest reconstruction error was selected for cross-animal analysis. Point positions were aligned across mice using rigid image registration (Section 5.1.1).

Because temporal factors are session-specific, cross-animal component matching used only the spectral and spatial factors, and not the temporal factors. For each cross-animal session pair, per-mode cosine similarities were computed and combined into a joint cross-similarity matrix 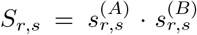, following the same procedure as Section 5.1.5 but restricted to two modes. The Hungarian algorithm was used to find the optimal one-to-one assignment *π*^∗^ maximizing ∑ _*r*_ *S*_*r,π*(*r*)_.

After matching, similarity was evaluated using Pearson correlation *ρ* between matched factor vectors. The product score for a session pair was defined as:

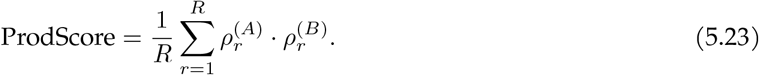

To assess whether cross-animal similarity exceeded chance, a one-sided permutation test was performed. For each of the 59 cross-animal session pairs (pairs of sessions from different animals), 1000 random component assignments were drawn and the mean product score was computed. The p-value was the fraction of permutations in which the mean null score (averaged across all cross-animal pairs) exceeded the actual mean cross-animal product score.

### 5.3 Ablation of point-tracking corrections

To quantify the contribution of each correction applied during point-tracking (Section 5.1.2), we performed a full 2 × 2 × 2 factorial ablation across three factors: mesh correction (M), relaxation (R), and violation detection with point pausing (V). This yielded eight conditions (Table 1), run on *n* = 9 sessions from 7 mice (∼60 min each, 63.56 Hz, 1985 tracked points per session).

**Table 1.** Ablation conditions. λ_mesh_: mesh rigidity; λ_relax_: relaxation strength; *d*_thresh_: displacement threshold for violation detection (px; “off” = disabled). Production parameters: λ_mesh_ = 0.1, λ_relax_ = 0.01, *d*_thresh_ = 300 px.

| Label | Condition | $\lambda_{\text{mesh}}$ | $\lambda_{\text{relax}}$ | $d_{\text{thresh}}$ |
| --- | --- | --- | --- | --- |
| MRV | full model | 0.1 | 0.01 | 300 |
| MV | no relaxation | 0.1 | 0 | 300 |
| RV | no mesh | 0 | 0.01 | 300 |
| V | pure LK | 0 | 0 | 300 |
| MR | no violations | 0.1 | 0.01 | off |
| M | M only | 0.1 | 0 | off |
| R | R only | 0 | 0.01 | off |
| none | none | 0 | 0 | off |

The following Lucas-Kanade parameters were used: `winSize`= [40, 40], `maxLevel`= 3, `maxCount`= 3, mesh neighbor count |*ν*(*p*)| = 24.

#### Ablation metrics

Tracking quality was evaluated using three per-frame metrics: (1) *cumulative drift*: 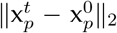, displacement of each point from its initial position; (2) *neighbor velocity coherence*: mean cosine similarity of unit-normalized velocity vectors between each point and its 4 nearest mesh neighbors (velocity estimated as derivative between point positions from consecutive frames), where values near 1 indicate locally coherent motion; and (3) *violation rate*: fraction of points exceeding *d*_thresh_ on any given frame.

#### Statistical comparisons

Each correction was assessed by comparing the conditions in which the correction was present against those in which it was absent (one-sided Wilcoxon signed-rank test across *n* = 9 sessions; *p* = 0.002 for each). Specifically, relaxation was tested on final cumulative drift (with-R {MRV, RV} versus without-R {MV, V}), mesh physics on final neighbor velocity coherence (with-M {MRV, MV} versus without-M {RV, V}), and violation detection on violation rate (with-V {MRV, MV, RV, V} versus without-V {MR, M, R, none}).

### 5.4 Widefield imaging regression comparison analysis

#### 5.4.1 Video motion energy and PCA analysis

To capture facial movements from the behavioral video, we computed the per-pixel absolute difference between consecutive frames to generate the video motion energy (video ME). To isolate movements of specific parts of the face, we manually selected a rectangular area of 50 pixels around the nose, jaw, or whiskers and created the average ME across all pixels within each area. To reduce the dimensionality of the behavioral data, we then used principal component analysis (PCA) to compute the 200 highest-variance dimensions. These dimensions accounted for at least 90% of the total variance in the data, indicating that 200 dimensions were sufficient to describe the main temporal dynamics in the video. We then used the PCA scores as temporal components of animal behavior and used either 10 or all 200 components in the linear encoding model to predict cortical dynamics.

#### 5.4.2 Widefield imaging and ridge regression analysis

Experimental details of the cortical widefield imaging are described previously in Musall et al. 2019 (Musall, Kaufman, et al. 2019). For data processing, we reduced the dimensionality of the video data with singular value decomposition (SVD) and retained the top 200 components, which explained ≥ 95% of the total variance. SVD yielded spatial components (U; pixels × components), temporal components (VT; components × frames), and singular values (S). Subsequent analyses were performed on the product SVT to reduce computational load. SVT was high-pass filtered above 0.1 Hz using a zero-phase lag, second-order Butterworth filter. To project results back into pixel space, outputs were multiplied with U. All widefield data were rigidly aligned to the Allen common coordinate framework.

To link behavioral video components to cortical activity, we used a linear encoding model with a ridge penalty where the regularization penalty λ was estimated separately for each column of the widefield data (Couto et al. 2021; Musall, Sun, et al. 2023). The same model structure was used for all comparisons of cross-validated EVR in **Figure 2**, using either ME PCA, FR components, or point-tracking data as model regressors to predict cortical activity.

To account for the nested structure of sessions within mice, we additionally fit a linear mixed-effects model (LMM) to the cross-validated EVR values. The model was specified as EVR ∼ method + (1|mouse), with method (FR vs. ME PCA) as a fixed effect and mouse as a random intercept, fit using restricted maximum likelihood (REML) via the `statsmodels MixedLM` implementation in Python. Significance is assessed with the Wald z-test on the estimated coefficient (`statsmodels` default). Animal-level paired *t-*tests were computed on per-mouse mean EVR values. We also report the non-parametric Wilcoxon signed-rank test outputs alongside each main parametric comparison.

#### 5.4.3 Decomposition ablation analysis

To evaluate the contribution of each pipeline stage, we compared the standard FR decomposition (rank 10 non-negative TCA on VQT magnitude spectrograms) against alternative decompositions spanning three families. Spectrogram-domain methods operate on the VQT spectrogram tensor with the original shape ℝ^2×*P* ×*F* ×*T*^, which is flattened along the *x-* and *y-* coordinate dimensions to give ℝ^2*P* ×*F* ×*T*^ . The nnTCA-spectrogram (rank 10) decomposition operates directly on this 3D tensor, while all 2D decompositions operate on a flattened version of this tensor (ℝ^2*PF* ×*T*^ ); these include PCA (rank 10 and 1000), NMF (rank 10), and related decompositions.

Point-tracking-domain methods operate on the raw displacement traces (prior to any spectral transformation) with the original shape (ℝ^2×*P* ×*T*^ ). For the “TCA: 3D unflattened point-tracking rank=10” decomposition only, the original 3-way point-tracking tensor was used, preserving the separation between spatial and coordinate dimensions. For matrix decomposition methods like PCA, the *x-* and *y-*coordinate dimensions were first concatenated along the spatial dimension to form a 2D matrix (ℝ^2*P* ×*T*^ ). For the “PCA: lagged point-tracking” decomposition, time-shifted versions of the 2D matrix were concatenated to construct a truncated Hankel (time-delay embedding) matrix with 200 lags (∼3.1 s), then PCA (rank 1000) was performed; anticausal, causal, and centered lag variants were tested and the center lag is presented in the main text. For the “TCA: band-passed point-tracking rank=10” decomposition, the 2D flattened point-tracking matrix was band-pass filtered using the same spectral filters as the VQT spectrograms, then the resulting band-passed tensor was reshaped to ℝ^2*P* ×*F* ×*T*^ and nnTCA (rank 10) was performed on this tensor.

All decompositions were aligned to the widefield imaging timebase and evaluated using the same cross-validated ridge regression framework (Section 5.4.2). The same set of decompositions was applied to each alternative input source (FaceMap, CoTracker, CEBRA, and ViT-MAE; see below), enabling controlled comparisons across tracking and embedding methods. For decompositions (like ‘PCA with rank=1000’), the requested rank serves as an upper bound on the number of extracted components: the achieved rank equals min(requested rank, input feature dimensionality). For example, using CEBRA embeddings with 10 dimensions as input, requesting rank 1000 yields the same result as requesting rank 10.

#### 5.4.4 FaceMap comparison

To compare FR’s dense optical-flow tracking against sparse keypoint tracking, we ran FaceMap (Syeda et al. 2024) on the same widefield-session videos (7 mice, 1–2 sessions per mouse, 9 sessions total). FaceMap uses a neural-network-based detector to track 15 facial keypoints, yielding a point-tracking tensor ***X***_FM_ ∈ ℝ^*T* ×15×2^. We finetuned the FaceMap keypoint tracking model through an iterative process of manual labeling (15-frame batches), retraining (finetuning), and visual inspection on held-out sessions. After six iterations (90 total frames; ∼180 active labeling minutes), performance stabilized across all seven mice and nine sessions. We then deployed this final model to extract keypoints across all sessions. These trajectories were passed through the same decomposition ablation pipeline used for FR: VQT spectrograms followed by each decomposition method (PCA, ICA, NMF, nnTCA, and Hankel-PCA variants) at matched ranks. FaceMap keypoints were also evaluated directly as regressors (without spectral transformation) via the same ridge regression framework described in Section 5.4.2, allowing a controlled comparison of point-tracking frontends while holding all downstream analysis steps constant.

#### 5.4.5 CoTracker point-tracking comparison

To test whether FR’s custom Lucas-Kanade tracker could be replaced by a modern deep-learning tracker, we applied CoTracker v3 (Karaev, Rocco, et al. 2024; Karaev, Makarov, et al. 2024) to the same behavioral videos used in the widefield regression analysis. CoTracker is a transformer-based dense point tracker; we used the pretrained offline model (`scaled_offline.pth`) with 6 refinement iterations. Query points were initialized at the FR mesh positions (∼1,481 points per session). Videos were processed in non-overlapping 300-frame chunks, with tracked positions carried across chunk boundaries.

CoTracker outputs absolute pixel coordinates; to obtain displacement signals comparable to FR, the temporal mean position was subtracted from each point. The resulting displacement tensor was processed through the full decomposition ablation pipeline (Section 5.4.3), aligned to the widefield timebase, and evaluated via the same ridge regression framework (Section 5.4.2).

#### 5.4.6 CEBRA comparison

To evaluate whether a learned nonlinear embedding could substitute for the explicit spectral transformation in FR, we applied CEBRA (Schneider, Lee, and Mathis 2023), a contrastive-learning method that optimizes an InfoNCE objective (Oord, Li, and Vinyals 2019) to map high-dimensional time series into low-dimensional embeddings. We used the self-supervised CEBRA-Time variant (positive pairs defined by temporal proximity, no behavioral labels). The encoder was a single-frame MLP (`offset1-model`, 128 hidden units, GELU activations, *L*^2^-normalized output), trained with batch size 1024, learning rate 10^−4^, for 5,000 iterations. Input to CEBRA was the first 50 PCA components of the flattened FR point-tracking data. Embeddings were computed at dimensions 10 and 50. We chose these two dimensions deliberately: *d* = 10 matches the rank used for FR and the other decompositions throughout this work, enabling a like-for-like comparison at equal dimensionality, while *d* = 50 provides a higher-capacity embedding that tests whether additional dimensions improve predictivity.

Each embedding matrix was reshaped to *T* × *d* × 1 (where *d* is the embedding dimension) and processed through the full decomposition ablation pipeline, including VQT spectrogram computation, all decomposition methods, temporal alignment, and ridge regression evaluation.

##### CEBRA architecture and hyperparameter sweep

To find the CEBRA architectural parameters that best predict cortical activity, we performed a sweep over encoder architectures, contrastive time-offsets, and embedding dimensions. We crossed three encoder receptive-field architectures (`offset1-model`, single frame; `offset10-model`, ∼10 frames at the ∼60 frames/s frame rate; and `offset20-model`, a 20-frame window with a stride length of 4) with contrastive time-offsets of {1, 3, 10} frames and embedding dimensions {10, 50} (14 configurations total; `offset20-model` was evaluated only at time-offset 10). To avoid confounding architecture with tuning quality, each cell’s remaining CEBRA hyperparameters (learning rate, training iterations, hidden-unit count, and weight decay) were tuned independently by Optuna (Tree-structured Parzen Estimator sampler) (Akiba et al. 2019) to directly maximize the downstream objective: the 10-fold block-cross-validated EVR of the widefield ridge regression (Section 5.4.2). Each tuned embedding was then evaluated through the full decomposition / ‘ablation’ battery (Section 5.4.3) over *n* = 9 sessions from 7 mice, using the same analysis window as the main widefield regression figure. The single-frame `offset1-model` performed comparably to or better than the wider architectures at matched embedding dimension and time-offset (**Figure 2 supplement 2**); therefore, it was selected for the main comparison (**Figure 2, Figure 2 supplement 1C**).

#### 5.4.7 ViT-MAE feature extraction

Motivated by BEAST (Wang et al. 2025), which demonstrated that vision-transformer representations of behavioral video predict neural activity, we evaluated frozen ViT-MAE embeddings as an alternative behavioral representation. We used a ViT-Base/16 model pretrained with masked autoencoding (He et al. 2021; Dosovitskiy et al. 2021) (checkpoint: `facebook/vit-mae-base`). To restrict the encoder to the same facial region used by FR, video frames were masked to the face region defined for FR point-tracking (Section 5.1.1); pixels inside the region retained their original values, and pixels outside were set to zero. Masked frames were then resized to 224 × 224 pixels, normalized to ImageNet statistics, and passed through the encoder with masking ratio 0. The CLS-token embedding (768 dimensions) was extracted per frame using batched inference (batch size 128, float32).

CLS embeddings were aligned to the widefield timebase by linear interpolation. For compatibility with the decomposition pipeline, the 768-dimensional embedding was reshaped to 384 × 2 pseudo-channels; this reshaping is numerically transparent because all decomposition methods except TCA operate on a flattened 2D matrix and are therefore invariant to the (768, ) versus (384, 2) representation. VQT spectrograms were computed on all 768 channels using the same parameters as for FR data. The full decomposition ablation suite was then applied. We additionally evaluated 200-dimensional SVD and 10-dimensional PCA reductions of the CLS embeddings directly via ridge regression.

### 5.5 Pavlovian conditioning analysis

To analyze the utility of FR components in describing uninstructed behaviors during Pavlovian conditioning, we used a separate facial video dataset acquired during the high- and low-reward probability classical conditioning tasks described previously (Matsumoto et al. 2016). These facial video data were not analyzed in that study and are distinct from the 60-Hz eye-video dataset used there for eye-blinking analysis.

#### 5.5.1 Behavioral paradigm and data acquisition

Thirsty mice were head-fixed and randomly presented with one of three conditioned stimuli (CS) odors on each trial:

- CS+: odor predictive of a water reward,
- CS0: odor predictive of no outcome,
- CS-: odor predictive of an aversive air puff.

In the high-reward probability task, reward was delivered in 90% of CS+ trials, whereas in the low-reward probability task, reward was delivered in 20% of CS+ trials; air puff was delivered in 90% of CS-trials in both task variants, as described previously (Matsumoto et al. 2016). Each trial consisted of a 1 s CS odor presentation, followed by a 1 s delay epoch and then the unconditioned stimulus (US). Analyses were restricted to cued trials, yielding 240 analyzed trials per session (80 trials per CS class), across 20 sessions from 5 animals. Behavioral event times were taken from the original experimental records for alignment of FR temporal factors to task events. Continuous video of the face was recorded at ∼ 30 Hz. The standard face-rhythm pipeline was used to extract 10 temporal factors using right-asymmetric (‘causal’) kernels/filters (see Section 5.14.2 for details). The four temporal factors shown in **Figure 3D-E** were selected based on visual inspection for display purposes only; all 10 temporal factors were used for the classification analysis.

#### 5.5.2 Elastic net CS classifier

CS identity was predicted from FR temporal factors using multinomial logistic regression with elastic net regularization (L1+L2 penalty). For each trial, the 10 temporal factor traces were concatenated across all time bins in a [ − 2.5, +1.9] s window relative to CS onset ( ≈ 25 bins per factor, ≈ 250 features total), ending one time bin before US onset (2.0 s) to exclude the US-evoked response. Regularization strength was selected via cross-validation within each session, and classifier performance was evaluated using stratified k-fold cross-validation (*k* = 5, 20 repeats).

For each session, a confusion matrix (rows: true class, columns: predicted class) was computed by summing across folds and row-normalized so that each entry gives *P*(*ĉ* | *c*). The null expectation for each cell is the marginal prediction frequency *P*(*ĉ*), estimated from the column sums of the raw confusion matrix; under a classifier with no information about the true class, all rows of the confusion matrix are identical and equal to these marginal frequencies (sometimes referred to as a ‘mean expectation model’). Per-animal means were computed by averaging across sessions within each animal. Significance for each cell was assessed with a two-sided one-sample *t-*test (*N* = 5 animals), testing whether the observed confusion-matrix entry differed from the corresponding null expectation P(*ĉ*) of 33.3%. A multiple-comparison Bonferroni correction over classes would give *α*/3 = 0.017.

### 5.6 Decoding of BMI reward events

We compared four decoders that were trained to predict reward events (onset times of reward-triggering threshold crossings). These models make up a 2 × 2 matrix of input data type × architecture: input data include (FR temporal factors vs. raw point-tracking) and model classes include (linear vs. RNN). We reference each as FR-RNN, FR-linear, Points-linear, and Points-RNN. All four share a common analysis framework, and we describe model-specific details below. See Section 5.11 for the behavioral paradigm and data acquisition.

#### 5.6.1 Shared procedures

The goal of each decoder is to classify the instantaneous probability of a reward event at each video frame from either the FR temporal factors or the raw point-tracking data. Reward times are converted to a Boolean target vector ***y*** ∈ {0, 1} ^*T*^ with value 1 at each reward-event frame, widened by 200 ms in the backward (causal) direction. To test the temporal dependence of decoding on reward events, the target vector is shifted/lagged by Δ*τ* before each training run: four ‘in-context’ lags Δ*τ* {−1.5, −0.5, 0, +0.5 } s relative to reward delivery, and an ‘out-of-context’ control at Δ*τ* = +1200 s (+20 min) that displaces the target far outside its reward window. For all FR-based runs, a right-asymmetric (‘causal’) VQT kernel is used (see Section 5.14.2), ensuring each temporal factor depends only on past information.

Cross-validation uses a group-shuffle-split protocol: each session is divided into contiguous, non-overlapping ∼ 10 min blocks randomly assigned to training (70%) and test (30%) partitions. Classifier performance is summarized by the area under the receiver-operating-characteristic curve (AUC-ROC), the area under the precision-recall curve (AUCPR), and the maximum *F*_1_ score; see Section 5.6.5 for details.

#### 5.6.2 FR-RNN

The FR-RNN receives rank-R temporal factors **C** ∈ ℝ^*T* ×*R*^ from the FR pipeline. The architecture is a single-layer LSTM with hidden size *h* = 5 and input dimensionality *R* = 5, trained on ∼ 5 s batches using the Adam optimizer with cross-entropy loss. Regularization is not used given the small network size. The model is implemented in PyTorch.

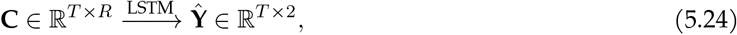

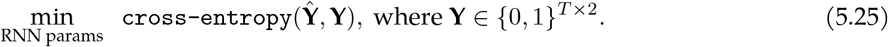

#### 5.6.3 Linear decoders (FR-linear and Points-linear)

Both linear models are L2-regularized logistic regression classifiers with class weights inversely proportional to class frequency, implemented in scikit-learn.

##### FR-linear

Receives the same rank-R temporal factors as the FR-RNN, min-max scaled to [0, 1] and mean-centered. `LogisticRegressionCV` with the L-BFGS solver is used; regularization strength C is selected via 5-fold cross-validation over ∼30 logarithmically spaced values in [10^−6^, 10^3^], optimizing AUC.

##### Points-linear

Operates on raw tracked-point positions (*P* ≈ 930 keypoints × 2 coordinates, yielding **X** ∈ ℝ^*T*^*′*^×2*P*^ at 120 Hz), each column mean-centered. `LogisticRegression` with the SAGA solver is used; *C* is tuned via Bayesian optimization (Optuna; up to 20 trials over [10^−6^, 10^3^]).

#### 5.6.4 Points-RNN

Operates on the same raw tracked-point positions as Points-linear, reduced via per-session PCA to *K* = 100 components (computed on the full session prior to train/test splitting, analogous to how nnTCA is pre-computed for FR models). The model uses the same single-layer LSTM architecture as the FR-RNN (h = 5), with input dimensionality *K* = 100. Because reward events are rare ( ∼ 1.1% of frames), the loss is weighted binary cross-entropy via PyTorch’s `BCEWithLogitsLoss` (pos_weight = n_neg_/n_pos_), trained with Adam (lr = 10^−3^) on random chunks of ∼ 5.3 s (636 frames at 120 Hz). Early stopping terminates training when smoothed validation loss improvement falls below 10^−4^ over a 30-check window, or after 1500 epochs; the checkpoint with highest test-set AUC is retained. At test time, inference is chunked, carrying the LSTM hidden state across chunks. Within each session, 10 independent repeats are run; the median AUC across repeats is computed per session, averaged per mouse, and summarized as mean ± s.e.m. (*N* = 4).

#### 5.6.5 Precision-recall evaluation of reward-time decoders

We sweep a threshold *θ* over the model’s output probabilities 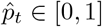 using held-out test data and compute receiver operating characteristic (ROC) and precision-recall (PR) curves relative to the binary reward-event target *y*_*t*_ ∈ {0, 1} . To pool across heterogeneous test sets, curves are linearly interpolated onto common FPR and recall grids (200 points on [0, 1]), then averaged hierarchically (sessions within mouse, then across mice). Chance performance is the identity line for ROC and the positive-class rate *π* for PR.

The AUCPR metric is computed directly from probabilities and labels via `sklearn.metrics.average_precision_score`, which uses the step-function estimator.

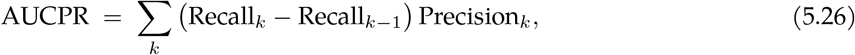

(this avoids the linear-interpolation bias of the trapezoidal rule on PR curves). The maximum *F*_1_ score is the best achievable *F*_1_ over all classifier thresholds, taken on the same PR curve,

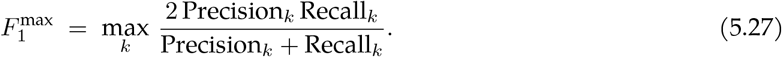

At prevalence *π* the chance baselines are AUCPR_chance_ ≈ *π* and 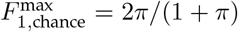. Values are averaged across cross-validation repeats and sessions.

To test whether in-context decoding exceeds the out-of-context (long-shifted control) baseline, for each (model, lag) condition, the per-mouse metric value m_*i*_ was paired with the same mouse’s per-mouse value *c*_*i*_ at Δ*τ* = +1200 s, and the paired differences *d*_*i*_ = *m*_*i*_ − *c*_*i*_ were tested for E[d] > 0 using a one-sided paired *t-*test. Effect size is reported as paired Cohen’s 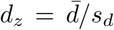. Multiple comparisons are corrected using the Benjamini-Hochberg FDR procedure.

#### 5.6.6 Decoder COEFFICIENT interpretability

To visualize and compare the coefficient structure of the two linear decoders (**Figure 4 supplement 2**), FR-linear and Points-linear were re-fit on single representative sessions using the same fitting procedure as above. Here FR-linear receives the 10 FR temporal factors; Points-linear receives the raw point displacements.

Model-free summaries in **Figure 4 supplement 2Bii-Biv** are: per-point mean reward-triggered displacement (mean absolute per-point displacement in a window around reward events minus the baseline-window mean); per-point SNR (ratio of reward-window mean to baseline-window mean); and per-point signed point-biserial correlation (`scipy.stats.pointbiserialr` between the per-frame reward label and each point’s per-frame displacement, signed by whichever axis has the larger absolute correlation).

### 5.7 Convolutional reduced-rank regression

Here, we describe a custom model that is used to predict neural activity from raw point-tracking data. This model combines convolution with reduced-rank regression (RRR) to learn a mapping from raw facial movements to neural activity. The overall structure of the model is related to a small linear convolutional neural network; however, the convolutional layer is implemented differently and is described below.

#### 5.7.1 Model architecture

Let **X** ∈ ℝ^2*P* ×*T*^ be the behavioral data matrix (raw displacements of points on the face) and **Y** ∈ ℝ^*N* ×*T*^ the deconvolved neural activity matrix, where 2*P* is twice the number of points, *N* is the number of neurons, and *T* is the number of time samples. For a temporal window *τ* = 0, …, *L* − 1 that describes the index of the convolutional kernel or the temporal lag between the kernel and the signal, we define a rank-R convolutional reduced-rank regression model in three stages.

i. **Encoding** Project the raw face point displacement signals onto *R* spatial weight vectors. For each rank in **W**_enc_ ∈ ℝ^2*P* ×*R*^ we can calculate an encoding vector **e**_*r*_:

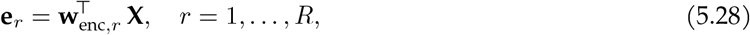

resulting in the intermediate matrix **E** ∈ ℝ^*R*×*T*^.
ii. **Convolution** The *R* encoded channels are convolved with a set of *R* convolution kernels (Fourier convolution is used in code, see Section 5.12 for details). The convolution operation proceeds in one of three ways, depending on the model type:
  - (ii.a) Basic convolution

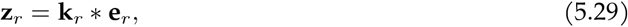

where **k**_*r*_ ∈ ℝ^*L*^ is the convolution kernel for component r.
  - (ii.b) Spectral convolution

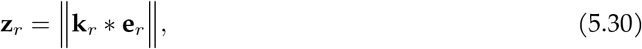

where **k**_*r*_ ∈ ℂ^*L*^ is the convolution kernel for component r, and optimization independently learns the real and imaginary parts of the kernel.
  - (ii.c) Hilbert-spectral convolution

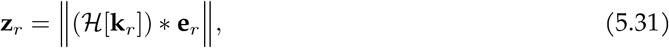

where **k**_*r*_ ∈ ℝ^*L*^ is the convolution kernel for component r, and ℋ[·] is the Hilbert transform operator.
iii. **Decoding** Linearly combine convolved channels:

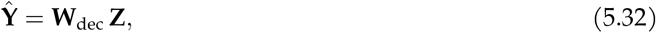

##### Unified form

We combine the encoding and decoding steps that make up the RRR with the single notation summarizing the three versions of the convolution step using the following equation:

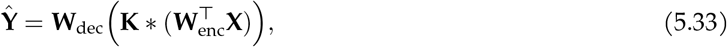

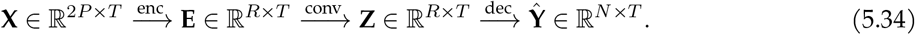

where **W**_enc_ ∈ ℝ^2*P* ×*R*^, ***K*** ∈ ℝ^*R*×*L*^ or ℂ^*R*×*L*^, **W**_dec_ ∈ ℝ^*N* ×*R*^, and the result **Ŷ** ∈ ℝ^*N* ×*T*^ . The convolution operator [∗] represents one of the three convolution types described above and is performed on each rank independently.

#### 5.7.2 Optimization

The model is trained to minimize the mean squared error (MSE) between the predicted neural activity **Ŷ** and the true neural activity **Y**:

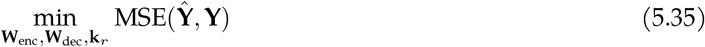

All parameters {**W**_enc_, **W**_dec_, **K}** are optimized jointly with ridge regularization using either the Adam or L-BFGS optimizer (using PyTorch’s built-in implementations). Hyperparameter optimization is performed using cross-validation and the `optuna` library over the ridge regularization strength. Model performance evaluation is performed on a fully held-out test set that is not used for training or hyperparameter optimization.

#### 5.7.3 Model performance

We calculate model performance as the held-out explained variance ratio (EVR),

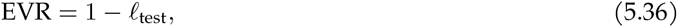

where *ℓ*_test_ is the test-set mean-squared error between the predicted and true neural activity, evaluated after normalizing each neural dimension by its training-set standard deviation. Because this metric is not rescaled to the predicted signal, it can take small negative values on poorly predicted held-out data; per-mouse EVR values are clamped at zero before averaging across animals.

Statistical comparisons between model types (**Figure 5F**) use the two-sided paired Student’s *t-*test, across *N* = 5 mice.

#### 5.7.4 Band-pass filtering

To examine frequency-specific performance, we additionally rerun analyses using band-pass filtered behavioral input signals. A set of 10–20 band-pass filters is designed with center frequencies logarithmically spaced between 0.01–40 Hz, a Q-factor of 5, and Hamming windowing. FIR filtering is used to apply these filters (see Section 5.13). The filtered behavioral signal is then used in the same model pipeline as above to compute the band-specific EVR. Statistical comparisons between model types at each frequency band are corrected for multiple comparisons using the Bonferroni method across bands.

#### 5.7.5 Comparison with motion-energy PCA

To test whether point-tracking discards facial movements that a denser description would retain for predicting M1 population neural activity, we repeated the convolutional reduced-rank regression analysis using a pixelwise motion-energy predictor, holding the neural target **Y**, the cross-validation, the three model arms, the temporal window, and the performance metric fixed (Section 5.7).

For each mouse, per-frame motion energy was computed as the absolute grayscale frame difference, |*g*_*t*_ −*g*_*t* −1_, within the face region covered by the tracked points. Multiple sessions from the same mouse were rigidly registered onto a common pixel grid, so that a single spatial basis could be used across sessions. We then reduced the dimensionality of the dense pixelwise signal into a rank-200 time series using singular value decomposition.

### 5.8 Single-neuron coupling to face-rhythm components

We quantify how individual M1 neurons relate to rhythmic facial-movement components in two complementary ways: (i) phase locking (phase-variant) and (ii) envelope correlation (phase-invariant). For this analysis, we use rank-R = 12 FR decompositions per mouse across *N* = 5 mice (yielding 60 total behavioral components).

#### 5.8.1 Phase-coupling analysis

For each FR component *r* we generate a matched pair of time series, starting with the raw point-tracking displacement matrix **X** ∈ ℝ^2*P* ×*T*^ and the deconvolved neural activity matrix **Y** ∈ ℝ^*N* ×*T*^, where *N* is the number of neurons and T is the number of time samples.

We generate phase-variant behavior signals by projecting the **X** matrix onto individual FR spatial factors **a**_*r*_ ∈ ℝ^2*P*^, and then spectrally filtering the result by treating the corresponding spectral factor **b**_*r*_ ∈ ℝ^*F*^ as a finite-impulse response (FIR) filter kernel (see Section 5.13 for details on FIR filtering):

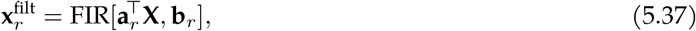

Applying a Hilbert transform over the time dimension yields the analytic signal 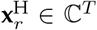 taking its angle gives the instantaneous phase of the behavioral signal:

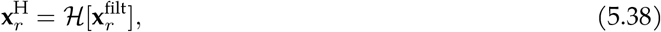

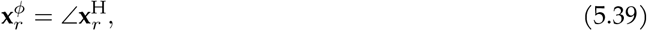

where ℋ[·] is the Hilbert transform operator, ∠ is the complex angle/phase operator, and 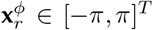 is the phase of the filtered behavioral signal.

A similar procedure is applied to the deconvolved neural activity **Y**. The same spectral filter **b**_*r*_ is used to match the frequency content of the neural activity to that of the behavioral signal. Combining the above 3 steps, we obtain the phase-variant coupling signal for neuron *n* and component r:

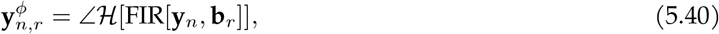

##### Pairwise phase consistency (PPC)

For neuron *n* and component r, we define the phase difference between the neural activity and the behavioral signal as:

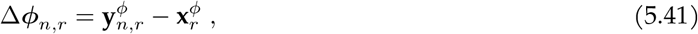

We can then measure the phase-variant coupling using a simplified implementation of the `pairwise phase consistency` (PPC) metric (Vinck, Van Wingerden, et al. 2010; Vinck, Battaglia, et al. 2012):

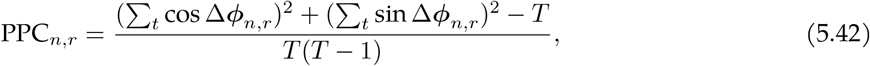

We assessed the statistical significance of each observed PPC value using a parametric z-score approach: PPC was computed repeatedly on 60 randomly circularly time-shifted behavioral signals to estimate the null mean *µ*_null_ and standard deviation *σ*_null_, and a z-score for the observed PPC value was calculated as *z*_*n,r*_ = (PPC_*n,r*_ − *µ*_null_)/*σ*_null_. Two-sided p-values were obtained from the standard normal CDF. These p-values are corrected for multiple comparisons across the *N* neurons tested using the Bonferroni method (*N* comparisons per component).

#### 5.8.2 Envelope correlation

To obtain a phase-invariant behavioral signal, we take the magnitude of the analytic signal of the filtered behavioral signal (from Eq. 5.38), commonly referred to as computing the spectral envelope:

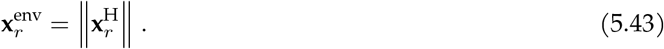

Phase-invariant coupling is measured with Pearson’s correlation (*ρ*) between the filtered behavioral envelope and the unfiltered neural activity:

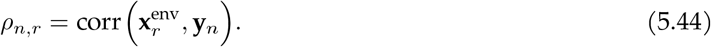

Significance is determined using the same parametric z-score procedure as for PPC, with Bonferroni correction applied across the *N* neurons (*N* comparisons per component).

### 5.9 Independent factorization of neural and facial activity

Here, we describe the approach used to model the joint space of facial movements during head-restrained free locomotion and simultaneously recorded neural activity.

#### 5.9.1 Data acquisition and preprocessing

Data were acquired from the baseline recording sessions of the BMI experiment described in Section 5.11 and Section 5.11.5. This includes widefield facial video and two-photon calcium imaging of layer 2/3 neurons in the primary motor cortex (M1) of head-fixed mice during free locomotion and sparse reward consumption.

Facial videos are processed through the first two steps of the face-rhythm pipeline (Section 5.1) to produce spectrogram tensors ***S*** ∈ ℝ^2*P* ×*F* ×*T*^ . Calcium imaging data are processed to produce a deconvolved neural activity matrix **Y** ∈ ℝ^*N* ×*T*^ . Spectrogram tensors are linearly interpolated to synchronize with the neural time series.

#### 5.9.2 Decomposition of temporal factors

We note that non-negative CP decomposition (as used in the standard FR pipeline) is equivalent to non-negative matrix factorization (NMF) when the input data are two-dimensional (i.e., a matrix).

We applied non-negative CP decomposition (using nn-CP-HALS, see Section 5.1.4) to both the neural activity and spectrogram tensors independently, producing temporal factors for each modality. The rank *R* was set to 10 for both decompositions.

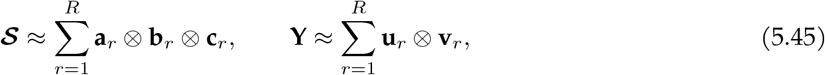

where **a**_*r*_ ∈ ℝ^2*P*^, **b**_*r*_ ∈ ℝ^*F*^, **c**_*r*_ ∈ ℝ^*T*^ are the spatial, spectral, and temporal factors of the behavioral spectrogram tensor; and **u**_*r*_ ∈ ℝ^*N*^, **v**_*r*_ ∈ ℝ^*T*^ are the neural and temporal factors, respectively.

#### 5.9.3 Matching temporal factors

The temporal factors of each modality **V** ∈ ℝ^*T* ×*R*^ and **C** ∈ ℝ^*T* ×*R*^ are then compared by finding a one-to-one pairing that maximizes the summed Pearson correlation.

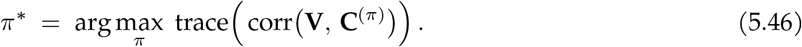

where **C**^(*π*)^ denotes the column-permutation of **C** by *π*. The Hungarian algorithm (implemented using the `scipy.optimize.linear_sum_assignment` function) is used to find *π*^∗^, producing the optimally matched columns of **V** and **C**.

Pearson’s correlation is calculated between the matched temporal factors, and significance is assessed via a parametric z-score approach: the null mean and standard deviation are estimated from n_shuf_ = 25,000 circularly time-shifted controls, and one-sided p-values are obtained from the standard normal CDF. Bonferroni correction is applied across all matched pairs pooled across mice.

### 5.10 Joint factorization of neural-behavioral coherence

Here, we describe the approach used to decompose the joint space of facial movement and neural activity during the performance of the BMI task (see Section 5.11 for details on the experimental setup).

#### 5.10.1 Data acquisition and preprocessing

Data are acquired from the experimental sessions described in Section 5.11. This includes widefield facial video and two-photon calcium imaging of layer 2/3 neurons in the primary motor cortex (M1) of head-fixed mice.

Data are synchronized and peri-reward epochs are extracted from both datasets. The peri-reward epochs are defined as a [ − 5 s, 0 s] window relative to the reward onset time. The facial video is processed through the face-rhythm pipeline (Section 5.1) to produce the peri-reward point-tracking tensor ***X*** ^(*M* )^ ∈ ℝ^2*P* ×*T* ×*M*^ . Neural activity data are processed to produce a similar tensor ***Y*** ∈ ℝ^*N* ×*T* ×*M*^, where *N* is the number of neurons, *T* is the number of time samples, and *M* is the number of trials.

We compute the coherence between the point-tracking signals of all the facial points and the neural activity of all neurons.

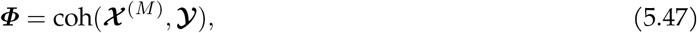

where coh(· ) is the coherence function, which is computed along the temporal dimension, producing ***Φ*** ∈ ℝ^2*P* ×*N* ×*F* ×*M*^ .

We then factorize the coherence tensor using non-negative CP decomposition (mini-batch nn-CP-HALS, see Section 5.1.4) to obtain joint factors:

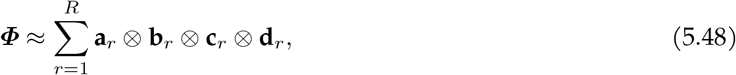

where **a**_*r*_ ∈ ℝ^2*P*^ is the spatial factor for facial points, **b**_*r*_ ∈ ℝ^*N*^ is the neural factor, **c**_*r*_ ∈ ℝ^*F*^ is the frequency factor, and **d**_*r*_ ∈ ℝ^*M*^ is the trial factor. The rank *R* is set to 6 to approximately match the number of factors extracted in the baseline of the BMI experiment (see Section 5.11.6 for details).

#### 5.10.2 Component selection and comparison to BMI decoder

We compare the neural loading that each animal is tasked with activating during the BMI experiment **w**_dec_ ∈ ℝ^*N*^ (the same weighting for all sessions; see Section 5.11.5 and Section 5.11.6 for details) against the neural factors extracted using the joint neural-behavioral factorization. Pearson’s correlation is calculated between each ‘joint neural’ factor 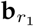 in 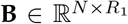 and each factor extracted during the baseline/day 0 recording sessions of the BMI experiment (‘BMI baseline factors’) 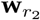 in 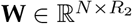, which includes **w**_dec_. We define one of the joint neural factors as associated with the animal’s ‘strategy’ **b**_strat_ by looking at the trial factors **D** ∈ ℝ^*M* ×*R*^ and selecting the component r_strat_ with the largest proportional increase in magnitude over all trials (concatenated across all sessions). We find that in 3 out of 4 mice, **w**_dec_ is most strongly correlated with **b**_strat_, and in the fourth mouse it is the second-strongest match. We assess the significance of this correspondence against a null in which, within each mouse, each of the R_1_ = 6 joint components is equally likely to be the top match to **w**_dec_. Observing **b**_strat_ as the best match in 3 of 4 mice and the second-best in the remaining mouse yields p ≈ 0.004.

### 5.11 BMI experiment data

#### 5.11.1 Animals and surgery

6-week-old male C57BL/6J mice underwent surgery and were group housed in a reverse light/dark cycle. Craniotomies and intracranial injections were performed under isoflurane anesthesia (5% for induction, 0.5– 1.5% for maintenance). Mice were implanted with a titanium headplate and a glass window over the right motor cortex (M1) for two-photon imaging. Stereotactic coordinates were (AP +1.6 mm, ML − 1.9 mm) relative to bregma. The headplate was secured to the skull using dental cement, and the glass window was glued to the skull using cyanoacrylate glue. AAV injections of AAV9-hsyn-jGCaMP8m-WPRE or PHP.eB-synRibo-L1-jGCaMP8s were made using a beveled pulled glass micropipette needle; 6–14 injections at 200 nl each were made near the center of the craniotomy based on vascular anatomy with an infusion rate of 50– 75 nl per minute. Virus was diluted to 1.5 × 10^12^ vg/mL. Mice were allowed to recover for at least 10 days before any experimental procedures were performed. Buprenorphine, ketoprofen, and saline were given for post-surgical analgesia and recovery. After the recovery period, mice were partially water restricted to 85–90% body weight and given 1.0–1.5 mL of water for maintenance. All procedures were approved by the Institutional Animal Care and Use Committee (IACUC) at Harvard Medical School.

#### 5.11.2 Data acquisition

Mice are head-fixed, placed on a linear treadmill, and allowed to run freely. Widefield video of the face is acquired at 120 Hz, 1 ms exposure time, and MPEG-2 compression, using a FLIR Flea3-13Y3M-C camera illuminated with infrared light. Videos of calcium activity in layer 2/3 neurons of M1 are acquired at 30 Hz. A custom-built resonant-galvo two-photon microscope is controlled using ScanImage. A 16 × 0.8-NA Nikon (N16XLWD-PF) objective is used, the excitation laser wavelength is 960 nm from a Ti:sapphire laser (Coherent Chameleon Ultra), the power is set to at most 70 mW, and the resolution is set to 512 × 1024 pixels. Data are also collected from a treadmill velocity sensor and a lick spout sensor. A synchronization pulse is sent to and recorded by each camera, and is used to align all data streams.

#### 5.11.3 Neural data preprocessing

Neural data are preprocessed using the Suite2p (Pachitariu et al. 2016) and ROICaT pipelines. For each region-of-interest (ROI), Suite2p produces: an ROI mask, neuropil mask, ROI fluorescence time series, neuropil time series, and deconvolved calcium event time series. For some analyses, the deconvolved calcium event time series is used, while for others the dF/F is computed as:

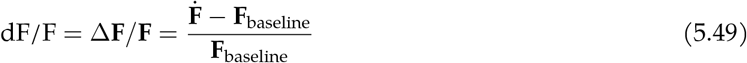

where 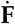 is the neuropil-corrected fluorescence time series computed as:

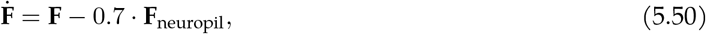

and **F**_baseline_ is the baseline fluorescence time series computed as the rolling 30th percentile of the fluorescence time series over a 10-minute window.

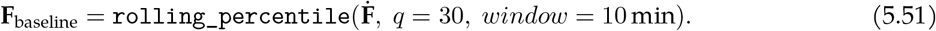

ROI classification is performed using the ROICaT library. Image embeddings are computed for each ROI mask, visualized in a 2D space using UMAP (McInnes, Healy, and Melville 2020), and a region of the embedding projection is selected to include all ROIs that look like neurons. ROIs that look like dendrites, axons, non-neuronal structures, or out-of-focal-plane neurons are excluded. Subsequent quality inclusion criteria are applied to each ROI based on their dF/F time series. These include: (i) minimum ROI to neuropil variance ratio, (ii) maximum explained variance of ROI by neuropil, (iii) minimum baseline ROI fluorescence, (iv) minimum signal-to-noise ratio (SNR), (v) maximum dF/F value, and (vi) maximum variance of baseline fluorescence time series. Only ROIs that pass all inclusion criteria and are also classified as neurons are included in subsequent analyses.

#### 5.11.4 BMI design

Mice are trained to control their neural activity to achieve rewards in a closed-loop brain-machine interface (BMI) experiment. The data gathered during this experiment fall into two categories: (i) baseline recordings of natural behavior prior to training, and (ii) BMI training sessions in which mice are provided with real-time feedback of their neural activity and are rewarded for achieving a target.

During BMI training sessions, the neural activity of the animal is processed, decoded, and converted into a controllable auditory and visual cursor that is played back to the animal in real-time to allow the animal to learn to control the cursor’s dynamics (see Section 5.11.6 and Section 5.11.7 for details on decoder design and Section 5.11.8 on the decoder forward algorithm). The input to the decoder is the instantaneous activity of all neurons being monitored in the field-of-view (FOV). The output, referred to as the ‘cursor’, is a bounded scalar value c ∈ [0, 1] representing the animal’s instantaneous performance. The cursor value is converted into a tone frequency that is played back to the animal, as well as the position of an illuminated LED on a linear LED strip. If the cursor value is above a certain threshold, the animal is also rewarded with a drop of soy milk. The tone frequency is logarithmically mapped to the cursor value between 2 kHz and 18 kHz, with the reward threshold at 18 kHz. The blue LED strip is oriented from left to right across the mouse’s visual field, and ∼ 1 light on the LED strip is illuminated at a time; the position is linearly mapped to the cursor value with the reward threshold at the right-most position.

- **Auditory feedback:** A sine wave / pure tone is generated on each frame with tone frequency *f*_tone_(t):

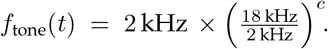
- **Visual feedback:** A pixel on a linear LED strip containing *L*_LED_ = 72 contiguous segments is illuminated on each frame, where the illuminated segment index *i*_LED_(t) is computed as:

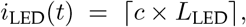

which maps left-to-right across segments 1–72.

#### 5.11.5 Baseline session recordings

##### Baseline session

Baseline recordings are collected for 1–4 days / sessions prior to BMI training. During these 1-hour sessions, mice are played back the auditory and visual cursor feedback from a randomly selected previous BMI session of a trained animal. Sparse soy milk rewards are given by setting the cursor threshold value so that they receive ∼ 0.5 rewards per minute. This playback is fully open-loop and independent of the animal’s behavior or neural activity. Neural data from these days are preprocessed using Suite2p; region-of-interest (ROI) classification and cross-session tracking are performed using ROICaT, and custom pipelines for quality control are used to obtain the final dF/F or deconvolved calcium event time series of high-quality neurons tracked across the baseline sessions. The number of neurons tracked that passed the inclusion criteria is roughly 200–800 neurons per session. For some analyses, single-day recordings are used without tracking, with the number of included neurons being roughly 700–1500 per session.

##### Reference image acquisition

A three-dimensional reference image is acquired on the first day of the baseline sessions to enable precise spatial alignment across all experimental days. A dense z-stack is acquired with 41 optical slices at a 0.8 *µ*m step size, spanning a total axial range of 32 *µ*m centered around the chosen focal plane. Each slice is averaged across 10 acquisition iterations with 60 frames each to create a high-quality reference image for each z-position. The final z-stack that is used for online motion correction is a subset of the dense z-stack, consisting of 5 evenly spaced slices at 4 *µ*m intervals for computational efficiency during online motion correction.

#### 5.11.6 Decoder axis selection process

We use the neural activity recorded from the baseline sessions to define a set of up to 6 neural factors 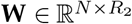. One of these factors **w**_dec_ is used as the ‘decoder axis’ for the real-time decoder, which maps the neural activity to the instantaneous cursor value.

##### Pre-orthogonalization

To reduce the influence of task-irrelevant factors on the selected decoder axis, we orthogonalized out signals like slow temporal trends, motion artifacts, structured noise, and population-averaged neural activity from the baseline neural activity matrix **X** prior to any factorization. These included:

1. Average neural activity
2. Average neuropil activity
3. Slow temporal trends
4. Session indicator variables (for concatenated baseline sessions)

The average neural (1) and neuropil (2) activity is split into band-passed components using a set of FIR band-pass filters with boundaries at [0, 0.003, 0.05, 0.5, 5, ∞ ] Hz, and Kaiser windowing with *β* = 3.0. The slow temporal trends (3) are modeled using a linear time trend and a nonlinear M-spline basis function over the duration of the temporal dimension with order 3 and 4 basis functions. The session indicator variables (4) are used to remove any spurious ‘jumps’ in offsets introduced by concatenating multiple sessions of neural activity. These variables are placed into a design matrix **D** ∈ ℝ^*T* ×*Z*^, where *Z* is the total number of featurized time series variables.

Orthogonalization is performed by subtracting off the projections of the design matrix from the neural activity matrix **X** ∈ ℝ^*T* ×*N*^ (transposed from other sections), where *N* is the number of neurons and *T* is the number of time samples. First, to improve stability, PCA is performed on the design matrix:

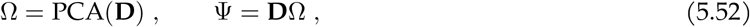

where Ω ∈ ℝ^*Z*×*R*^ are principal components (loadings) and Ψ ∈ ℝ^*T* ×*R*^ are scores. Then, we compute the optimal projection of the design matrix onto the neural activity matrix using ordinary least squares (OLS):

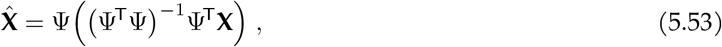

where 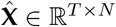 is the reconstructed neural activity matrix based on the design matrix. The orthogonalized neural activity matrix is then computed as:

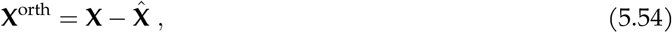

where **X**^orth^ is the orthogonalized neural activity matrix. For simplicity, the above three steps can be notated as:

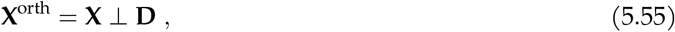

##### SPCA

Following pre-orthogonalization, a low-dimensional neural subspace 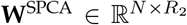 is generated using sparse-PCA (sklearn’s `sklearn.decomposition.SparsePCA` implementation). We used a grid search over the sparsity hyperparameter (L1 penalty) *α* such that the number of non-zero neuron loadings is approximately 100, averaged across factors. The rank *R* is set to 6, and any factors with less than 30 non-zero neuron loadings are discarded. A small L2 penalty is also used to ensure the stability of the optimization. The resulting factors are ordered by their variance explained in the pre-orthogonalized neural data. Sparse PCA was selected because it discovers factors that are aligned to the latent distributional modes better than standard PCA.

##### Decoder weight fine-tuning

To define the final factor space **W**, we then process the SPCA factors with the following steps:

- **Reprojection:** The SPCA factors **W**^SPCA^ are reprojected using Ridge regression to regularize and distribute the weight representation across more neurons so as to increase robustness to changes in single-neuron signal-to-noise or tuning. This is done by defining a new weight matrix 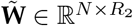 to predict the SPCA scores **XW**^SPCA^ from the pre-orthogonalized neural activity **X**^orth^:

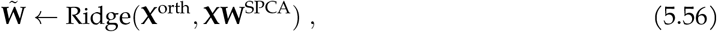
- **Orthogonalization:** The weight matrix 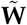 is then orthogonalized to ensure that the factors are orthogonal to each other. This is done using a Gram-Schmidt-like process, where the orthogonalization operator ⊥ is derived from Eq. 5.55

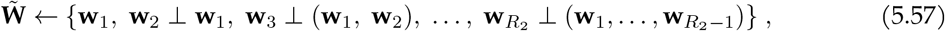

and **w**_*i*_ is the ith factor in 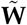.
- **Mean subtraction:** The mean of each factor in 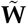 is then subtracted to center the factors around zero (effectively orthogonalizing the factors to the mean of the neural population activity).
- **Sign flipping:** The sign of each factor in 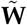 is flipped, if necessary, so that the skewness of the factor distribution is positive. In nearly all cases, no flipping was necessary, as SPCA factors tend to align with the natural positive skewness of the neural activity distribution.

The final decoder weight matrix is then defined as 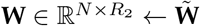.

#### 5.11.7 Neural decoder definition

The neural decoder is designed to incentivize the animal to voluntarily control its neural activity specifically along the decoder axis. The decoder is implemented as:

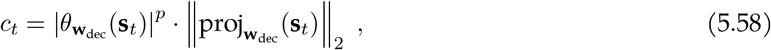

where **w**_dec_ ∈ ℝ^*N*^ is the decoder weight vector (‘decoder axis’), which is selected as a column from the ***W*** matrix. **s**_*t*_ ∈ ℝ^*N*^ is the instantaneous neural activity vector (see Section 5.11.9). 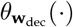 is the cosine similarity between **w**_dec_ and another vector, and 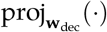 is the projection operator onto **w**_dec_. Lastly, a scalar power coefficient p is used to control the angular selectivity of the decoder.

An offline simulation of BMI performance is conducted on the selected factor using the baseline recording dataset to validate the decoder’s performance and tune any task-related parameters. We typically select the factor with the second highest explained variance, dependent on the simulation being able to achieve a satisfactory number of rewards. We avoid choosing **w**_1_, the first factor / principal component, as a target factor, as a previous report suggests that the first factor is significantly related to a “neural engagement” during the trial period (Hennig et al. 2021). The main parameter that is tuned is *θ*_reward_, which defines the cursor reward threshold; this value is tuned so that the simulated reward rate is ∼ 1.25 rewards per minute.

Qualitatively, the cursor output increases linearly with increases along the axis of the selected factor (i.e., the ‘decoder axis’), but falls off sublinearly with angle changes away from the decoder axis. When the power coefficient is zero (*p* = 0), the decoder becomes a simple projection magnitude decoder, and when *p* → ∞, the cursor output is only non-zero when the neural activity is perfectly aligned with the factor direction. For the BMI task, we use *p* = 2 to balance selectivity and linearity.

#### 5.11.8 Task structure

Here, we describe the methods used to control the task’s state-transitions and the delivery of rewards during the BMI task. The task is structured as a state machine with several states: (i) Trial periods, when the sensory feedback is on and the animal is eligible to receive a reward, (ii) reward periods, when the feedback is briefly paused at the current value and a ∼ 3.5 *µ*l soy milk droplet is delivered, (iii) the inter-trial-interval, when feedback is off and rewards are not available, (iv) timeout periods when the animal has not achieved a reward over the maximum trial period duration ( ∼ 20 s), and (v) ‘waiting for quiescence’ periods where the animal must reduce its neural activity along the decoder axis in order to start a new trial period.

##### Algorithm 1

Finite-state controller for the closed-loop BMI task

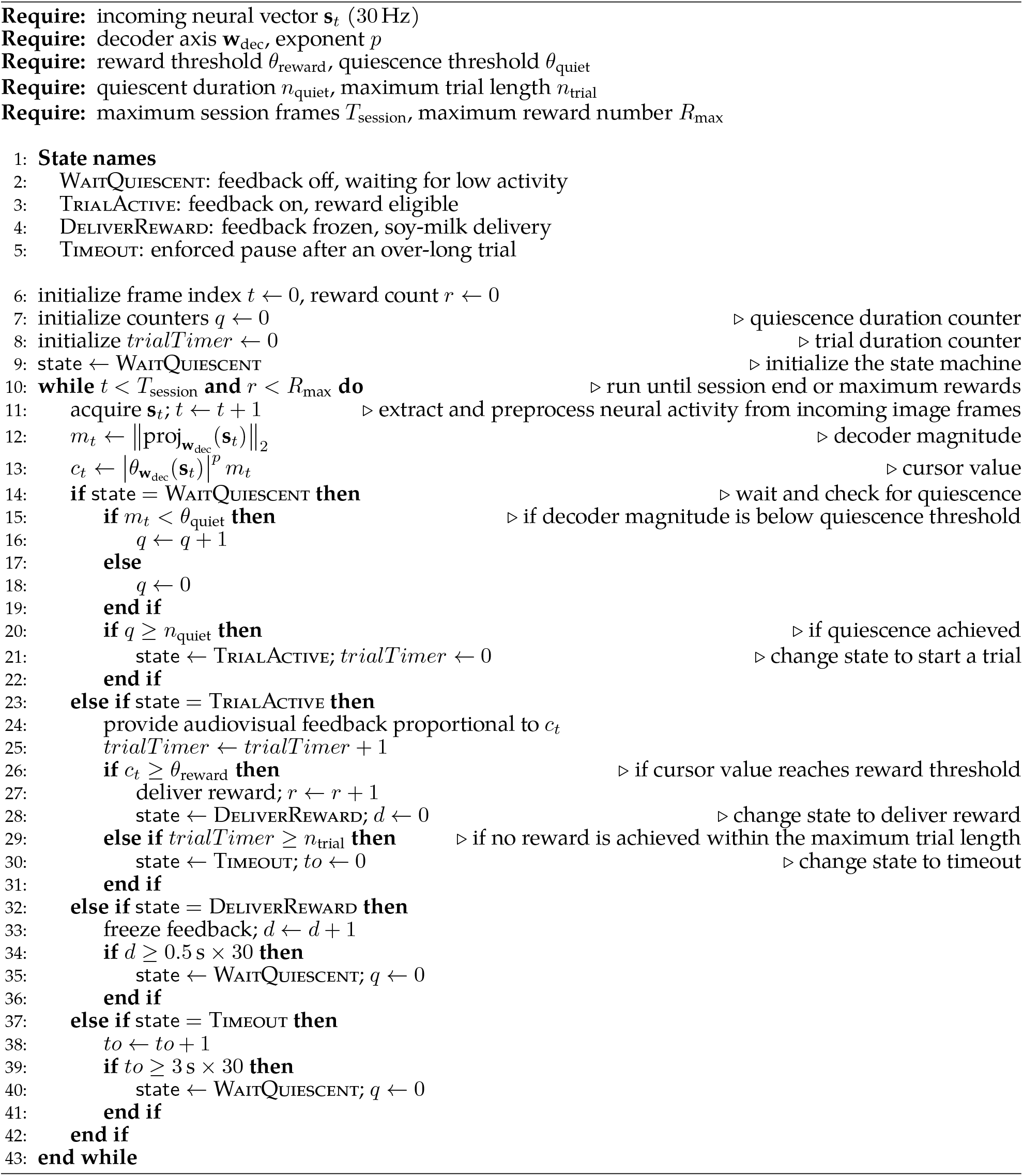

#### 5.11.9 BMI software system

The software suite consists of an offline analysis pipeline written in Python for defining the decoder weights and preparing ROI masks, and a real-time integration and control system written in MATLAB and controlled via a ScanImage user function for online decoding and feedback. Code is available at https://github.com/RichieHakim/BWAIN.

On ‘day 0’ (defined as the day prior to the first BMI session with feedback), the decoder weights are computed, ROI masks are defined, and a simulation is conducted to set the cursor reward threshold. The decoder weights are then used to compute the cursor value in real-time during the BMI sessions.

Closed-loop latency was found to be 10–25 ms, defined as the time from the end of scanning a frame to the output of new feedback signals.

##### Non-rigid image transformation

To compensate for tissue deformation over days/sessions and maintain spatial correspondence of neural decoder masks across sessions, we apply a non-rigid transformation to the ROI masks, neuropil masks, z-stack, and spatial decoder weights on each experiment day. To compute the warping field, we use MAT-LAB’s `imregdemons` function with 500 iterations across 4 pyramid levels and an accumulated field smoothing parameter of 2. The reference image is acquired on ‘day 0’ and each experiment day we acquire a 10-minute-long calibration recording and generate an image to warp to the reference. During this time, the animal is in an idle state without sensory feedback.

On each experiment day, after manually aligning the current field of view to coordinates from the day 0 session, a mean reference image is computed using an average of the most stable frames (identified by minimal motion artifacts). Then, we apply a non-rigid image warp to ensure robust coordinate alignment between sessions.

##### Motion correction

Online motion correction has two components: a fast planar correction to the current imaging frame and a slower axial correction to a rolling average of the imaging plane.

##### Planar correction

Online motion correction during a BMI experiment is implemented using a custom phase-correlation algorithm. To improve robustness, we apply band-pass filtering to the input images using a third-order Butterworth filter with cutoff frequencies at [1/64, 1/4] cycles/pixel (image resolution is cropped from 512 × 1024 to 512 × 512).

##### Axial correction

For online axial (‘z-axis’) drift correction, we use the warped reference z-stack (from day 0) to compare the current image frame in order to determine the current z-slice position. A rolling average of 60 frames is correlated against the warped z-stack, and the estimate of current z-position is determined by maximum phase-correlation. When the estimated current z-position is different from the center plane, we move the imaging plane by 0.5 *µ*m in the direction of the center plane. Axial drift correction is only performed during the inter-trial-interval (ITI) period of the task.

##### Online BMI

During BMI sessions, neural signals are extracted, classified, and used to update the current cursor value in real time.

##### Online neural data preparation

Online neural signal extraction is performed in real-time at a 30 Hz frame acquisition rate. The online extraction process is designed to match, as closely as possible, the extraction process used by Suite2p during offline ROI time series signal extraction. Raw fluorescence signals *F* ∈ ℝ^*N* ×*T*^ are calculated as weighted sums of pixel intensities from each flattened frame **a**_*t*_ in the video **A** ∈ ℝ^*H*·*W* ×*T*^ (where *H* and *W* are the height and width of the image frame) and ROI masks **M** ∈ ℝ^*N* ×*H*·*W*^ :

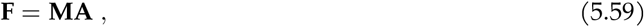

where ***F*** ∈ ℝ^*N* ×*T*^ is the fluorescence time series for *N* ROIs and *T* time samples. The ROI masks are non-rigidly transformed to match the current image frame using the non-rigid transformation field computed during the motion correction step.

A similar process is used to compute the neuropil signal **F**_neuropil_ ∈ ℝ^*N* ×*T*^, using the non-rigidly transformed neuropil masks. The neuropil signal is then subtracted from the ROI signal using a scaling factor of 0.7, matching the offline dF/F calculation (similar to Eq. 5.49).

Baseline fluorescence **F**_baseline_ ∈ ℝ^*N* ×*T*^ is updated every 10 frames using a rolling 30th percentile calculation over a 10-minute sliding window. For online quality inclusion criteria, the minimum baseline ROI fluorescence and the maximum dF/F value are applied with parameters that match the offline process (see Section 5.11.3). Any ROIs that do not meet the inclusion criteria are replaced with NaN and ignored in subsequent calculations.

##### Trial types

BMI training sessions included two types of catch trials, which are sparsely distributed throughout the session. These are not used in any of the analyses in this study. In the first type of catch trial, feedback is decoupled and randomly played back from a different session, and rewards could occur. In the second type, neither the feedback nor the reward is available.

##### Task parameters

Each trial had a maximum duration of 20 s, with minimum inter-trial intervals (ITIs) of 3 s. Any threshold crossing event is discarded if it lasted less than 3 frames. The reward is calibrated to be ∼3.5 *µ*l per delivery.

### 5.12 Fourier convolution

For several analyses, we used Fourier convolution (‘FFTconv’) in place of direct convolution for efficiency and simplicity. We note that the convolution theorem provides the following equality:

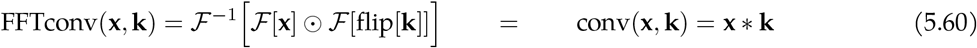

where ℱ [· ] is the Fourier transform operator, ℱ^−1^[· ] is the inverse Fourier transform operator, ⊙ is elementwise multiplication, flip[·] is the flip operator (which reverses the order of the elements along the time dimension). For specific applications, we checked for equivalence with direct convolution to avoid instances of non-negligible ringing and edge artifacts.

### 5.13 Finite-impulse response (FIR) filtering

For several analyses, we used a zero-phase finite-impulse response (FIR) spectral filter. Filtering is applied via forward-backward convolution to prevent phase distortion.

We implemented the spectral filter using Fourier convolution, similar to above (see Section 5.12):

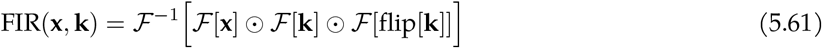

### 5.14 Variable-Q transform (VQT)

The VQT is a time-frequency analysis method that is similar to the short-time Fourier transform (STFT) but with a logarithmic frequency scale and a tunable frequency-dependent bandwidth. The VQT is often used to allow for low-frequency signals to be represented with high temporal-resolution and low frequency-resolution (Q ≈ 4), while high-frequency signals can be represented with low temporal-resolution and high frequency-resolution (Q ≈ 20). The VQT is related to a Morlet wavelet transform, and is similar to the constant-Q transform (CQT) but with a variable Q-factor.

Our implementation computes spectrograms via a convolution between the signal and a set of pre-computed filters, whereas the original VQT implementation uses recursive down-sampling to approximate lower-frequency filters (Brown 1991; Chatterji et al. 2004). Our implementation results in an exact solution without artifacts, more flexibility in the choice of parameters, and the ability to arbitrarily customize filter kernels.

#### 5.14.1 VQT implementation

First, a set of filters/kernels is generated. Let f_*s*_ be the sampling rate, *F* the number of filters, and 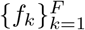 the center frequencies. We specify two endpoints Q_low_ and Q_high_ and interpolate each filter’s quality factor logarithmically:

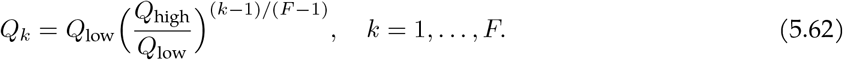

The window length for filter k is

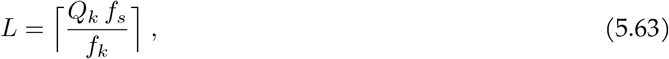

We define the filter kernel / impulse response function **h**_*k*_ of a single filter k as a complex-valued windowed sinusoid:

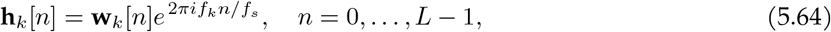

where **w**_*k*_ is a (possibly asymmetric) window function (like Hanning) of length L. The VQT coefficients can be computed by convolution (denoted by [∗]) between the input signal **x** and the filter kernel **h**_*k*_. Optionally, convolution can be implemented using Fourier convolution for efficiency (see Section 5.12 for details):

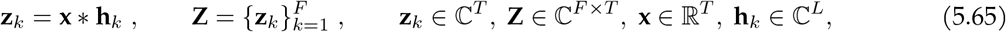

where ***Z*** is the complex spectrogram. Finally, the magnitude spectrogram **S** is obtained via:

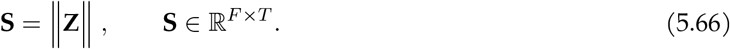

#### 5.14.2 Asymmetric windows

Asymmetric windows can be used to zero out one side/half of the filter. For several analyses, we used right-asymmetric (i.e., ‘causal’) kernels/filter windows, which only encode past information (i.e., ‘trailing signal’) at each time point. Our implementation applies a Heaviside window (on top of any other tapering window functions, like Hanning), and optionally tapers/smooths the step transition to reduce ringing artifacts.

### 5.15 Computational complexity

**Point-tracking** is the dominant computational step in the entire pipeline with a per-frame cost 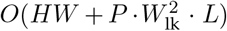 for frame preprocessing (grayscale conversion, CLAHE) and sparse pyramidal optical flow, plus *O*(*PK*) for mesh regularization (where *K* is the number of mesh neighbors). This complexity then scales linearly with total number of frames. Peak memory is determined by the video read buffer, which is user-configurable. In practice, for maximum speed, this value is around 2–20 GB depending on the size of the video and speed of the processor (faster processors benefit from larger buffers to reduce I/O overhead).

**Table 2.** Variables used in computational complexity analysis.

| Symbol | Definition |
| --- | --- |
| $F$ | Number of video frames |
| $H, W$ | Frame height and width (pixels) |
| $P$ | Number of tracked points |
| $K$ | Mesh neighbors per point |
| $W_{\text{lk}}$ | Lucas-Kanade window size |
| $L$ | Lucas-Kanade pyramid levels |
| $B_f$ | VQT frequency bins |
| $D$ | Temporal downsampling factor |
| $T$ | Downsampled time frames ($= F/D$) |
| $R$ | nnTCA rank |
| $I_{\text{max}}$ | nnTCA maximum iterations |

**Spectral analysis (VQT)** applies *B*_*f*_ band-pass filters to each of 2*P* displacement traces via FFT convolution, with total time complexity *O*(*P* · *B*_*f*_ · *F* log *F*) per video.

**Tensor decomposition (nnTCA)** uses Hierarchical Alternating Least Squares (HALS) with per-iteration cost *O*(*R* ·*P* ·*B*_*f*_ ·*T* ), operating on the downsampled spectrogram tensor. Total time is *O*(*I*_max_ ·*R* · *P* ·*B*_*f*_ ·*T* ). Twice the size in memory of the spectrogram tensor is required for this step; in practice, spectrogram tensors are often 1–20 GB in size depending on the user-configured parameters and video length. Therefore, some consideration is required when choosing to use a GPU for this step since GPU memory is often more limited than CPU RAM.

In practice, point-tracking accounts for the majority of wall-clock time. For a one-hour recording on a typical consumer computer, VQT typically completes in well under 10 min on CPU and under 1 min on GPU; nnTCA typically completes in well under an hour on CPU and a few minutes on a GPU. Peak memory usage is typically determined by the nnTCA decomposition.

The throughput benchmarks in **Figure 2 supplement 3** report steady-state processing speed (frames/s) and do not include any one-time preprocessing or warm-up period; this is more reflective of how each method scales with video duration, and for the ∼ 60 min videos used here the warm-up time is small relative to total runtime.

### 5.16 Software dependencies and availability

The face-rhythm pipeline is implemented in Python and depends on standard scientific computing packages including NumPy, SciPy, PyTorch, and OpenCV. GPU acceleration requires an NVIDIA GPU with CUDA support. The complete list of dependencies and version requirements is specified in the package repository. All code repositories developed for this work include open-source licenses:

- face-rhythm: https://github.com/RichieHakim/face-rhythm
- ROICaT: https://github.com/RichieHakim/ROICaT
- Incremental PCA in PyTorch: https://github.com/RichieHakim/incremental_pca
- FastICA implemented in PyTorch: https://github.com/RichieHakim/FastICA_torch
- Variable-Q transform: https://github.com/RichieHakim/vqt
- BMI control software: https://github.com/RichieHakim/BWAIN
- Methods for ablation / benchmarking comparison: https://github.com/RichieHakim/face_rhythm_methods_comparison

### 5.17 Best practices

#### 5.17.1 Video capture and acquisition

Although FR is fairly robust to acquisition conditions, several choices can improve the discovery and signal-to-noise ratio (SNR) of the extracted components. Acquisition frame rates should be at least twice the highest movement frequency of interest (the Nyquist-Shannon sampling criterion (Shannon 1949)); in practice we use 60–250 Hz to capture the full range of rodent facial rhythms (e.g., whisking up to ∼25 Hz, licking ∼5–10 Hz, sniffing ∼3–15 Hz). Spatial resolution should provide ∼10–30 px/mm over the region of interest, ensuring that the Lucas-Kanade integration window ( ∼ 1–3 mm, Section 5.1.2) contains adequate local texture. Global-shutter sensors are preferred over rolling-shutter sensors to avoid intra-frame motion artifacts, and short exposure times (e.g., <4 ms) reduce motion blur.

Heavy video compression, especially codecs with temporal compression (e.g., low-bitrate H.264), can introduce block artifacts that propagate through point-tracking, and thus should be avoided. In our experiments, we use MPEG-2 compression at a high bitrate (e.g., >50 Mbps), though many other settings may be suitable.

Stable, high-contrast illumination is the second major determinant of tracking quality. We illuminate the face with bright and diffuse infrared light (two 50 W, ∼ 850 nm LED chips, invisible to rodents) positioned at a distance to produce shadowing over the face.

#### 5.17.2 Processing hardware

The face-rhythm pipeline is designed to run on consumer-grade hardware. For video processing, a modern multi-core CPU with at least 16 GB of RAM is recommended for the kinds of datasets presented in this work ( ∼ 1500 points, ∼ 1 hr duration, ≥ 600 × 600 pixels). For spectral analysis and tensor decomposition, any CPU is sufficient, but a GPU with at least 8 GB of VRAM is recommended for efficient processing.

### 5.18 Notation guide and style conventions

**Table 3.** Glossary of dimensions.

| Index | Length | Description |
| --- | --- | --- |
| $t$ | $T$ | time bins |
| $p$ | $P$ | points |
| $k$ | $F$ | spectrogram kernels or frequency bins |
| $n$ | $N$ | neurons |
| $r$ | $R$ | rank of CP decomposition or convolutional reduced-rank regression |
| $l$ | $L$ | convolutional kernel index (samples) |

**Table 4.** Glossary of common matrix notation convention.

| Symbol | Number of dimensions | Description | Convention |
| --- | --- | --- | --- |
| $a$ | 0 | scalar | lower-case, plain |
| **$a$** | 1 | vector | lower-case, bold |
| **$A$** | 2 | matrix | upper-case, bold |
| **$\mathcal{A}$** | 3+ | tensor | calligraphic, bold |

**Table 5.**
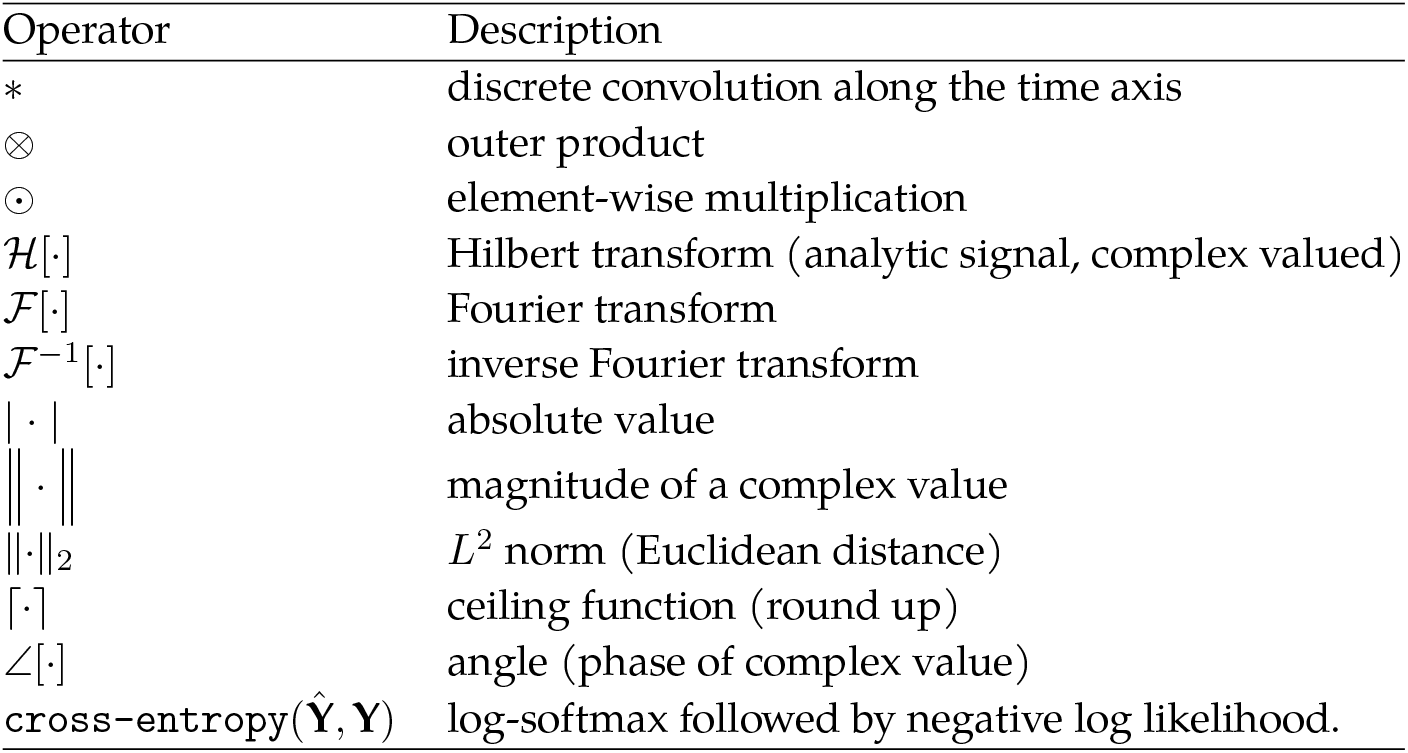
Glossary of operators.

**Table 6.**
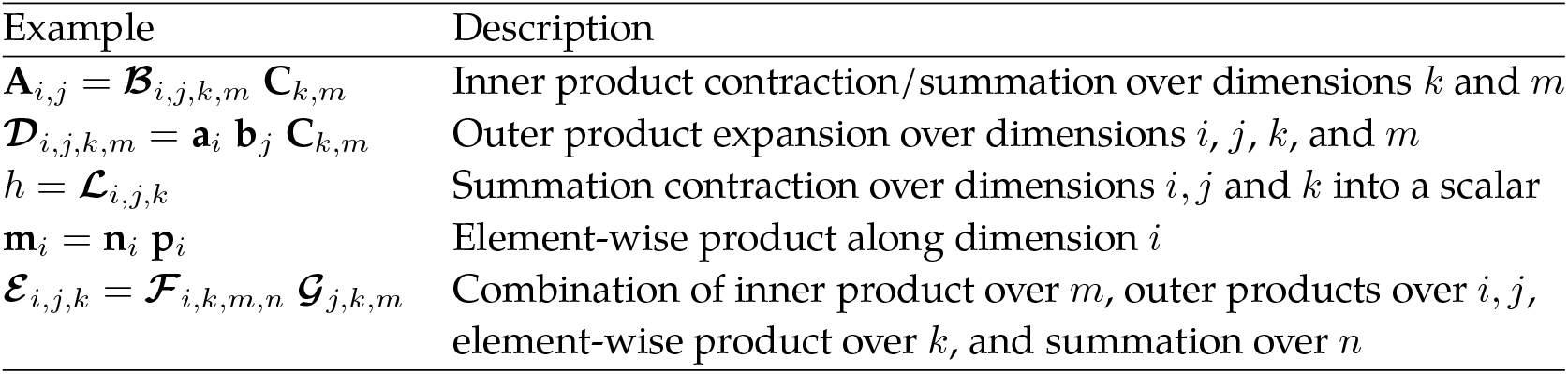
Glossary of Einstein summation notation examples.

| Example | Description |
| --- | --- |
| $\mathbf{A}_{i,j} = \mathcal{B}_{i,j,k,m} \mathbf{C}_{k,m}$ | Inner product contraction/summation over dimensions $k$ and $m$ |
| $\mathcal{D}_{i,j,k,m} = \mathbf{a}_i \mathbf{b}_j \mathbf{C}_{k,m}$ | Outer product expansion over dimensions $i, j, k$, and $m$ |
| $h = \mathcal{L}_{i,j,k}$ | Summation contraction over dimensions $i, j$ and $k$ into a scalar |
| $\mathbf{m}_i = \mathbf{n}_i \mathbf{p}_i$ | Element-wise product along dimension $i$ |
| $\mathcal{E}_{i,j,k} = \mathcal{F}_{i,k,m,n} \mathcal{G}_{j,k,m}$ | Combination of inner product over $m$, outer products over $i, j$, element-wise product over $k$, and summation over $n$ |

## 6 Contributions

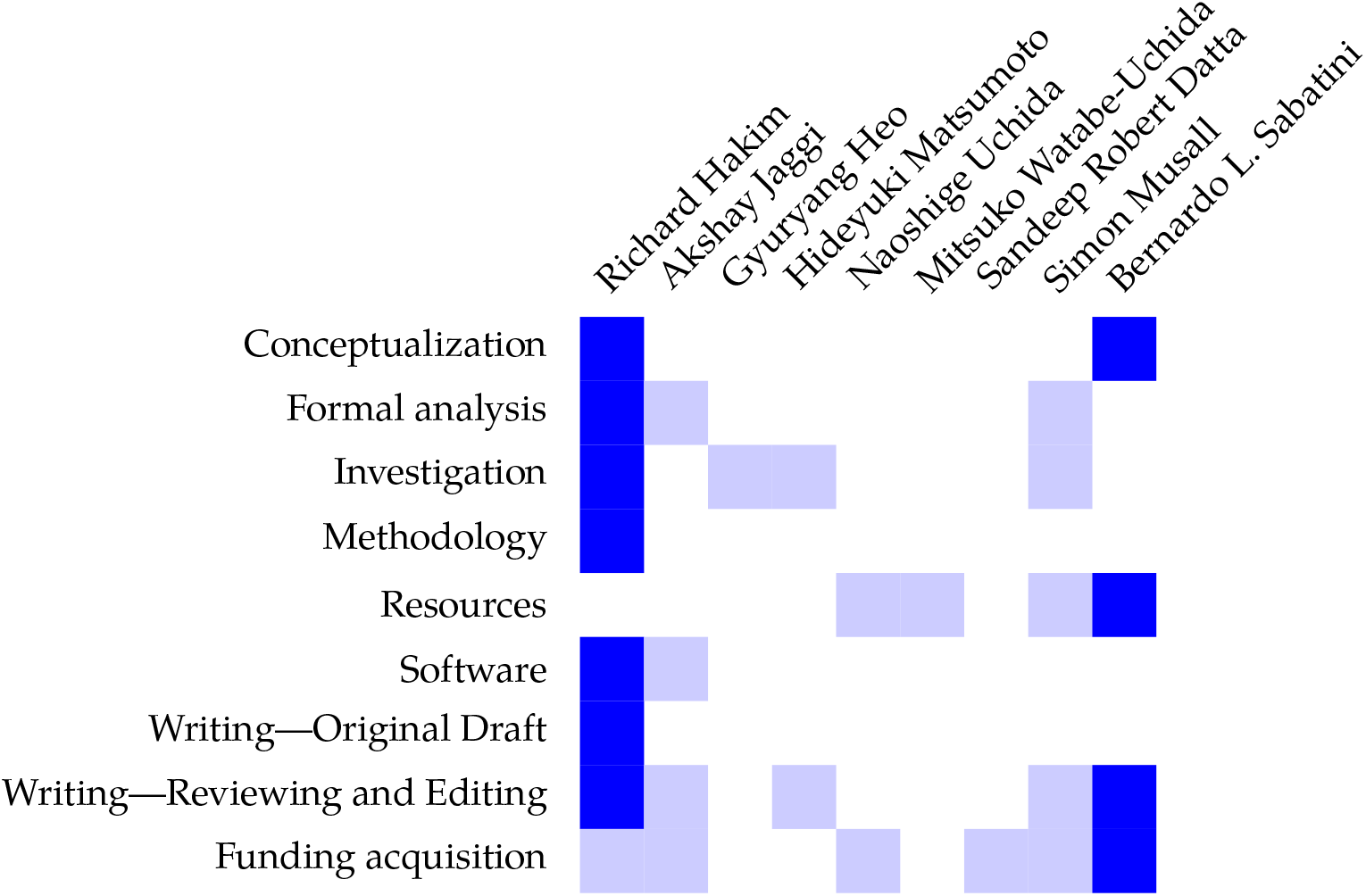

## Acknowledgements

We thank: Alex Williams and Scott Linderman for helpful discussions; and Christopher Harvey, John Assad, and Mark Harnett for guidance.

## 8 Funding attribution

RH is supported by the Kempner Institute for the Study of Natural and Artificial Intelligence at Harvard University, and NSF Graduate Research Fellowship 1000244863.

AJ is supported by award Number T32GM007753 and T32GM144273 from the National Institute of General Medical Sciences. The content is solely the responsibility of the authors and does not necessarily represent the official views of the National Institute of General Medical Sciences or the National Institutes of Health.

BLS is supported by the Howard Hughes Medical Institute, the Kempner Institute for the Study of Natural and Artificial Intelligence at Harvard University, and NIH grants R37NS046579 (Action and interaction of ionotropic and metabotropic neurotransmission), and R35NS137336 (Action and Interaction of Neurotransmitters and Neuromodulators).

BLS, SRD, and NU are supported by NIH grant U19NS113201 (Team DOPE: Towards a unified framework for dopamine signaling and basal ganglia function).

